# Comprehensive mapping of avian influenza polymerase adaptation to the human host

**DOI:** 10.1101/512525

**Authors:** Y.Q. Shirleen Soh, Louise H. Moncla, Rachel Eguia, Trevor Bedford, Jesse D. Bloom

## Abstract

Viruses like influenza are infamous for their ability to adapt to new hosts. Retrospective studies of natural zoonoses and passaging in the lab have identified a modest number of host-adaptive mutations. However, it is unclear if these mutations represent all ways that influenza can adapt to a new host. Here we take a prospective approach to this question by completely mapping amino-acid mutations to the avian influenza virus polymerase protein PB2 that enhance growth in human cells. We identify numerous previously uncharacterized human-adaptive mutations. These mutations cluster on PB2’s surface, highlighting potential interfaces with host factors. Some previously uncharacterized adaptive mutations occur in avian-to-human transmission of H7N9 influenza, showing their importance for natural virus evolution. But other adaptive mutations do not occur in nature because they are inaccessible via single-nucleotide mutations. Overall, our work shows how selection at key molecular surfaces combines with evolutionary accessibility to shape viral host adaptation.

## Introduction

Viruses are exquisitely adapted to interact with host-cell machinery in order to facilitate their replication. Despite significant differences in this machinery across host species, some viruses like influenza can evolve to infect divergent hosts (Parrish et al., 2008; Webster et al., 1992). Such zoonotic transmissions can have severe public health consequences: transmission of influenza virus from birds or pigs to humans has resulted in four pandemics over the last century (Taubenberger and Kash, 2010). These pandemics require the virus to adapt to the new host (Long et al., 2018). Delineating how viruses adapt to new hosts will aid in our ability to understand what determines if a chance zoonotic infection evolves into a human pandemic.

One critical determinant of influenza host range is the viral polymerase (Long et al., 2018), which transcribes and replicates the viral genome (Eisfeld et al., 2014; Te Velthuis and Fodor, 2016). Avian influenza polymerases perform poorly in mammalian cells (Cauldwell et al., 2014; Mänz et al., 2012). This host range restriction likely arises from the need for the viral polymerase to interact with host proteins such as importin-a (Resa-Infante and Gabriel, 2013) and ANP32A (Long et al., 2016), which differ between avian and mammalian hosts. However, it remains unclear exactly how the molecular interfaces between the polymerase and these host proteins are altered during adaptation to humans (Long et al., 2018).

Studies of natural zoonoses and experimental passaging of viruses in the lab have identified a number of mutations that adapt avian influenza polymerases to mammalian hosts (Bussey et al., 2010; Cauldwell et al., 2013, 2014; Chen et al., 2006; Finkelstein et al., 2007; Gabriel et al., 2005; Kim et al., 2010; Mänz et al., 2016; Mehle and Doudna, 2009; Miotto et al., 2008; Mok et al., 2011; Naffakh et al., 2000; Reperant et al., 2012; Taft et al., 2015; Yamada et al., 2010; Zhou et al., 2011). The best known of these mutations is E627K in the PB2 subunit of the polymerase (Subbarao et al., 1993). This mutation alone significantly improves avian influenza polymerase activity in mammalian cells (Long et al., 2013; Mehle and Doudna, 2008), and was considered a key step in adaptation to humans (Taubenberger et al., 2005). But surprisingly, the recent 2009 H1N1 pandemic lineage lacks the E627K mutation. Instead, it has acquired mutations to

PB2 at sites 590 and 591 that similarly confer improved polymerase activity (Mehle and Doudna, 2009; Yamada et al., 2010). This fact underscores the possibility that natural evolution has explored only a small fraction of the possible host-adaptation mutations. Examining only the currently available instances of adaptation in nature or the lab may therefore overlook additional mechanisms of adaptation and evolutionary paths to future zoonoses.

Here, we map all single amino-acid mutations to an avian influenza PB2 protein that enhance growth in human cells versus avian cells. We do so by leveraging deep mutational scanning (Boucher et al., 2014; Fowler and Fields, 2014), which previously has only been used to measure the functional effects of mutations to several influenza proteins in mammalian cells (Ashenberg et al., 2017; Bloom, 2014; Doud and Bloom, 2016; Du et al., 2018; Jiang et al., 2016; Lee et al., 2018; Wu et al., 2014, 2015). We show that comparative deep mutational scanning in human versus avian cells identifies numerous human-adaptive mutations that have never before been described. These mutations cluster on the surface of the PB2 protein, highlighting potential interfaces with host factors. Some of these mutations are enriched in avian-human transmission of H7N9 influenza, demonstrating the utility of our experiments for anticipating PB2’s adaptation in nature. The human-adaptive mutations that have not been observed in nature are often inaccessible by single-nucleotide mutations. Overall, our complete map of human-adaptive mutations sheds light on how species-specific selection and evolutionary accessibility shape influenza virus’s evolution to new hosts.

## Results

### Deep mutational scanning of an avian influenza PB2

To identify host-adaptation mutations in PB2, we used deep mutational scanning to measure the effects of all amino-acid mutations to this protein in both human and avian cells. We performed these experiments using the PB2 from an avian influenza strain, A/Green-winged Teal/Ohio/175/1986 (also previously referred to as S009) (Jagger et al., 2010; Mehle and Doudna, 2009). The PB2 from this strain is representative of avian influenza PB2s, most of which are highly similar (average pairwise amino-acid identity of 98.7%) (Fig S1). We mutagenized all codons in PB2 to create three replicate mutant plasmid libraries with an average of 1.4 codon substitutions per clone (Fig 1A, S2A-F). Since there are 759 residues in PB2, there are 759 × 19 = 14,421 amino acid mutations, virtually all of which are represented in our libraries (Fig S2G).

**Figure 1.**
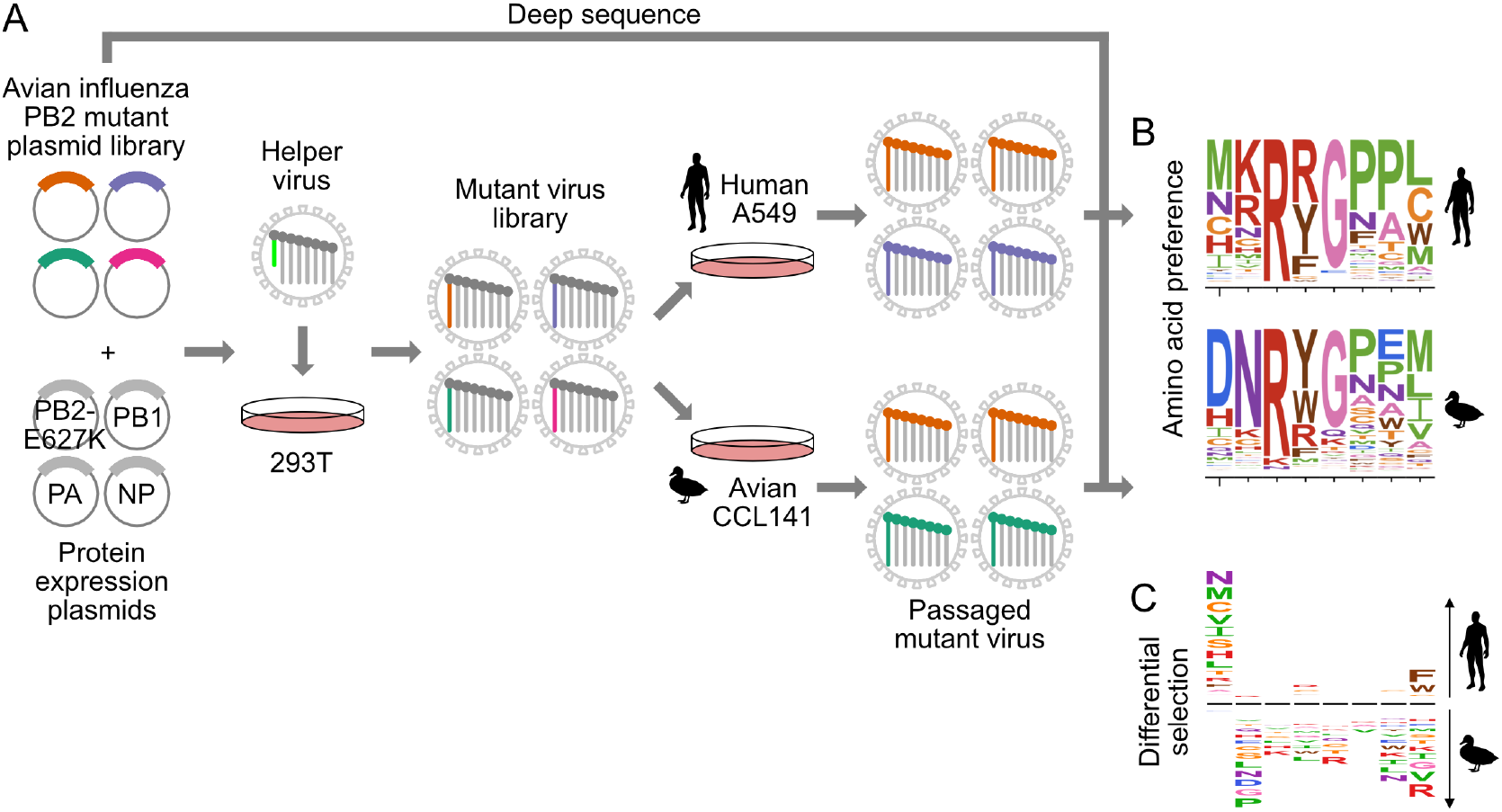
Deep mutational scanning of avian influenza PB2 in human and avian cells. (A) We mutagenized all codons of PB2 from an avian influenza strain. We generated mutant virus libraries using a helper-virus approach, and passaged libraries at low MOI in human (A549) or duck (CCL-141) cells to select for functional PB2 variants. (B) We deep sequenced PB2 mutants from the initial mutant plasmid library and the mutant virus library after passage through each cell type. We computed the “preference” for each amino acid in each cell type by comparing the frequency of each mutation before and after selection. In the logo plots, the height of each letter is proportional to the preference for that amino acid at that site. (C) To identify mutations that are adaptive in one cell type versus the other, we computed the differential selection by comparing the frequency of each amino-acid mutation in human versus avian cells. Letter heights are proportional to the log enrichment of the mutation in human versus avian cells. *See also Figure S1 and S2*.

We generated a mutant virus library from each of the triplicate plasmid mutant libraries using a helper-virus approach, which reduces bottlenecks during generation of complex viral libraries (Doud and Bloom, 2016) (Fig 1A). For biosafety reasons, we rescued reassortant virus using polymerase (PB2, PB1, PA) and nucleoprotein (NP) genes from the avian influenza strain and the remaining viral genes (HA, NA, M, NS) from the lab-adapted A/WSN/1933(H1N1) mammalian influenza strain. We wanted to minimize selection for host-adaptive mutations during the initial library generation. Therefore, we generated the libraries in human HEK293T cells with a co-transfected protein-expression plasmid encoding the human-adapted PB2-E627K protein variant, so that all cells had a PB2 protein that could complement poorly functioning library variants.

To select for functional PB2 variants in human versus avian cells, we passaged each replicate mutant virus library at low MOI in the A549 human lung epithelial carcinoma line and CCL-141 duck embryonic fibroblasts (Fig 1A, S2A). To quantify the functional selection on each mutation during viral growth, we deep sequenced the initial plasmid mutant libraries and the passaged mutant viruses to measure the frequency of mutations before and after selection (Fig 1B). All experiments were also performed in parallel on virus carrying wild-type PB2 as a control to quantify the rate of errors arising during sequencing, library preparation, and viral replication (Fig S2H).

To assess the efficacy of selection without the complication of errors arising from sequencing and passaging, we examined the post-selection frequency of stop and nonsynonymous mutations accessible by >1 nucleotide substitution. Stop and nonsynonymous mutations fell to 2-7% and 26-35% of their initial frequencies respectively (Fig S2H). In contrast, synonymous mutations remained at 68-87% of their initial frequency. Therefore, the experiments effectively selected for functional PB2 mutants.

We quantified selection at the amino-acid level in terms of the “preference” of each site in the protein for each amino acid (Fig 1B) (Bloom, 2015). The preference for an amino acid is proportional to its enrichment during functional selection. We assessed the reproducibility of our experiments across biological replicates by examining the correlations of preferences for all 14,421 amino acid (Fig S2I). Biological replicates passaged in each cell type were well correlated (Pearson’s *R* in human cells was 0.74 to 0.79; Pearson’s *R* in avian cells was 0.76 to 0.79), and were generally better correlated within cell types than between cell types (Pearson’s *R* between cell types was 0.67 to 0.78). For downstream analyses, we rescaled our preferences to match the stringency of selection in nature (see Methods, Table S1, S2).

### Experimental measurements are consistent with natural selection and known functional constraints on PB2

Our experiments reflect known functional constraints on PB2 (Fig 2A, Fig S3). As expected, the start codon shows a strong preference for methionine in both human and avian cells. PB2’s cap-binding function is mediated by a hydrophobic cluster of five phenyalanines (F404, F323, F325, F330, F363), H357, E361, and K376 (Guilligay et al., 2008). Phenylalanines are strongly preferred in the hydrophobic cluster in both host cell types, with the exception of site 323, which also tolerates aliphatic hydrophobic residues in human cells (Fig 2A). E361 is also strongly preferred in both cell types, as is K376 in the duck cells. A number of other amino acids are tolerated at site 376 in human cells, and at site 357 in both cell types. At site 357, aromatic residues tyrosine, tryptophan, and phenylalanine are preferred in addition to histidine, consistent with previous observations that the H357W substitution enhances binding to the m^7^GTP base (Guilligay et al., 2008). Finally, the two motifs comprising the C-terminal bipartitite nuclear import signal, 736-KRKR-739 and 752-KRIR-755 (Tarendeau et al., 2007), are strongly and similarly preferred in both host cell types. Thus, our experimentally measured preferences largely agree with what is known about PB2 structure and function, and further suggest that functional constraints at these critical sites are similar in both human and avian cells.

**Figure 2.**
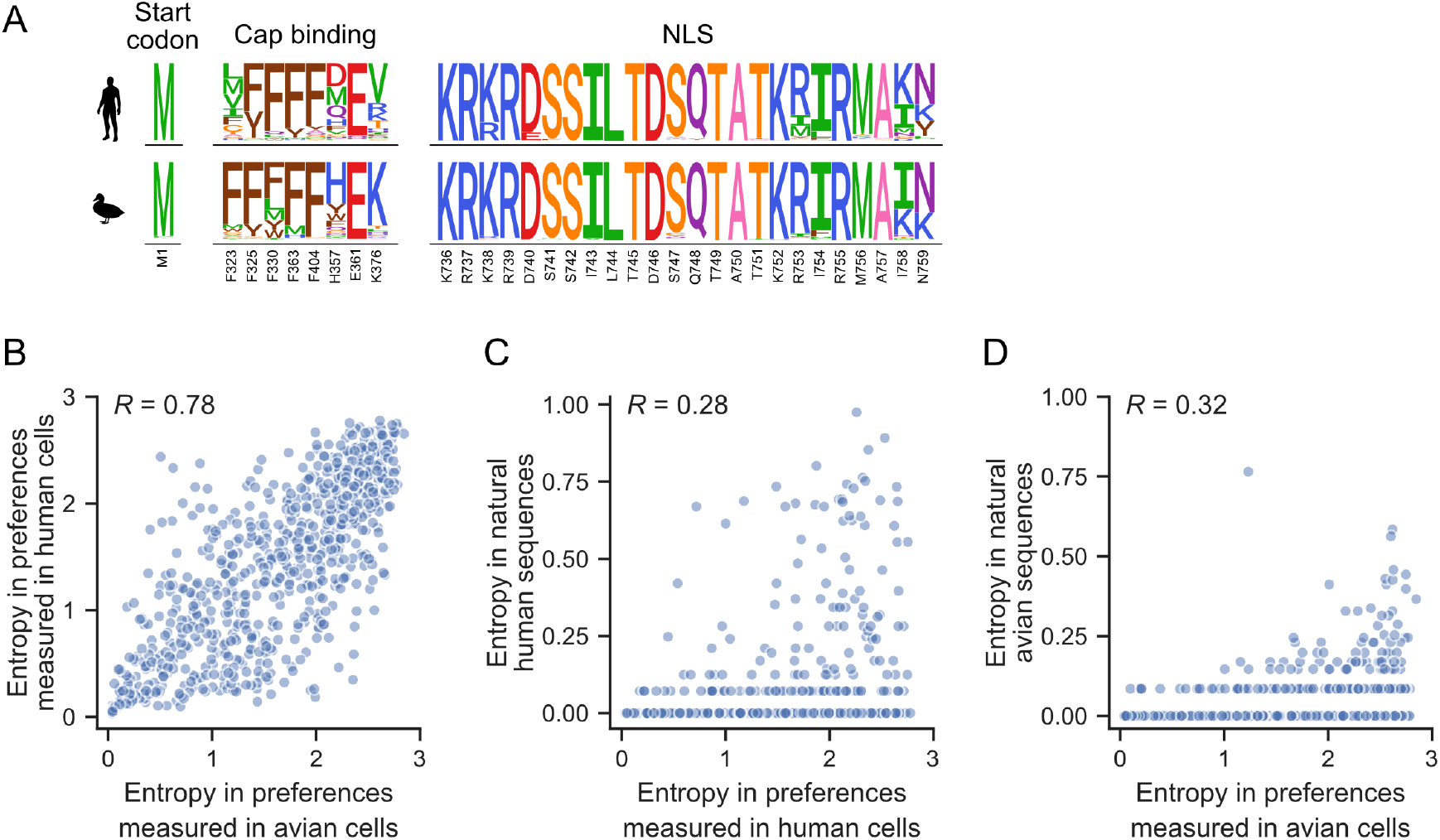
Functional constraints on PB2. (A) The amino acid preferences measured in human and avian cells for key regions of PB2: the start codon, sites involved in cap-binding, and sites comprising the nuclear localization sequence (NLS). The height of each letter is proportional to the preference for that amino acid at that site. Known critical amino acids are generally strongly preferred in both cell types. (B) Correlation of the site entropy of the amino-acid preferences measured in each cell type. (C) Sites of high variability (as measured by entropy) in natural human influenza sequences occur at sites of high entropy as experimentally measured in human cells. (D) Sites with high variability in natural avian influenza sequences occur at sites of high entropy as experimentally measured in duck cells. See also Figure S3 and Table S1, S2.

To more broadly investigate whether functional constraints are similar between both cell types across the entire PB2 protein, we computed the entropy of the amino acid preferences at each site. A larger site entropy indicates a higher tolerance for mutations at that site. Site entropies are well correlated between cell types (Fig 2B, *R* = 0.78), indicating that sites are usually under similar functional constraint in both cell types. These protein-wide measures of mutational tolerance are also consistent with natural sequence variation: sites that are highly variable among publicly available natural influenza sequences tend to also be ones that we experimentally measured to be mutationally tolerant (Fig 2C, D).

### Identification of human-adaptive mutations

To identify mutations that are adaptive in human versus avian cells, we quantified the host-specific effect of each mutation using two different metrics. The first metric, differential selection, quantifies how much a mutation is selected in one condition versus another (Doud et al., 2017). Differential selection is computed by taking the logarithm of the relative enrichment of the mutation relative to the wild-type residue in human versus avian cells (Fig 1C, S4A, Table S2). Differential selection greater than zero indicates that a mutation is relatively more favorable in human than avian cells.

To test if differential selection accurately identifies host-specific mutations, we asked if a set of 24 previously characterized human-or mammalian-adaptive mutations (Table S3) have differential selection values greater than zero. Indeed, most of these previously characterized mutations had positive differential selection values, as expected for human-adaptive mutations (Fig 3A). In contrast, all other mutations have a distribution of differential selection values centered around zero.

**Figure 3.**
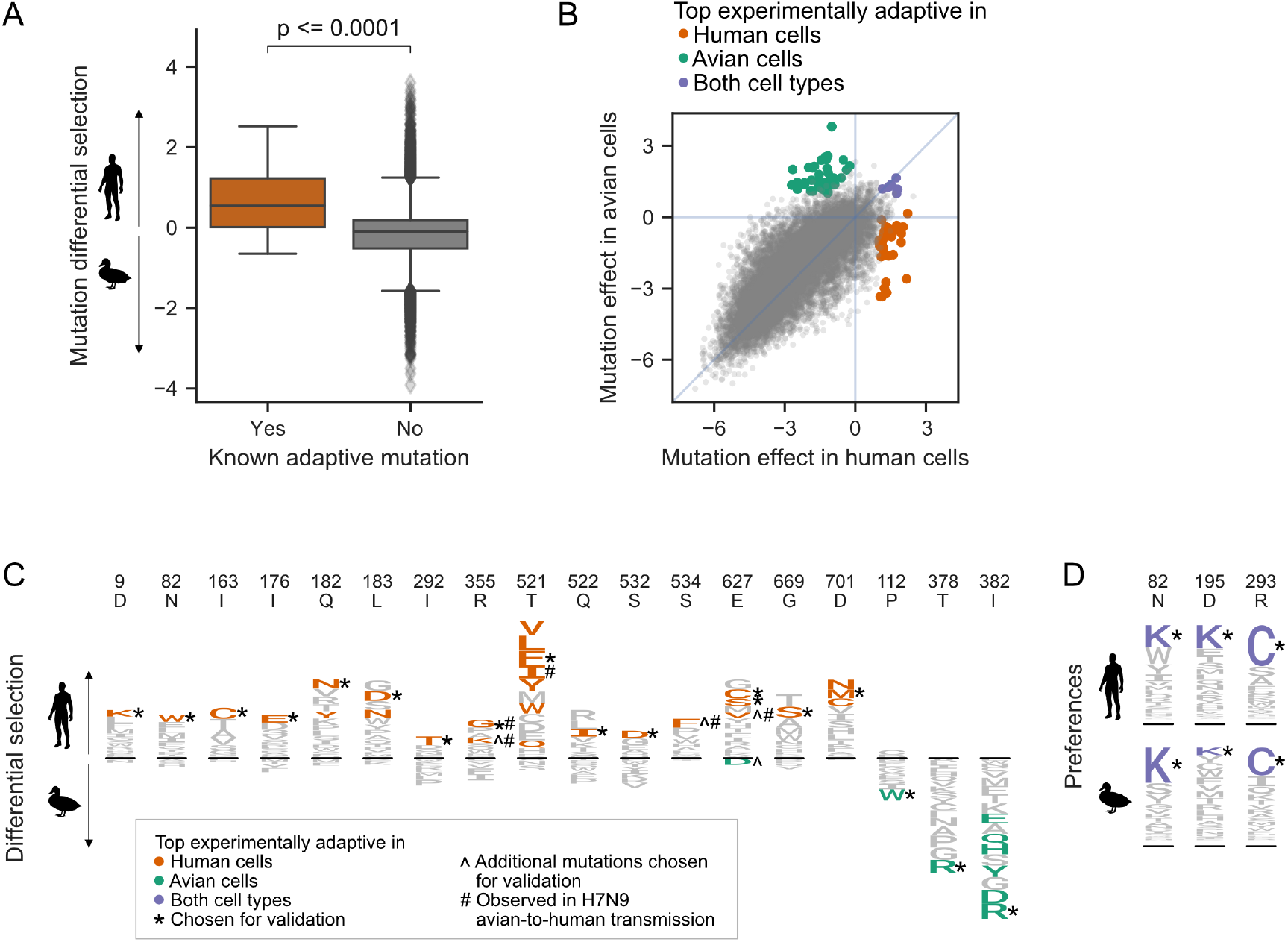
Deep mutational scanning identifies known and novel host-adaptive mutations. (A) Distribution of experimentally measured differential selection for previously characterized human adaptive mutations and all other possible mutations to PB2. Positive differential selection means a mutation is favored in human versus avian cells. (B) Scatterplot of each mutation’s effect in human versus avian cells, showing the top adaptive mutations identified in the deep mutational scanning. (C) Logoplots showing the differential selection at the sites of mutations that we chose for functional validation. The height of each letter above the line indicates how strongly it was selected in human versus avian cells. Top adaptive mutations are colored in orange (human-adaptive) or green (avian-adaptive). Top adaptive mutations chosen for functional validation are indicated by an asterisk(*). Additional mutations chosen for validation are indicated by ^ and are colored orange (human-adaptive) or green (avian-adaptive). Mutations observed in H7N9 avian-to-human transmission are indicated by #. Note that not all mutations with high differential selection in human versus avian cells are classified as top adaptive mutations because we also filtered for mutations that are substantially beneficial relative to wildtype. (D) Logoplots showing amino acid preferences at sites of top mutations measured to be beneficial in both human and avian cells. Mutations that we selected for further validation are colored in purple. *See also Figure S4 and Table S2, S3*.

However, there are many previously uncharacterized mutations that have differential selection values similar to or greater than those of known human-adaptive mutations (Fig 3A). Of course, differential selection only quantifies the extent to which a mutation is more beneficial in human than avian cells. But importantly, for a mutation to be truly adaptive, it must also be more beneficial than the wild-type amino acid in human cells. To quantify each mutation’s effect relative to wild type in each cell type, we computed the logarithm of the ratio of preferences of the mutant versus wild-type amino acid (Fig 3B, Table S3). Mutation effect values greater than zero indicate that a mutation is more preferred than the wild-type residue.

We identified top experimentally adaptive mutations using both differential selection and mutation effect metrics (Fig 3B, S4B). We focused on the 34 mutations most adaptive in our human cell selection (differential selection >1.5 and mutational effect in human cells >1). Among these 34 mutations, only one (D701N) has already been described as human adaptive. The E627K mutation is favored in human cells in our experiments, though it is not in this set of top 34 mutations. However, two other mutations at this site (E627C and E627S) are among the top 34 mutations (Fig 3C). In fact, it appears that many mutations at site 627 are human adaptive, with the exception of E627D. We additionally identified 42 mutations as adaptive in avian cells (differential selection <-2 and mutational effect in avian cells >1), and 7 mutations that are more favorable than the wild-type amino acid in both cell types (mutational effect >1 in both human and avian cells).

From these top adaptive mutations identified in our deep mutational scanning, we chose 26 for experimental validation. Specifically, we chose 18, 4, and 3 mutations adaptive in human, avian, or both cell types, respectively (Fig 3C, D). We prioritized mutations that had consistent measurements across biological replicates. When there were multiple strongly adaptive mutations at a site, we chose just one mutation at that site to test mutations across more sites. Finally, we also validated additional mutations of particular interest, such as those observed in avian-to-human transmission of H7N9 influenza (see below for more details).

### Human-adaptive mutations identified in deep mutational scanning improve polymerase activity and viral growth in human cells

The main function of the influenza polymerase is to transcribe and replicate the viral genome. We quantified the effect of mutations on polymerase activity using a minigenome assay which measures transcription of an engineered viral RNA encoding GFP by reconstituted influenza polymerase. To test whether the results of our deep mutational scanning are generalizable to human cells beyond the A549 cell line used in the scanning, we performed the minigenome polymerase activity assay in HEK293T as well as A549 cells.

Almost all the putative human-adaptive mutations identified in the deep mutational scanning improved polymerase activity in human cells relative to the wild type or a synonymous mutant (Fig 4A, B, Table S4). Mutations that did not improve polymerase activity retained at least wild-type activity. The effect of mutations in both human cell lines were remarkably consistent, suggesting that our deep mutational scanning yielded results that generalize across human cells.

**Figure 4.**
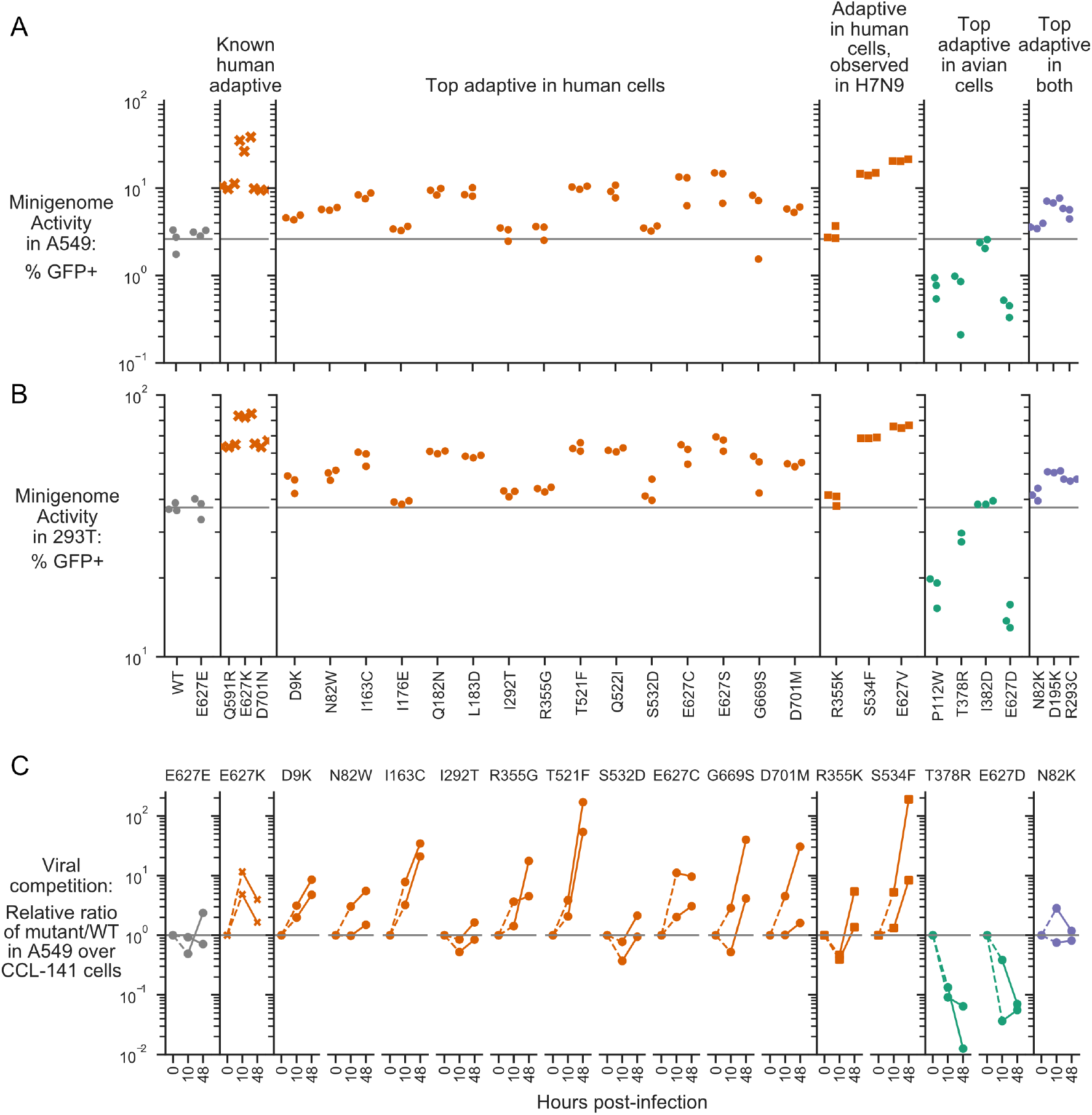
Validation of top experimentally adaptive mutations. The polymerase activity of selected PB2 mutants as measured using mini-genome assays in A549 (A) and HEK293T (B) human cells. Minigenome activity is represented as percent of transfected cells that expressed a viral GFP reporter. The gray horizontal line indicates the mean value measured for the wild type avian PB2. E627E is a synonymous mutation at site 627. (C) Competition of virus bearing the indicated mutant PB2 against virus with wild-type PB2. We mixed the mutant and wild-type viruses at a 1:1 ratio of infectious particles as determined in the respective cell type, infected human A549 and avian CCL-141 cells, and measured the frequency of each variant at 10 and 48 hours using deep sequencing. The plots show the relative ratios of the mutant over wild-type variant in A549 versus CCL-141 cells. These ratios are shown to start at 10^0^ at time zero, since the starting ratio of mutant to wildtype variant in each cell type should be one. Subsequent points above the horizontal line at 10^0^ indicate that a viral mutant grows better in human than avian cells. *See also Table S4 and S5*.

In contrast, all but one of the putative avian-adaptive mutations decreased polymerase activity in human cells compared to wild type, as expected (Fig 4A, B). The one mutation that did not decrease polymerase activity had comparable activity to wild type. Finally, mutations that are putatively adaptive in both human and avian cells had modestly improved or comparable polymerase activity in human cells compared to wild type.

We also tested the effect of some of the mutations on viral growth in order to capture any effects of mutations beyond polymerase activity. We performed viral growth assays by competing virus carrying each mutant PB2 against wild-type virus. We mixed mutant and wild-type virus at a 1:1 ratio, infected human (A549) or avian (CCL-141) cells at low MOI, and then measured the frequencies of mutant to wild-type virus at 10 and 48 hours post infection by deep sequencing.

Almost all putative human-adaptive mutations identified in the deep mutational scanning improved growth in human over avian cells, as reflected by an increase in the ratio of mutant to wild type in human cells versus avian cells over the time course of the competition (Fig 4C, Table S5). One of the putative human-adaptive mutations that did not improve polymerase activity (R355G) improved growth in human over avian cells at both 10 and 48 hours post-infection. An additional three of the putative human-adaptive mutations that did not have an effect on polymerase activity (I292T, S532D, R355K) improved growth in human cells by 48 hours compared to 10 hours post-infection. Therefore, these four mutations confer a human-specific growth benefit due to some mechanism other than polymerase activity. As expected, both putative avian-adaptive mutations resulted in poorer growth in human versus avian cells. Finally, the N82K mutation that is putatively adaptive in both human and avian cells resulted in comparable growth in both human and avian cells, as expected.

Thus, our deep mutational scanning identified numerous previously undescribed PB2 mutations that improve polymerase activity or viral growth in human cells. In addition, it also identified an intriguing small set of mutations that enhance viral growth but not polymerase activity in human cells.

### Human-adaptive mutations cluster in regions of PB2 that are potentially important for host adaptation

The additional human-adaptive mutations we identified may improve PB2’s ability to interact with important human cell factors. To identify potential interfaces for such interactions, we mapped the sites of top human-adaptive mutations identified in the deep mutational scanning onto the structure of PB2. Many of the sites cluster in regions of PB2 that may play a role in host adaptation (Fig 5, S5).

**Figure 5.**
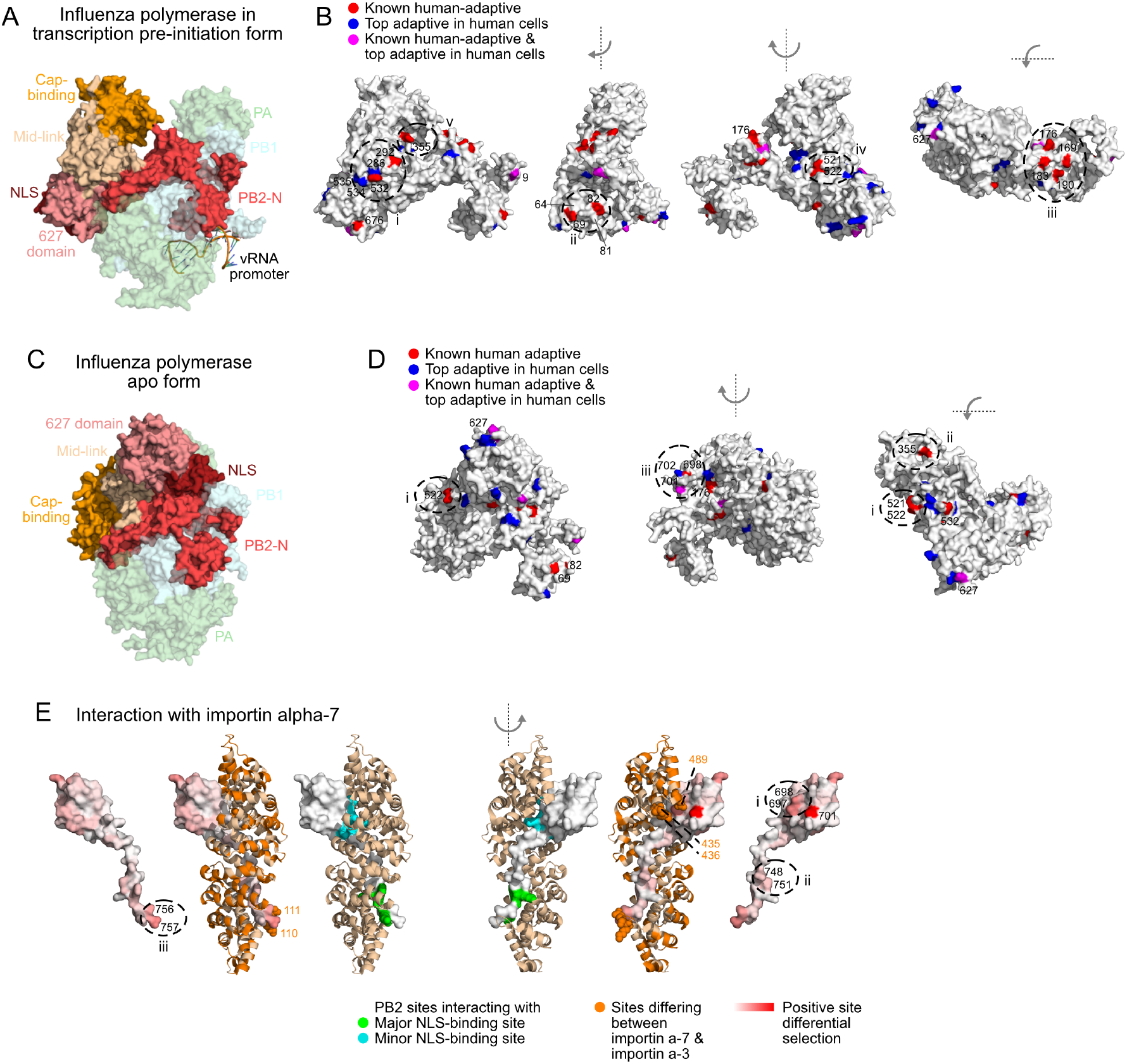
Locations of top human-adaptive mutations on the structure of the influenza polymerase. Overall structure of the influenza polymerase complex comprising PB2, PB1 and PA in (A, B) the transcription pre-initiation form (PDB: 4WSB) and (C, D) the apo form (PDB: 5D98). PB2 domains defined as in Pflug et al. (2017). (B, D) Sites of top human-adaptive mutations identified by deep mutational scanning are shown in red on the PB2 subunit of the structure. Sites of previously known human-adaptive mutations are in blue. Sites identified by deep mutational scanning and which were also previously known are in purple. Surface-exposed adaptive sites, particularly those forming clusters, are circled and numbered. (E) Structure of PB2 C-terminal fragment co-crystalized with importin-α 7 (PDB: 4UAD). Sites on PB2 interacting with major and minor NLS binding surfaces of importin-α7 are in green and cyan respectively. Importin-α7 is depicted in ribbon form in tan. We used the deep mutational scanning to define a continuous variable indicating the extent of host-specific adaptation at each site of PB2. Specifically, for each site, we computed the positive site differential selection by summing all positive mutation differential selection values at that site (i.e., the total height of the letter stack in the positive direction in logoplots such as in Fig 3D). We mapped this differential selection onto the PB2 C-terminal fragment in red; PB2 sites with high differential selection are numbered. Regions of importin-α7 that differ from importin-α3 are colored in orange, those near PB2 sites with high differential selection are shown as spheres. For all structures, the avian influenza (S009) PB2 amino acid sequence was mapped onto the PB2 chain by one-2-one threading using Phyre2 (Kelley et al., 2015) (Confidence in models for 4WSB, 5D98, and 4UAD are 100%, 100%, and 99% respectively). Sites are numbered according to the S009 PB2 sequence. *See also Figure S5*.

Specifically, in the transcription pre-initiation form of the polymerase (PDB: 4WSB) (Reich et al., 2014) (Fig 5A, B), top sites 532 and 292 occur near sites of known human-adaptive sites 286, 534, and 535 (Cauldwell et al., 2013; Mänz et al., 2016) (Fig 5B: i). Top sites 69 and 82 occur close to sites 64 and 81 (Fig 5B: ii), which are located near the template exit channel and were recently shown to modulate generation of mini viral RNAs that act as innate-immune agonists (te Velthuis et al., 2018). We also find a cluster of top sites in the PB2-N terminal domain (169, 176, 183, 190, Fig 5B: iii) that is partially occluded by the flexible PA endonuclease domain in the transcription preinitiation structure. The influenza polymerase, particularly the domains of PB2 outside of the N-terminal domain, undergoes dramatic conformational rearrangement between the transcription pre-initiation and apo form (PDB: 5D98) (Hengrung et al., 2015). Some sites, such as the pair 521 and 522, and 355, which face the product exit channel and core of the polymerase in the transcription pre-initiation structure (Fig 5B: iv, v), are more fully surface-exposed in the apo structure (Fig 5C, D: i, ii). Similar results are obtained if we instead analyze the structures in terms of a continuous variable representing the extent of human-specific adaptation at each site (Fig S5). Taking a comprehensive approach to identify human-adaptive mutations has therefore allowed us to map surfaces of PB2 that might mediate host-interactions.

Next, we asked if host-adaptation mutations occur at known interaction interfaces with host proteins. One known interacting host protein is importin-a, which mediates nuclear import of PB2 and has been proposed to have a role in viral transcription and replication (Resa-Infante et al., 2008; Tarendeau et al., 2007). PB2 of avian viruses uses importin-a3 in human cells, whereas PB2 of mammalian-adapted viruses uses importin-α7 (Gabriel et al., 2008, 2011). We mapped total positive differential selection on each site of PB2 (see legend for Fig 5E), and asked how this selection on PB2 relates to its interaction with importin-α. Sites on PB2 that interact with the major and minor NLS-binding surfaces of importin-α (Pumroy et al., 2015; Tarendeau et al., 2007) generally have low differential selection, indicating that host-adaptation mutations do not occur at these sites (Fig 5E; PDB: 4UAD) (Pumroy et al., 2015). This is expected, since all importin-α isoforms share an invariant NLS-binding surface. However, adjacent PB2 sites have higher differential selection (Fig 5E: i-iii). Some of these PB2 sites are in close proximity to regions of importin-α that differ between the α-7 and α-3 isoforms (Fig 5D: i, iii), suggesting that adaptation at these PB2 sites affects importin-α usage.

PB2 also interacts with the C-terminal domain of RNA polymerase II, and this interaction is proposed to stabilize the polymerase in the transcription-competent conformation (PDB: 6F5O) (Serna Martin et al., 2018). Similar to what we observe with importin-α, PB2 sites thought to directly interact with RNA polymerase II tend to have low differential selection, whereas adjacent PB2 sites have higher differential selection (Fig S5C). Thus, it appears that host adaptation may involve mutations at sites adjacent to the core residues that directly interact with host proteins.

### Experimentally defined human-adaptive mutations are enriched in avian-to-human transmission of H7N9 influenza

A challenge in the surveillance of non-human influenza and assessment of pandemic risk is determining which of the many mutations that occur during viral evolution are human-adaptive (Lipsitch et al., 2016; Russell et al., 2014). We investigated whether our experimental measurements can identify host-adaptation mutations that occur during the actual transmission of avian influenza to humans.

Avian H7N9 influenza viruses have recently caused a large number of sporadic human infections (Su et al., 2017). We examined mutations occurring during the evolution of H7N9 viruses that have jumped from avian to human hosts to determine whether they were enriched for changes predicted by our deep mutational scanning to be human adaptive. First, we constructed a phylogeny of H7N9 PB2 sequences, inferred ancestral sequences for all internal nodes, and assigned mutations to specific branches of the phylogenetic tree (Fig 6A). We then classified each mutation on the phylogenetic tree as “avian” or “human” based on whether it occurred on a branch connecting two avian isolates, or on a branch leading to a human isolate respectively (Table S6). As human infections by H7N9 are evolutionary dead-ends, mutations occurring during human infections should appear immediately proximal to human isolates in the phylogeny, while mutations occurring during bird infections will be ancestral.

**Figure 6.**
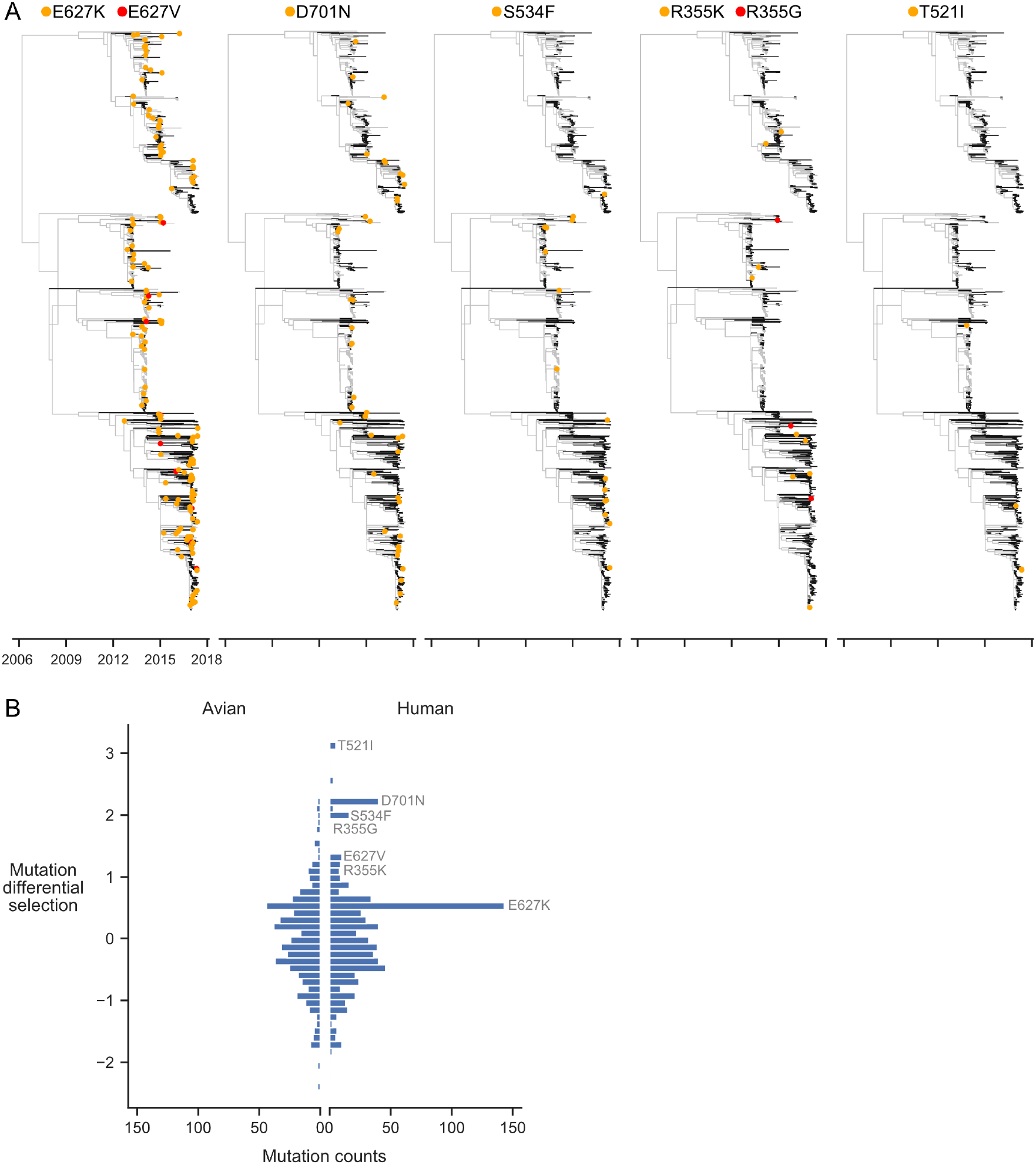
Experimentally identified human-adaptive mutations are enriched in avian-human transmission of H7N9 influenza. (A) Phylogeny of H7N9 influenza PB2 sequences. Branches in human and avian hosts are colored black and grey respectively. Orange or red dots indicate where a mutation was inferred to have occurred. Branch lengths are scaled by annotated and inferred dates of origin of each sequence. (B) Distribution of experimentally measured differential selection values for mutations occurring during H7N9 evolution in human and avian hosts. A positive differential selection value means that our experiments measured the mutation to be beneficial in human versus avian cells. Top differentially selected mutations that occur frequently are labeled. *See also Table S6*.

We next asked whether mutations occurring in human hosts had higher differential selection values in our deep mutational scanning than mutations in avian hosts. Indeed, human mutations more often had high differential selection (a value >0.5) than avian mutations (Fisher’s exact test, p = 2.69e-7) (Fig 6B). The H7N9 human mutations with differential selection >0.5 include the well-studied human-adaptive mutations 627K and 701N. Indeed, these two mutations make up the majority of H7N9 human mutations with high differential selection. But we also identified a number of other mutations with high differential selection that occurred at least four independent times in jumps of H7N9 influenza into humans: 627V, 534F, 355K, and 521I (Fig 6A, 6B, 3D), only one of which has been previously characterized (A627V, Taft et al., 2015). A second mutation at site 355 (355G) with high differential selection also occurs during jumps of H7N9 influenza into humans. Thus, our deep mutational scanning identifies both previously characterized and novel mutations that occur in natural avian-to-human transmission of influenza.

### Most human-adaptive mutations are not accessible by single nucleotide substitutions

Our deep mutational scanning identifies many human-adaptive mutations. Why do we not observe all of them in nature? One possible explanation is that some of these mutations are inaccessible by single nucleotide substitution from existing sequences, and are therefore less likely to arise during natural evolution (Fragata et al., 2018).

We examined if accessibility by single nucleotide substitution imposes constraint on which human-adaptive mutations arise in nature. To do so, we calculated the mean nucleotide substitutions required to access known human-adaptive mutations from all avian influenza PB2 sequences collected in the past three years. This mean number of nucleotide substitutions can range from less than one (if the mutation is already present in some avian PB2 sequences) to three (if the mutation requires three nucleotide changes from all avian PB2 sequences). The majority of previously characterized human-adaptive mutations are accessible by single nucleotide substitutions from avian PB2 sequences (Fig 7), suggesting that these mutations have already been characterized because they readily occur in the context of current avian influenza viruses. In contrast, most of the top human-adaptive mutations identified in our deep mutational scanning require multiple nucleotide substitutions from current avian PB2 sequences (Fig 7). Therefore, many of the novel human-adaptive mutations uncovered by our experiment have probably not been previously identified because they are evolutionarily inaccessible from current avian influenza sequences.

**Figure 7.**
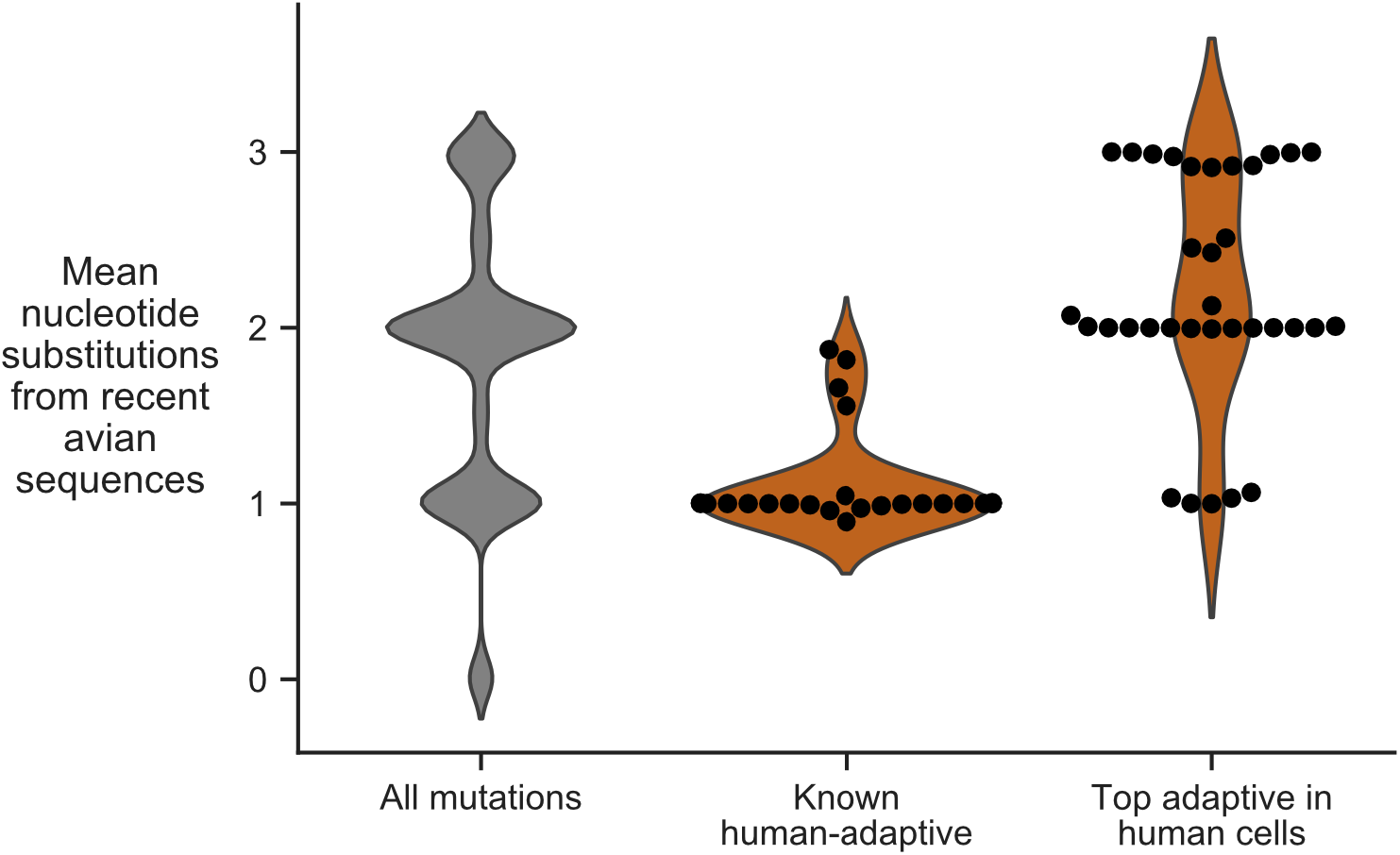
Evolutionary accessibility of mutations from current avian influenza PB2 sequences. Distribution of mean nucleotide substitutions required to access all amino-acid mutations, previously characterized human-adaptive mutations, and top human-adaptive mutations identified in our deep mutational scanning. Mean nucleotide substitution is calculated by averaging over all avian influenza PB2 sequences collected from 2015 to 2018. Most previously characterized human-adaptive mutations are accessible by single nucleotide substitution, whereas many of the new adaptive mutations that we identified require multiple nucleotide substitutions.

The importance of evolutionary accessibility is especially obvious if we examine the adaptive mutations that actually occur in natural influenza virus evolution. Of the five top human adaptive mutations accessible by single nucleotide substitution (Fig 7), three have occurred repeatedly during recent transmissions of H7N9 influenza to humans (355G, 521I, 701N; see Fig 6A). Thus, it may be possible to combine deep mutational scanning measurements of phenotypes with analyses of evolutionary accessibility to anticipate which mutations are most likely to arise in the context of any given starting sequence (Fragata et al., 2018).

## Discussion

We have measured how all single amino-acid mutations to an avian influenza PB2 affect viral growth in both human and avian cells. Our results separate the constraints on PB2 into those that are maintained across cells from diverse species versus those that are specific to human or avian cells. The vast majority of sites are under extremely similar constraint in human and avian cells, including at residues already known to be critical for PB2 function. Layered upon this conserved constraint are mutations with host-specific effects. Our approach therefore represents a powerful strategy for mapping viral determinants of cross-species transmission.

While our complete experimental mapping of human-adaptive mutations does not directly explain how these mutations act, it does provide molecular footholds for investigating their mechanism. Specifically, human-adaptive mutations cluster on the structure of PB2, highlighting regions of the protein that impact host adaptation. Further, we identify some novel human-adaptive mutations that improve viral growth but not polymerase activity in human cells. We speculate that the effects of these mutations on viral growth is mediated by other effects of PB2, such as in modulating the innate-immune response (Graef et al., 2010; te Velthuis et al., 2018).

We also show that our comprehensive experimental measurements can identify human-adaptive mutations that occur during avian-to-human transmission of H7N9 influenza. These measurements therefore help address a fundamental challenge in assessing the risk of potential pandemic influenza virus strains: determining which of the many mutations observed during viral surveillance affect whether a virus will be successful in human hosts (Lipsitch et al., 2016; Russell et al., 2014). Our high-throughput approach therefore enables phenotypic measurements to keep pace with the challenge of interpreting the many viral mutations that are observed during genotypic surveillance of virus evolution (Grubaugh et al., 2019).

Some amino-acid mutations we experimentally identified as human-adaptive occur frequently in nature, but others have never been observed. Most human-adaptive mutations observed in natural influenza evolution are accessible by single nucleotide substitution from current avian genotypes, showing that evolutionary accessibility shapes which mutations occur during viral adaptation (Fragata et al., 2018). But of course, the evolutionary accessibility of codon mutations can change over time as avian influenza sequences themselves evolve. Integrating our complete maps of the effects of amino-acid mutations with other data that sheds light on evolutionary opportunity, such as nucleotide accessibility, mutation rates, transmission bottlenecks, and environmental and epidemiological factors (Geoghegan and Holmes, 2017; Geoghegan et al., 2016; Moncla et al., 2016; Olival et al., 2017; Peck and Lauring, 2018; Varble et al., 2014) will help us understand how viruses cross species barriers in nature.

## Acknowledgements

We thank William Fowler for help with molecular cloning, Juhye Lee, Adam Dingens, and John Huddleston for advice and computer code, and Harmit Malik, Katherine Xue, Allison Greaney, and Tyler Starr for providing comments on this manuscript. We thank Jeffrey Taubenberger for providing plasmids for A/Green-winged Teal/Ohio/175/1986 (S009), and Andrew Mehle and Steven Baker for advice about growing virus containing S009 polymerase and NP genes.

YQSS was supported by the Mahan Postdoctoral Fellowship (Computational Biology Program of Fred Hutchinson Cancer Research Center) and the Damon Runyon Postdoctoral Fellowship. This work was supported by NIH grant R01 AI127893 from the NIAID to TB and JDB, and NIH grant R35 GM119774-01 from the NIGMS to TB. TB is a Pew Biomedical Scholar. JDB is an Investigator of the Howard Hughes Medical Institute.

## Author Contributions

Conceptualization, YQSS, JDB; Methodology, YQSS, LHM, TB, JDB; Validation, YQSS, RE; Investigation, YQSS, LHM, RE; Software, YQSS, LHM, JDB; Writing – Original Draft, YQSS; Writing – Review & Editing, YQSS, LHM, RE, TB, JDB; Supervision, TB, JDB; Funding Acquisition, TB, JDB

## Declaration of Interests

The authors have no competing interests.

## STAR Methods

### Contact for reagent and resource sharing

Further information and requests for resources and reagents should be directed to and will be fulfilled by the Lead Contact, Jesse D. Bloom.

### Experimental model and subject details

#### Cell lines and media

HEK293T, MDCK-SIAT1, and A549 (ATCC CCL-185) cells were maintained in D10 media (DMEM supplemented with 10% fetal bovine serum, 2 mM L-glutamine, 100 U/ml penicillin, and 100 μg/ml of streptomycin). CCL-141 cells (ATCC CCL-141) were maintained in E10 media (identical to D10 except that EMEM is used in place of DMEM). Cells were grown in WSN growth media (WGM: Opti-MEM supplemented with 0.5% FBS, 0.3% BSA, 100ug/ml CaCl_2_, 100 U/ml penicillin, and 100 μg/ml of streptomycin) for viral infections.

For an avian cell line, we chose to use a duck rather than a chicken cell line because ducks are natural hosts of influenza that (unlike chickens) possess RIG-I, a key innate-immune sensor of influenza (Barber et al., 2010).

For expansion of helper virus, we generated MDCK-SIAT1 cells expressing S009 PB2-E627K under control of a doxycycline-inducible promoter (MDCK-SIAT1-tet-S009-PB2-E627K) using a Sleeping Beauty transposon system (Kowarz et al., 2015). Briefly, MDCK-SIAT1 cells were transfected with pSBtet_RP_S009_PB2_E627K and pSB100X transposase vector using Lipofectamine 3000 (ThermoFisher Scientific, L3000015), and then subject to selection with 1 ug/ml puromycin. At three days post-transfection, we sorted for individual transfected cells expressing mCherry. All subsequent experiments were performed with a clonal expansion of a single transfected cell.

All cell lines tested negative for mycoplasma at the time they were expanded for either generating helper virus or passaging mutant plasmid libraries.

### Method details

#### Plasmids

Sequences for plasmids generated in this study are provided in File S1.

Avian influenza polymerase plasmids: Original plasmids for PB2, PB1, PA, and NP genes from avian influenza strain A/Green-winged Teal/Ohio/175/1986 (S009) were gifts of Jeffrey Taubenberger (Jagger et al., 2010). For generating the mutant plasmid library, we cloned the S009 PB2 coding sequence into a pHW2000 vector (Hoffmann et al.,

2000) from which we removed the CMV promoter (final plasmid pHW_noCMV_S009_PB2). The mutant library insert was cloned into the recipient vector pHW_noCMVnoTerm_BsmBI, which lacks the Pol I terminator (terminator sequence is part of the insert). The reason that we generated a pHW plasmid without a

CMV promoter is that we were unable to maintain a stable bacterial clone of the S009 PB2 coding sequence on the pHH21 plasmid backbone - we observed frequent deletions in the coding sequence during plasmid propagation, suggesting that the insert on the pHH21 plasmid backbone is toxic to the bacterial host. For generating helper virus and virus for viral competitions, we cloned the S009 PB2, PB1, PA, and NP coding sequences into pHW2000 (pHW_S009_PB2, pHW_S009_PB1, pHW_S009_PA, pHW_S009_NP). In all cases we used non-coding viral-RNA termini from the respective A/WSN/1933(H1N1) gene segment. For protein expression and the minigenome assay, we cloned the PB2, PB1, PA, and NP coding sequences from S009 into a protein-expression plasmid with a CMV promoter (HDM_S009_PB2, HDM_S009_PB1, HDM_S009_PA, HDM_S009_NP). All mutants of PB2 were made by site-directed mutagenesis on the appropriate plasmid backbone.

Helper virus plasmids: To generate a PB2 vRNA lacking a functional PB2 protein, we cloned GFP flanked by PB2 sequence into the pHH21 vector (Neumann et al., 1999) (pHH_PB2_S009_flank_99_eGFP_100). The flanking non-coding viral-RNA termini are from WSN PB2, and the coding sequences are from S009 PB2. The length of flanking sequences, 99 and 100 bases on the 5’ and 3’ end of the PB2 coding sequence respectively, are based on prior experiments analyzing how much terminal sequence is needed for effective genome packaging (Liang et al., 2005). We mutated out start codons 5’ to the GFP start site in the mRNA sense. Our helper virus rescue also required the reverse genetics plasmids encoding HA, NA, M, and NS from WSN (pHW184_HA, pHW186_NA, pHW187_M, pHW188_NS) (Hoffmann et al., 2000). To generate a cell line with doxycycline-inducible expression of S009 PB2 for expansion of PB2-deficient helper virus, we cloned the S009 PB2-E627K coding sequence into the pSBtet vector (pSBtet_RP_S009_PB2_E627K) (Kowarz et al., 2015).

Minigenome assay: In addition to protein expression plasmids described above, we used a pHH-PB1-flank-eGFP reporter (Bloom et al., 2010), and pcDNA-mCherry as a transfection control (pcDNA3.1_mCherry).

#### Primers

All primer sequences used in this study are provided in Table S7. Note that this Excel file has several worksheets giving primers for different aspects of the experiments.

#### PB2 codon mutant plasmid libraries

We generated all possible codon mutations of the entire PB2 coding sequence using the PCR-based strategy described in Bloom (2014) with the modifications described in Dingens et al. (2017). Briefly, we designed mutagenic primers tiling across the entire coding region (https://github.com/jbloomlab/CodonTilingPrimers). We performed 10 cycles of fragment PCR using the mutagenic primers and end primers flanking the vRNA, followed by 20 cycles of joining PCR using only end primers (Table S7: Mutagenesis worksheet). We generated three independent libraries starting from mutagenesis of independent bacterial clones. The PB2 variants were cloned into the BsmBI-digested vector pHW_noCMVnoTerm_BsmBI using NEBuilder HiFi DNA Assembly Master Mix (NEB, E2621S), and electroporated into ElectroMAX DH10B competent cells (Invitrogen, 18290015). We obtained 18-22 million transformants for each replicate library, from which we extracted plasmid by maxiprep. We randomly selected 48 clones for Sanger sequencing to evaluate the library mutation rate (https://github.com/jbloomlab/SangerMutantLibraryAnalysis) (Fig S1A-F).

#### Generation and passaging of mutant virus libraries

We generated mutant virus libraries using the helper-virus approach in Doud and Bloom (2016), with modifications. We rescued reassortant virus using polymerase and nucleoprotein genes (PB2, PB1, PA, NP) from S009, and remaining genes (HA, NA, M, NS) from A/WSN/1933(H1N1) (WSN).

##### Helper virus

We plated a co-culture of 4 × 10^5^ HEK293T and 0.5 × 10^5^ MDCK-SIAT1-tet-S009-PB2-E627K in D10 media per well of a 6-well plate. At 18 hours after seeding cells, we added 1 ug/ml doxycycline to induce PB2 expression. One hour after adding doxycycline, we transfected each well with 250 ng each of pHH_PB2_S009_flank_99_eGFP_100, HDM_S009_PB2-E627K, pHW_S009_PB1, pHW_S009_PA, pHW_S009_NP, pHW 184_HA, pHW186_NA, pHW187_M, and pHW188_NS using BioT (Bioland Scientific, B01-01). At four hours post-transfection, we replaced D10 media with WGM supplemented with 1 ug/ml doxycycline. We collected viral supernatant 52 hours post-transfection. To expand the helper virus, we seeded MDCK-SIAT1-tet-S009-PB2-E627K cells in D10 media 4 hours prior to infection at 4 × 10^6^ cells per 15 cm dish. We then infected each dish with 40 ul of fresh viral supernatant, using WGM supplemented with 1 ug/ml doxycycline to induce PB2 expression. At 48 hours post-infection, we collected viral supernatant containing expanded helper virus and clarified the supernatant by centrifugation at 400x g for 4 min. We measured the infectious particle (IP)/ul titer of the helper virus by infecting HEK293T cells with a known volume of viral supernatant, and quantifying the number of GFP+ cells by flow cytometry 18 hours post-infection.

##### Mutant virus library rescue

For each mutant plasmid library, we seeded 36 wells of a 6-well dish with 1 × 10^6^ HEK293T cells, and transfected each well 17 hours later with 375 ng each of HDM_S009_PB2-E627K, HDM_S009_PB1, HDM_S009_PA, HDM_S009_NP, and 500 ng of PB2 mutant plasmid library using BioT. For the wild-type control, we seeded 6 wells and used pHW_noCMV_S009_PB2 in place of the mutant plasmid library. At 6 hours post-transfection, we infected cells with helper virus at MOI of 1 IP/cell in WGM. At 2 hours post-infection, we replaced the inoculum with fresh WGM. At 20 hours post-infection, we harvested viral supernatants and clarified the supernatant by centrifugation at 400x g for 4 min. Supernatants were titered by TCID_50_ on MDCK-SIAT1 cells. The titers for the three library replicates and wild-type control were 262, 68, 100, and 1467 TCID_50_/ul respectively.

##### Passaging of mutant virus libraries

We aimed to passage 1 × 10^6^ TCID_50_ of each mutant virus library in A549 and CCL-141 cells at MOI of 0.01 TCID_50_/cell as determined in MDCK-SIAT1 cells. Therefore, we aimed to have 1 × 10^8^ total cells each at the time of infection. The day prior to each infection, we seeded between 8 × 10^7^ to 1 × 10^8^ A549 cells in D10 in 4 × 5-layered cell culture flasks (Corning, 353144), and 8 × 10^7^ to 1 × 10^8^ CCL-141 cells in E10 in 8 × 5-layered cell culture flasks. To estimate the total number of each cell type at the time of each infection, we plated an equivalent density of cells in a T225 flask. Just prior to infection, we counted the number of cells in the T225 flask, and extrapolated the total number of cells to be infected. We calculated, for an MOI of 0.01, the number of TCID_50_s to be passaged for each of the three library replicates and wild-type control in A549 cells to be 9.39 × 10^5^, 1.02 × 10^6^, 1.20 × 10^6^, and 9.30 × 10^5^ respectively, and in CCL-141 cells to be 8.46 × 10^5^, 8.92 × 10^5^, 9.54 × 10^5^, and 8.58 × 10^5^ respectively. We infected cells by removing D10 or E10 media, rinsing each flask with PBS, and then adding the calculated amount of virus diluted in WGM. At 3 hours post-infection, we replaced the inoculum with fresh WGM. We harvested viral supernatant 48 hours post-infection, and clarified the supernatant by centrifugation at 400x g for 4 min.

#### Barcoded subamplicon sequencing

We ultracentrifuged clarified viral supernatant at 27,000 rpm in a Beckman Coulter SW28 rotor, for 2 hours at 4ºC. We resuspended the virus the residual media, then extracted RNA from 280 ul of concentrated virus using the Qiagen RNeasy Mini Kit (Qiagen, 74104). We titered the concentrated viral supernatant by TCID_50_ on MDCK-SIAT1 cells to estimate the total TCID_50_ from which we extracted RNA (ranged from 2.80 × 10^5^ to 8.8 × 10^6^ TCID_50_); we expect the TCID_50_ titer to be a lower-bound and underestimate of the total viral variants present in the supernatant, since it measures only infectious virus, whereas we would extract RNA from both infectious and non-infectious virions.

We used a barcoded-subamplicon deep sequencing strategy that reduces the sequencing error rate (Doud and Bloom, 2016, https://jbloomlab.github.io/dms_tools2/bcsubamp.html). Briefly, we reverse transcribed the full PB2 vRNA with Accuscript Reverse Transcriptase (Agilent, 200820) (primer S009-PB2-full-1F, Table S7: Barcoded subamplicon sequencing worksheet), and then PCR amplified the full PB2 vRNA (primers S009-PB2-full-1F and S009-PB2-full-8R, Table S7) using KOD Host Start Master Mix (EMD Millipore, 71842), making sure to have amplified from an estimated 1 × 10^7^ cDNA molecules. We then PCR amplified the PB2 gene in 8 subamplicons using primers containing a random barcode to uniquely identify each template cDNA molecule (Table S7). Approximately 7.5 × 10^5^ uniquely barcoded molecules from each subamplicon library were then amplified by primers that add Illumina sequencing adaptors (Table S7). Finally, these libraries were deep sequenced on an Illumina HiSeq 2500 using 2 × 250 bp paired-end reads to a target 3.3x coverage per barcode.

#### Analysis of deep mutational scanning data

Deep mutational scanning sequence data was analyzed using dms_tools2 (https://jbloomlab.github.io/dms_tools2, version 2.3.0). The GitHub repository https://github.com/jbloomlab/PB2-DMS contains Jupyter notebooks that perform all steps of the analyses and provide detailed step-by-step explanations and plots. The README file explains the organization of the notebooks and other files. HTML renderings of the notebooks are provided in File S2. Processed results on preferences, differential selection, and mutation effect are provided in Table S3.

##### Rescaling of preferences

We rescaled our preferences to match the stringency of selection in nature. To do so, we first asked how well the preferences measured by the deep mutational scanning in either human or avian cells describes evolution of PB2 in both human and avian hosts. We used the preferences to generate an experimentally informed codon substitution model (ExpCM) (Bloom, 2017; Hilton et al., 2017), and asked if the ExpCMs described PB2’s natural evolution better than a standard phylogenetic substitution model. The ExpCMs using amino-acid preferences vastly outperformed standard phylogenetic substitution models, suggesting that our experiments do capture some of the natural evolutionary constrain on PB2 (Table S1). The ExpCM stringency parameter had a value of 2.5, indicating that natural selection favors the same amino acids as our experiments, but with greater stringency (Hilton et al., 2017). We thus rescaled our preferences to match the stringency of selection in nature using the ExpCM stringency parameter, and use these rescaled preferences for all subsequent analyses (Fig S3, Table S1, S2).

#### Minigenome activity

Minigenome assays were performed in biological triplicate (starting from independent bacterial clones of each PB2 mutant) in both A549 and HEK293T cells. We seeded 2.5 × 10^4^ A549 or HEK293T cells per well of a 96-well plate. Cells were transfected the next day with 10 ng each of HDM_S009_PB2 (for the respective mutant), HDM_S009_PB1, HDM_S009_PA, HDM_S009_NP, 30 ng of pHH-PB1-flank-eGFP reporter, and 30 ng of pcDNA-mCherry as transfection control, using Lipofectamine 3000 (A549) or BioT (HEK293T). At 22 hours post-transfection, cells were trypsinized and analyzed by flow cytometry. We report minigenome activity as the percent of mCherry-positive cells that are GFP-positive.

#### Viral competition

##### Mutant virus

We generated mutant virus by reverse genetics using pHW_S009_PB2 (for the respective mutant), pHW_S009_PB1, pHW_S009_PA, pHW_S009_NP, pHW184_HA, pHW186_NA, pHW187_M, and pHW188_NS. We seeded 5 × 10^5^ HEK293T cells per well of a 6-well plate, and transfected cells the next day with 250 ng each of the 8 pHW plasmids using BioT. At 2 hours post-transfection, we replaced media with WGM. At 40 hours post-infection, we collected viral supernatant. We measured the infectious particle (IP)/ul titer of the supernatant by infecting A549 or CCL-141 cells with a known volume of viral supernatant, and quantifying the number of HA+ cells 9 hours post-infection. Cells were stained for HA using the H17-L19 antibody (Gerhard et al., 1981), which reacts with the WSN HA (Doud et al., 2017).

##### Competitions

Viral competition assays were performed in biological duplicate (starting from independent bacterial clones of each PB2 mutant) in both A549 and CCL-141 cells. Cells were infected with a mixture of wild-type and PB2-mutant virus at a 1:1 ratio based on the IP/ul titer as measured in the given cell type. We infected a minimum of 3.78 × 10^5^ or 7.62 × 10^5^ cells, at MOIs of 0.1 or 0.01, for the 10 h and 48 h timepoints respectively. At 2 hours post-infection, we replaced either D10 or E10 media with fresh WGM. At 10 hours post-infection, cells infected at MOI of 0.1 were lysed in buffer RLT and cellular RNA was extracted using the Qiagen RNeasy Mini Kit. At 48 hours post-infection, we collected viral supernatant from cells infected at MOI of 0.01, and extracted RNA using the QIAamp Viral RNA Mini Kit (Qiagen, 52904).

Sequencing to determine mutant frequency: We reverse transcribed the full length PB2 vRNA from the extracted RNA with SuperScript III (ThermoFisher Scientific, 18080051) (primer S009-PB2-full-1F, Table S7: Viral competition). For each PB2 mutant, we PCR amplified from the cDNA the region of PB2 centered around that mutated codon site (Table S7). This product was then subject to a second PCR using primers that add Illumina sequencing adaptors. Finally, these libraries were deep sequenced on an Illumina MiSeq using 50 bp single-end reads. Computational analyses to quantify mutant versus wild-type frequency are provided in File S2, and at https://github.com/jbloomlab/PB2-DMS.

#### H7N9 phylogenetic analysis

The phylogenetic tree was generated using Nextstrain’s augur pipeline (Hadfield et al., 2018), and ancestral state reconstruction and adjustment of branch lengths according to sequence isolation date were performed with TreeTime (Sagulenko et al., 2018).

Ancestral state reconstruction was only performed for nucleotide states, and was not used to infer ancestral host states. Instead, we inferred host-state transitions associated with each branch of the tree in a way that leveraged the prior knowledge that most H7N9 viruses circulate in avian hosts, and that most human infections arise from direct avian-to-human transmissions (Su et al., 2017). Specifically, for each node in the tree, starting from the root, we gathered all tips descending from that node. If that clade included only human sequences, and its parent node also included only human sequences, then the current clade falls within a monophyletic human clade, and the branch leading to it was labeled human-to-human. If the current clade includes only human sequences but the parent node includes non-human sequences, then the branch leading to the clade was labeled avian-to-human. If the current clade includes both human and non-human sequences, then the branch leading to the clade was labeled avian-to-avian. Mutations were classified as human if they occurred on human-to-human and avian-to-human branches, and are classified as avian if they occurred on avian-to-avian branches. Note that since H7N9 human influenza typically results from avian-to-human transmissions, the branches we label as human-to-human in reality likely arise from avian-to-human transmissions, but which we cannot accurately reconstruct due to insufficient sampling of avian sequences. For this reason, human-to-human and avian-to-human branches were grouped together. Further details are provided in Jupyter notebook at https://github.com/jbloomlab/PB2-DMS, or in File S2.

#### Accessibility of mutations

We calculated the accessibility, or mean nucleotide substitutions required to access an amino-acid mutation, from avian influenza PB2 sequences collected from 2015 through 2018. The accessibility of codon *c* to amino-acid *a* by single-nucleotide mutations is defined as the minimum number of nucleotide mutations needed to generate any codon for that amino-acid. For a collection of sequences, we calculate the accessibility as the weighted average of the accessibilities of all codons observed at that site in the collection of sequences. Accessibility was calculated using code as documented here (https://jbloomlab.github.io/dms_tools2/dms_tools2.utils.html?highlight=accessibility#d ms_tools2.utils.codonEvolAccessibility), in Jupyter notebook at https://github.com/jbloomlab/PB2-DMS, and in File S2.

### Quantification and statistical analysis

Quantification and statistical analysis was performed in Python and a complete description is available in main text, methods, associated figure legends, and computational Jupyter notebooks.

### Data and software availability

Deep sequencing data have been deposited in the NCBI Sequence Read Archive under BioProject accession number PRJNA511556.

The GitHub repository https://github.com/jbloomlab/PB2-DMS contains Jupyter notebooks that perform all steps of computational analyses and provide detailed step-by-step explanations and plots. The README file explains the organization of the notebooks and other files. HTML renderings of the notebooks are provided in File S2.

## Supplemental Information

**Figure S1. Related to Figure 1.**
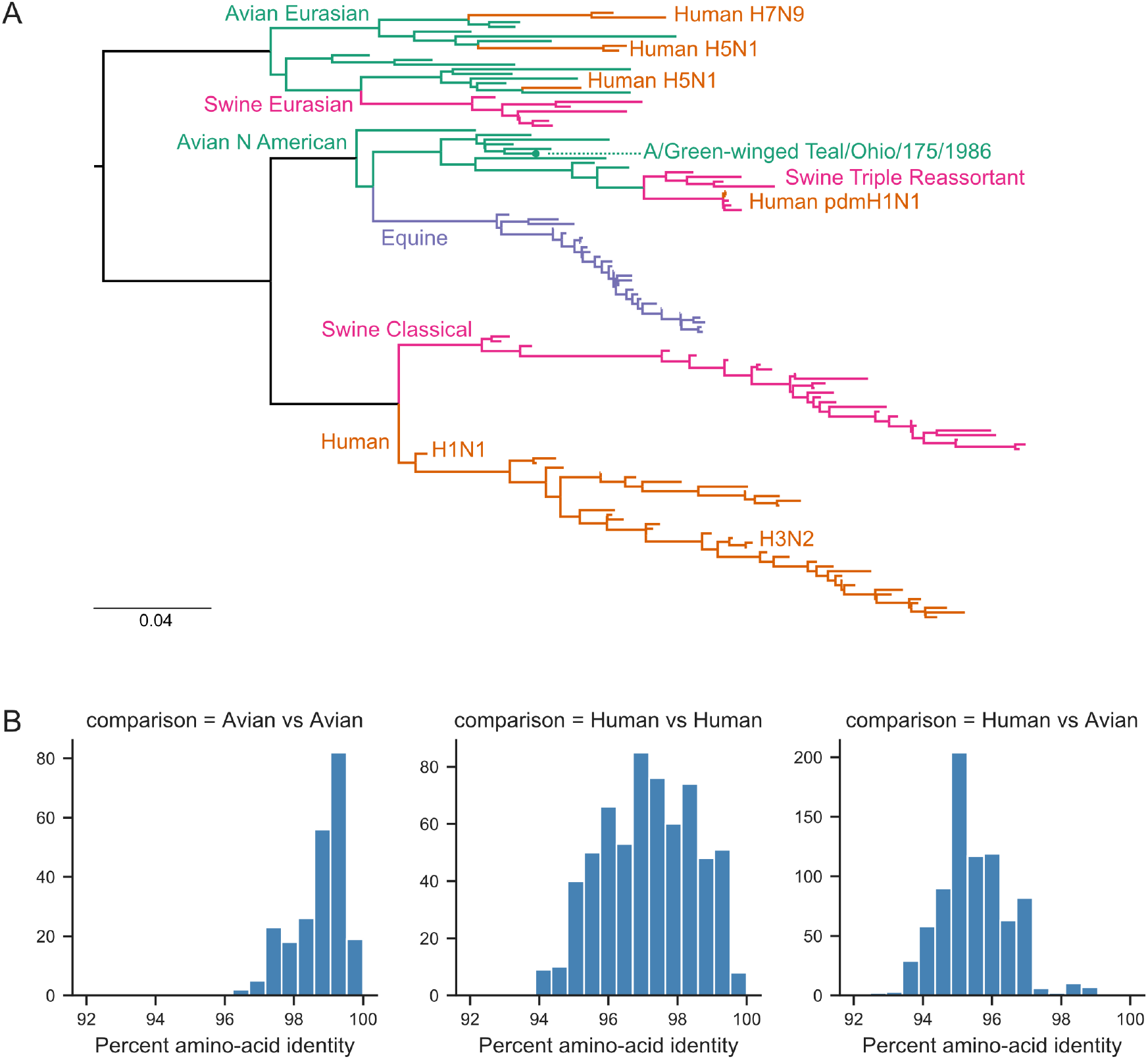
Phylogenetic relationship of PB2 sequence of chosen avian influenza strain to other influenza strains. (A) Phylogenetic tree of influenza PB2. We used PB2 sequences from the following influenza strains: A/Green-winged Teal/Ohio/175/1986 (indicated with a green dot), diverse strains sampled across years and hosts (Doud et al., 2015), and representatives of lineage-defining strains (human H3N2, human pandemic H1N1) and recent sporadic human cases of avian influenza strains (H5N1, H7N9). PB2 nucleotide sequences (of the coding sequence) were aligned using MAFFT and the phylogenetic tree was built using RAxML using the GTRCAT substitution model. Scale bar: mean nucleotide substitutions per site. (B) Pairwise amino-acid identity between all PB2 sequences shown in the tree, between just avian strains, between just human strains, and between human and avian strains.

**Figure S2. Related to Figure 1.**
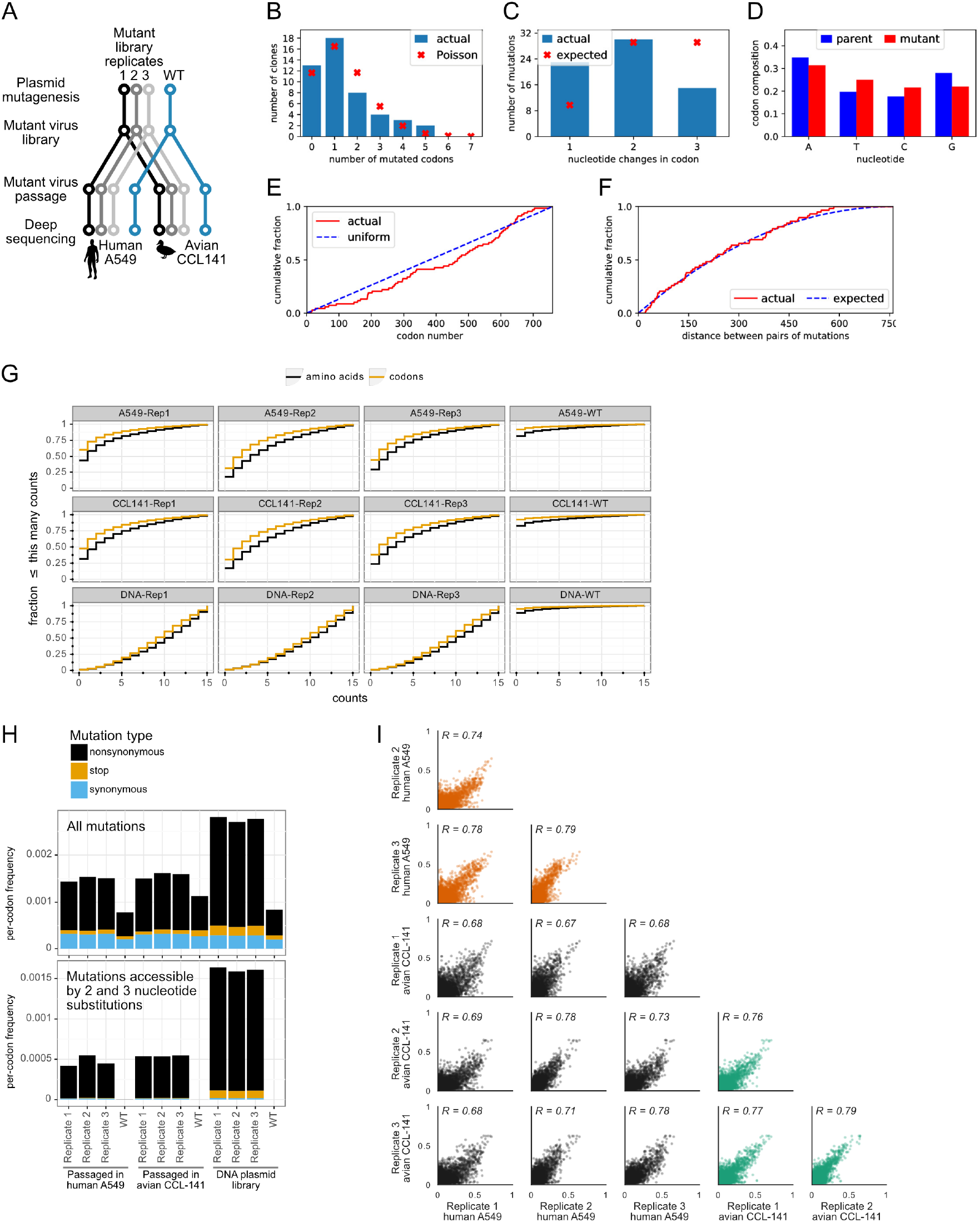
Details of deep mutational scanning experiment. (A) Experiments were performed in biological triplicate, starting from plasmid mutagenesis. All experimental steps were also performed on a wild-type PB2 gene (blue) in order to estimate error rates during deep sequencing and other experimental steps. (B—F) We picked 48 clones across the three replicate mutant plasmid libraries for Sanger sequencing. (B) There was an average of 1.4 codon mutants per clone, with the number of mutations per clone roughly following a Poisson distribution. (C) Distribution of number of nucleotide changes for each codon mutation. (D) Nucleotide frequencies in the mutant versus parent codons. (E) Mutations were distributed uniformly across the PB2 gene. (F) Cumulative distribution of pairwise distance between pairs of codon mutations, for clones with multiple mutations. The observed distribution is close to the expected distribution of pairs of mutations occurred independently. (G) Cumulative distribution of the fraction of mutations that are found less than or equal to the indicated number of times. “DNA” refers to the mutant plasmid library. (H) Per codon frequencies of nonsynonymous, stop, and synonymous mutations for each mutant library replicate and wild type, measured either in the DNA plasmid library or after passaging in human (A549) or avian (CCL-141) cells. The top plot shows all mutations; the bottom plot shows only mutations accessible by 2 and 3 nucleotide substitutions. (I) Correlations among experimental replicates of all amino-acid preferences. Correlations compare replicates passaged in human cells (orange), avian cells (green), and between the two cell types (black).

**Figure S3. Related to Figure 2.**
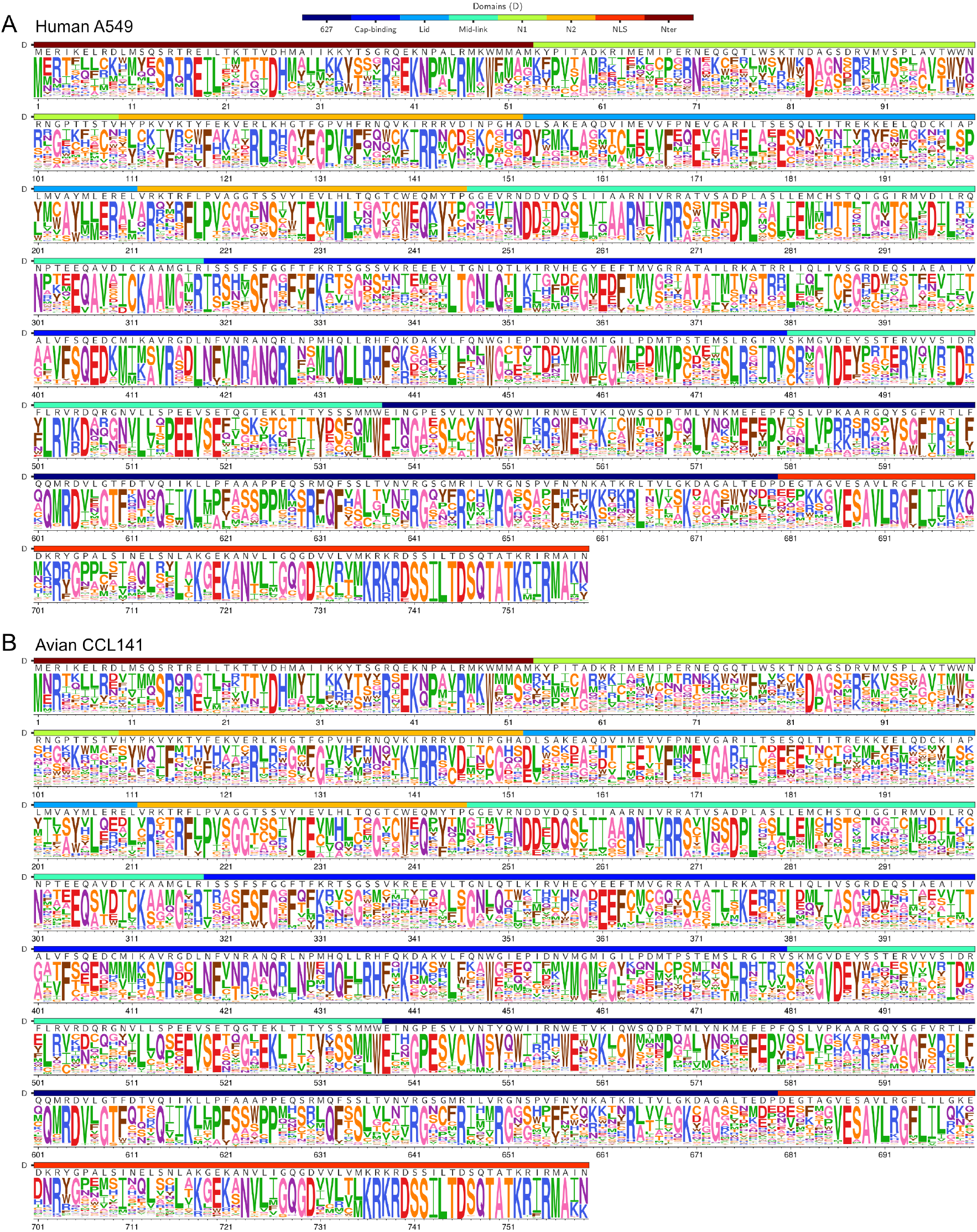
Complete map of amino acid preferences measured in human and avian cells. (A) Measurements in human (A549) cells. (B) Measurements in avian (CCL-141) cells. The height of the letter at each site is proportional to the rescaled preference for that amino acid at that site. Domains of PB2 (Pflug et al., 2017) are indicated by the top color bar. The wild-type S009 PB2 sequence is indicated by the letters above each site’s logoplot.

**Figure S4. Related to Figure 3.**
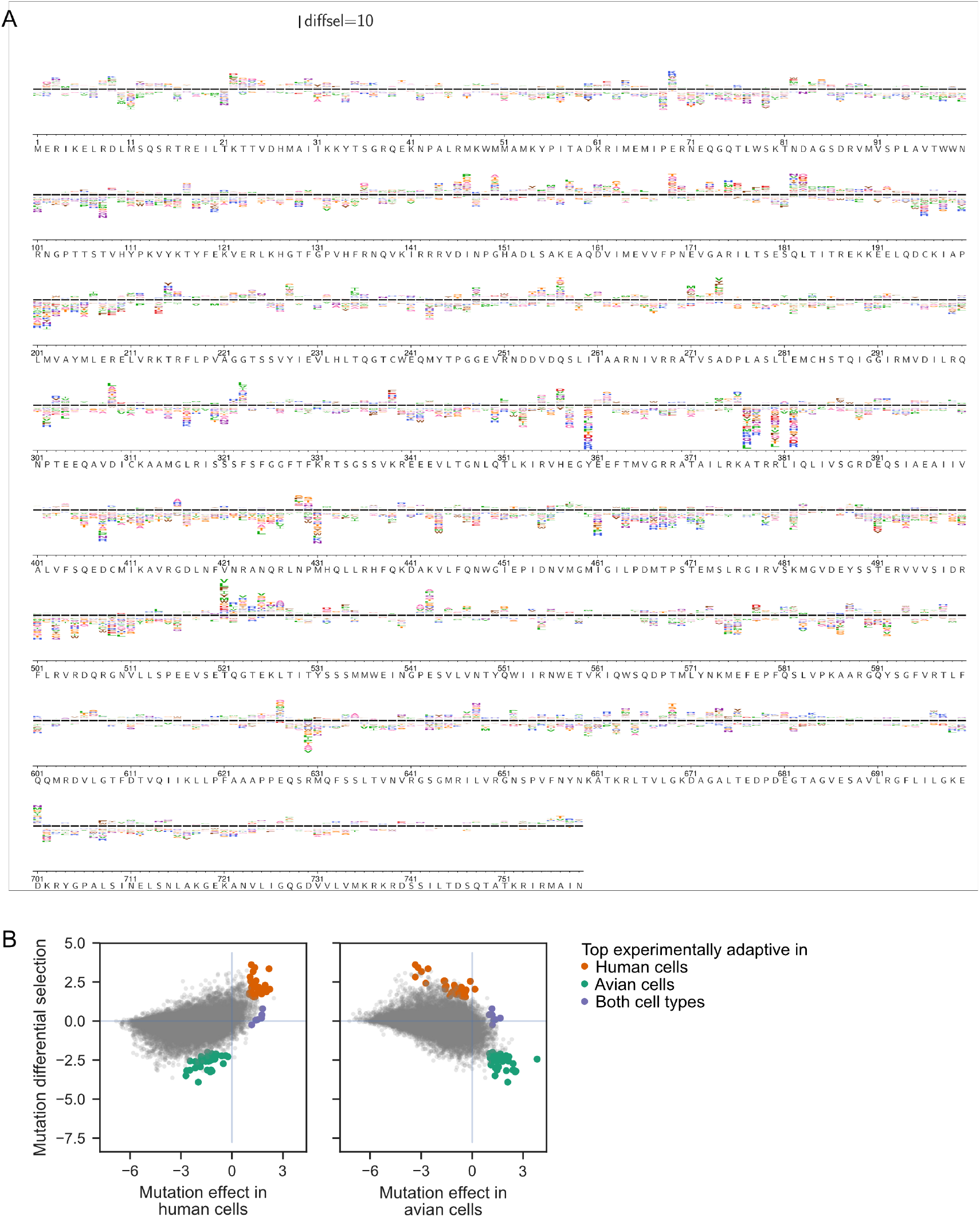
Complete map of differential selection in human versus avian cells. (A) Differential selection for PB2. The height of the letter at each site is proportional to the differential selection in human versus avian cells for that amino acid at that site. Letters above the center line are favored in human cells. The wild-type avian influenza (S009) PB2 sequence is indicated above each site’s logoplot. (B) Scatter plots of differential selection versus mutation effect as measured in human or avian cells. Top experimentally adaptive mutations identified in our deep mutational scanning (orange, green, purple dots) are indicated on the plots.

**Figure S5. Related to Figure 5.**
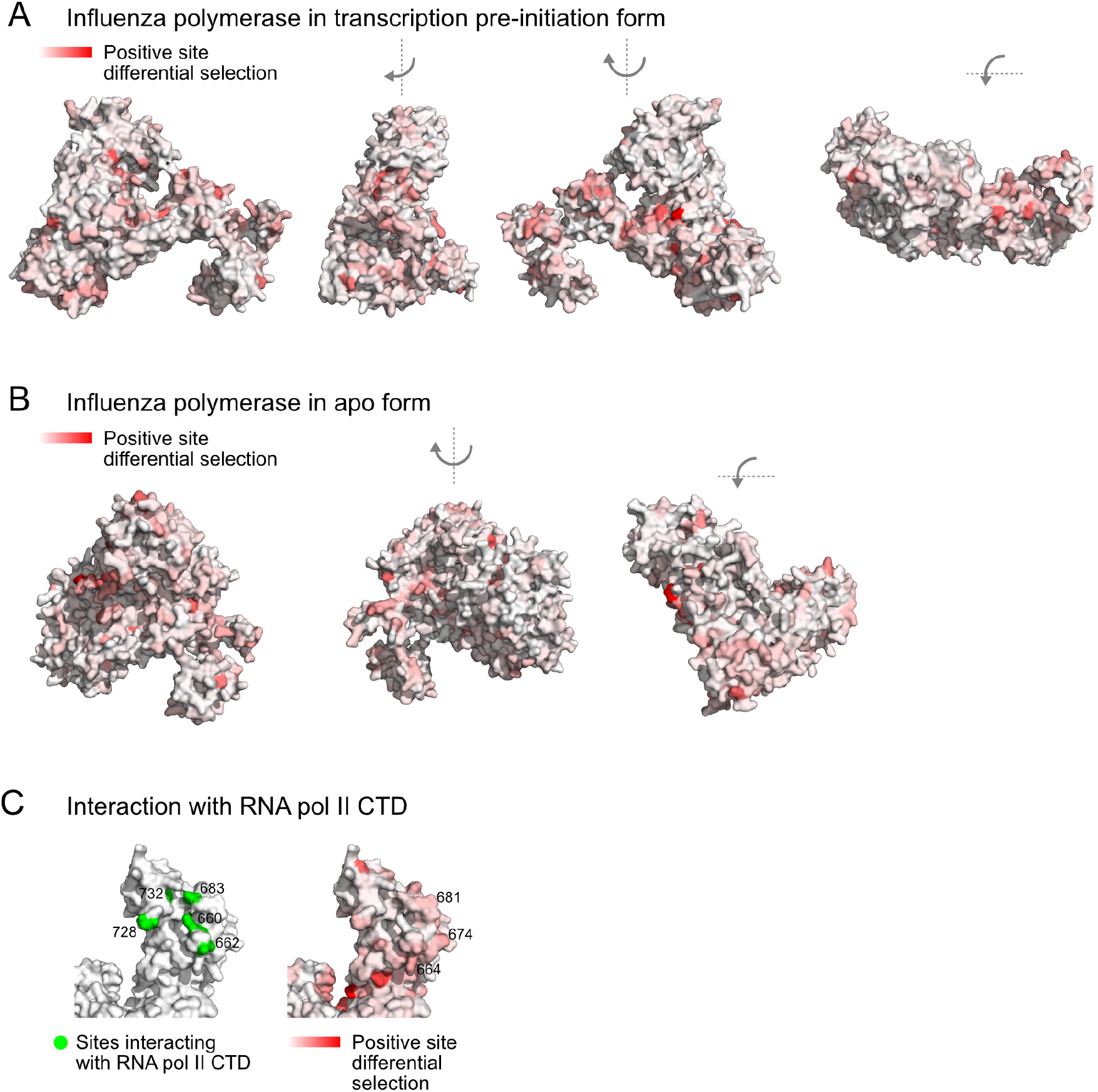
Locations of identified human adaptive mutations on influenza polymerase. Positive site differential selection mapped onto the PB2 subunit of the influenza polymerase complex in (A) the transcription pre-initiation form (PDB: 4WSB), and (B) the apo form (PDB: 5D98). (C) Positive site differential selection mapped onto PB2 (PDB: 6F5O). Sites in PB2 involved in RNA Pol II CTD binding indicated in green. Sites with high differential selection are numbered. The avian influenza (S009) PB2 amino acid sequence was mapped onto the PB2 chain by one-2-one threading using Phyre2 (Kelley et al., 2015) (Confidence in model for 6F5O: 100%). Sites are numbered according to the S009 PB2 sequence.

**Table S1.** *Related to Methods*. Comparison of ExpCM to standard phylogenetic substitution models.

| Model | deltaAIC | LogLikelihood | nParams | ParamValues |
| --- | --- | --- | --- | --- |
| ExpCM_A549avgprefs | 0 | -30634.89 | 6 | beta=2.51, kappa=8.63, omega=0.15 |
| ExpCM_CCL141avgprefs | 385.04 | -30827.41 | 6 | beta=2.57, kappa=8.58, omega=0.12 |
| YNGKP_M5 | 3507.84 | -32382.81 | 12 | alpha_omega=0.30, beta_omega=6.07, kappa=7.77 |
| averaged_ExpCM_CCL141avgprefs | 3516.34 | -32393.06 | 6 | beta=0.44, kappa=8.63, omega=0.04 |
| averaged_ExpCM_A549avgprefs | 3516.92 | -32393.35 | 6 | beta=0.33, kappa=8.64, omega=0.04 |
| YNGKP_M0 | 4131.2 | -32695.49 | 11 | kappa=7.74, omega=0.04 |

*Table S2. Related to Figure 2 and 3*. Preference, mutation effect, and differential selection results for all mutations.

*Table S3. Related to Figure 3*. Catalog of previously described human/mammalian adaptive mutations.

*Table S4. Related to Figure 4*. Flow cytometry data for minigenome assays.

*Table S5. Related to Figure 4*. Mutant frequency data for competition assay.

*Table S6. Related to Figure 6*. H7N9 human and avian mutation counts.

*Table S7. Related to Methods*. Primers sequences.

*File S1. Related to Methods*. Plasmid sequences.

*File S2. Related to Methods*. Jupyter notebooks documenting computational analyses.

