## Supplementary material for "Comprehensive mapping of avian influenza polymerase adaptation to the human host": File S2: Fig1_ProcessDMSdata.html


### Align deep sequencing data and count mutations¶

#### Import modules, define directories¶

In [14]:

```
import os
import re
import glob
import shutil
import subprocess
import string
import warnings
warnings.simplefilter('ignore') # don't print warnings to avoid clutter in notebook

import pandas as pd
import numpy as np
import matplotlib as mpl
import matplotlib.pyplot as plt
%matplotlib inline
# import seaborn as sns
# from Bio import SeqIO, AlignIO

import dms_tools2
import dms_tools2.sra
import dms_tools2.plot
from dms_tools2.plot import plotReadsPerBC
import dms_tools2.prefs
from dms_tools2.ipython_utils import showPDF

from IPython.display import display, HTML, Markdown, Image

from pymodules.utils import annotateCodonCounts_accessible 
from pymodules.utils import plotCodonMutTypes_accessible

print("Using dms_tools2 version {0}".format(dms_tools2.__version__))

# CPUs to use, should not exceed the number you request with slurm
# ncpus = -1
ncpus = 14

# do we use existing results or generate everything new?
use_existing = 'yes'

# Directories
fastqdir = './fastq/'
datadir = './data'
resultsdir = './results/' 
countsdir = os.path.join(resultsdir, 'codoncounts/')
paperdir = './paper'
figuresdir = os.path.join(paperdir, 'figures/')

for xdir in [fastqdir, datadir, resultsdir, countsdir, paperdir, figuresdir]:
    if not os.path.isdir(xdir):
        os.mkdir(xdir)
```

```
Using dms_tools2 version 2.3.0
```

#### Experimental set-up¶

I performed a full-scale passaging of all three replicate mutant libraries (named Lib4, 5, 6) and wild-type virus in human A549 and duck CCL141 cells. Virus supernatant was collected 48hpi.

Details of pre-selection samples and wild-type controls sequenced as follows:

| Plasmid | Bottleneck # molecules |
| --- | --- |
| DNA-Lib4/Rep1 | 750,000 |
| DNA-Lib5/Rep2 | 750,000 |
| DNA-Lib6/Rep3 | 750,000 |
| DNA-WT | 250,000 |

Details of post-selection samples and wild-type controls sequenced as follows:

| Cells | Virus | MOI | # TCID50s passaged | Timepoint sequenced | Bottleneck # molecules |
| --- | --- | --- | --- | --- | --- |
| A549 | Lib4/Rep1 | 0.01 | 9.39 x 10^5^ | 48hpi | 750,000 |
| A549 | Lib5/Rep2 | 0.01 | 1.02 x 10^6^ | 48hpi | 750,000 |
| A549 | Lib6/Rep3 | 0.01 | 1.20 x 10^6^ | 48hpi | 750,000 |
| A549 | WT | 0.01 | 9.30 x 10^5^ | 48hpi | 250,000 |
| CCL141 | Lib4/Rep1 | 0.01 | 8.46 x 10^5^ | 48hpi | 750,000 |
| CCL141 | Lib5/Rep2 | 0.01 | 8.92 x 10^5^ | 48hpi | 750,000 |
| CCL141 | Lib6/Rep3 | 0.01 | 9.54 x 10^5^ | 48hpi | 750,000 |
| CCL141 | WT | 0.01 | 8.58 x 10^5^ | 48hpi | 250,000 |

#### Download FASTQ files from the SRA¶

The deep mutational scanning FASTQ files are deposited on the Sequence Read Archive under the run numbers provided in the file `data/SraRunTable_DMS.txt`. To download these files, we use the dms\_tools2.sra.fastqFromSRA function. This requires the fastq-dump and aspera programs to be installed at the specified paths.

In [15]:

```
samples = (pd.read_table('data/SraRunTable_DMS.txt', sep='\t')[['Sample_Name', 'Run']]
           .sort_values('Sample_Name')
           .rename(columns={'Sample_Name': 'name', 'Run': 'run'})
          )
display(HTML(samples.to_html(index=False)))
```

| name | run |
| --- | --- |
| A549-MutVirusLib-Rep1 | SRR8366384 |
| A549-MutVirusLib-Rep2 | SRR8366383 |
| A549-MutVirusLib-Rep3 | SRR8366382 |
| A549-WT | SRR8366381 |
| CCL141-MutVirusLib-Rep1 | SRR8366380 |
| CCL141-MutVirusLib-Rep2 | SRR8366379 |
| CCL141-MutVirusLib-Rep3 | SRR8366378 |
| CCL141-WT | SRR8366377 |
| DNA-Lib-Rep1 | SRR8366386 |
| DNA-Lib-Rep2 | SRR8366385 |
| DNA-Lib-Rep3 | SRR8366376 |
| DNA-WT | SRR8366375 |

In [16]:

```
print('Downloading FASTQ files from the SRA...')
dms_tools2.sra.fastqFromSRA(
        samples=samples,
        fastq_dump='fastq-dump', # valid path to this program on the Hutch server
        fastqdir=fastqdir,
        aspera=(
            '/app/aspera-connect/3.5.1/bin/ascp', # valid path to ascp on Hutch server
            '/app/aspera-connect/3.5.1/etc/asperaweb_id_dsa.openssh' # Aspera key on Hutch server
            ),
        overwrite={'no':True, 'yes':False}[use_existing],
        )
print('Completed download of FASTQ files from the SRA')

print('Here are the names of the downloaded files now found in {0}'.format(fastqdir))
display(HTML(samples.to_html(index=False)))
```

```
Downloading FASTQ files from the SRA...
Completed download of FASTQ files from the SRA
Here are the names of the downloaded files now found in ./fastq/
```

| name | run | R1 | R2 |
| --- | --- | --- | --- |
| A549-MutVirusLib-Rep1 | SRR8366384 | A549-MutVirusLib-Rep1\_R1.fastq.gz | A549-MutVirusLib-Rep1\_R2.fastq.gz |
| A549-MutVirusLib-Rep2 | SRR8366383 | A549-MutVirusLib-Rep2\_R1.fastq.gz | A549-MutVirusLib-Rep2\_R2.fastq.gz |
| A549-MutVirusLib-Rep3 | SRR8366382 | A549-MutVirusLib-Rep3\_R1.fastq.gz | A549-MutVirusLib-Rep3\_R2.fastq.gz |
| A549-WT | SRR8366381 | A549-WT\_R1.fastq.gz | A549-WT\_R2.fastq.gz |
| CCL141-MutVirusLib-Rep1 | SRR8366380 | CCL141-MutVirusLib-Rep1\_R1.fastq.gz | CCL141-MutVirusLib-Rep1\_R2.fastq.gz |
| CCL141-MutVirusLib-Rep2 | SRR8366379 | CCL141-MutVirusLib-Rep2\_R1.fastq.gz | CCL141-MutVirusLib-Rep2\_R2.fastq.gz |
| CCL141-MutVirusLib-Rep3 | SRR8366378 | CCL141-MutVirusLib-Rep3\_R1.fastq.gz | CCL141-MutVirusLib-Rep3\_R2.fastq.gz |
| CCL141-WT | SRR8366377 | CCL141-WT\_R1.fastq.gz | CCL141-WT\_R2.fastq.gz |
| DNA-Lib-Rep1 | SRR8366386 | DNA-Lib-Rep1\_R1.fastq.gz | DNA-Lib-Rep1\_R2.fastq.gz |
| DNA-Lib-Rep2 | SRR8366385 | DNA-Lib-Rep2\_R1.fastq.gz | DNA-Lib-Rep2\_R2.fastq.gz |
| DNA-Lib-Rep3 | SRR8366376 | DNA-Lib-Rep3\_R1.fastq.gz | DNA-Lib-Rep3\_R2.fastq.gz |
| DNA-WT | SRR8366375 | DNA-WT\_R1.fastq.gz | DNA-WT\_R2.fastq.gz |

In [17]:

```
# Match library number to replicate number
samples_lib = (pd.read_table('data/SraRunTable_DMS.txt', sep='\t')[['Sample_Name', 'Library_Name']]
           .sort_values('Sample_Name')
           .rename(columns={'Sample_Name': 'name'})
          )
samples = pd.merge(samples_lib, samples, on='name')
# write sample information to a batch file for dms2_batch_bcsubamplicons
countsbatchfile = os.path.join(countsdir, 'batch.csv')
samples = (samples[['Library_Name', 'R1']]
                 .rename(columns={'Library_Name': 'name'})
                )
print("Here is the batch file that we write to CSV format to use as input to dms2_batch_bcsubamplicons:")
display(HTML(samples.to_html(index=False)))
samples.to_csv(countsbatchfile, index=False)
```

```
Here is the batch file that we write to CSV format to use as input to dms2_batch_bcsubamplicons:
```

| name | R1 |
| --- | --- |
| A549-Lib4 | A549-MutVirusLib-Rep1\_R1.fastq.gz |
| A549-Lib5 | A549-MutVirusLib-Rep2\_R1.fastq.gz |
| A549-Lib6 | A549-MutVirusLib-Rep3\_R1.fastq.gz |
| A549-WT | A549-WT\_R1.fastq.gz |
| CCL141-Lib4 | CCL141-MutVirusLib-Rep1\_R1.fastq.gz |
| CCL141-Lib5 | CCL141-MutVirusLib-Rep2\_R1.fastq.gz |
| CCL141-Lib6 | CCL141-MutVirusLib-Rep3\_R1.fastq.gz |
| CCL141-WT | CCL141-WT\_R1.fastq.gz |
| DNA-Lib4 | DNA-Lib-Rep1\_R1.fastq.gz |
| DNA-Lib5 | DNA-Lib-Rep2\_R1.fastq.gz |
| DNA-Lib6 | DNA-Lib-Rep3\_R1.fastq.gz |
| DNA-WT | DNA-WT\_R1.fastq.gz |

#### Barcoded subamplicon alignment¶

Use dms2\_batch\_bcsubamp to align sequences to the wildtype S009 PB2 coding sequence and count mutations.

In [18]:

```
# Reference coding sequence of S009 PB2
refseq = './data/S009-PB2.fasta'

alignspecs = ' '.join(['1,285,36,32', 
                       '286,570,32,35', 
                       '571,855,37,32', 
                       '856,1140,32,35', 
                       '1141,1425,28,34', 
                       '1426,1710,33,34',
                       '1711,1995,29,29',
                       '1996,2280,28,43'])

print('\nNow running:')
cmd = 'dms2_batch_bcsubamp \
    --batchfile {0} \
    --refseq {1} \
    --alignspecs {2} \
    --outdir {3} \
    --summaryprefix summary \
    --R1trim 200 \
    --R2trim 170 \
    --fastqdir {4} \
    --ncpus {5} \
    --use_existing {6}'\
    .format(countsbatchfile, refseq, alignspecs, countsdir, fastqdir, ncpus, use_existing)
print(cmd)
!$cmd
```

```
Now running:
dms2_batch_bcsubamp     --batchfile ./results/codoncounts/batch.csv     --refseq ./data/S009-PB2.fasta     --alignspecs 1,285,36,32 286,570,32,35 571,855,37,32 856,1140,32,35 1141,1425,28,34 1426,1710,33,34 1711,1995,29,29 1996,2280,28,43     --outdir ./results/codoncounts/     --summaryprefix summary     --R1trim 200     --R2trim 170     --fastqdir ./fastq/     --ncpus 14     --use_existing yes
Output summary files already exist and '--use_existing' is 'yes', so exiting with no further action.
```

Output counts of each codon at each site in each sample:

In [19]:

```
!ls {countsdir}/*_codoncounts.csv
```

```
./results/codoncounts//A549-Lib4_codoncounts.csv
./results/codoncounts//A549-Lib5_codoncounts.csv
./results/codoncounts//A549-Lib6_codoncounts.csv
./results/codoncounts//A549-WT_codoncounts.csv
./results/codoncounts//CCL141-Lib4_codoncounts.csv
./results/codoncounts//CCL141-Lib5_codoncounts.csv
./results/codoncounts//CCL141-Lib6_codoncounts.csv
./results/codoncounts//CCL141-WT_codoncounts.csv
./results/codoncounts//DNA-Lib4_codoncounts.csv
./results/codoncounts//DNA-Lib5_codoncounts.csv
./results/codoncounts//DNA-Lib6_codoncounts.csv
./results/codoncounts//DNA-WT_codoncounts.csv
```

#### Plots summarizing read alignments¶

Now we look at the summary plots created by dms2\_batch\_bcsubamp.

In [20]:

```
countsplotprefix = os.path.join(countsdir, 'summary')
countsplotprefix
```

Out[20]:

```
'./results/codoncounts/summary'
```

##### Number of aligned reads and barcodes¶

Below are the number of aligned reads and barcodes for each sample.

The `summary_readstats.pdf` plot below shows that most reads were retained, and a small fraction discarded because of low-quality barcodes, or because they failed the Illumina filter.

The `summary_bcstats.pdf` plot below shows statistics on the barcodes. Some of the barcodes had to be discarded because they had too few reads (these are the single-read barcodes in the plot below), a small fraction with adequate reads were not alignable, and the rest aligned to the PB2 gene properly.

I sequenced the WT controls to about 1/3 depth, so the number of reads are lower as expected.

In [21]:

```
showPDF([countsplotprefix + '_readstats.pdf', countsplotprefix + '_bcstats.pdf'])
```

The `*_readsperbc.pdf` plot below shows how many times different barcodes were observed for each sample.
Barcodes need to be observed multiple times to be useful for barcoded-subamplicon sequencing error correction.
Many barcodes are observed multiple times.
There is also a large bar at 1, which probably corresponds to a mix of barcodes that were only observed once and sequencing errors on other barcodes that spuriously gave rise to an apparently unique sequence.
Overall, the distributions here indicate that most multiply sequenced barcodes were sequenced > 2 times, which suggests that most of the peak at one is probably due to sequencing errors on barcodes, and suggests that sequencing to greater depth probably would not help all that much for these samples.

In [22]:

```
# showPDF(countsplotprefix + '_readsperbc.pdf')
# Made an identical plot to the one named aboe, but below with a different layout (maxcol=4) for easier reading.
names = samples['name']
readsperbcfiles = [countsdir+'/'+name+'_readsperbc.csv' for name in names]
plotReadsPerBC(names, readsperbcfiles, countsplotprefix + '_readsperbc2.pdf', maxreads=10, maxcol=4)
showPDF(countsplotprefix + '_readsperbc2.pdf')
```

##### Sequencing depth and mutation frequencies across sequences¶

The `summary_depth.pdf` plot below shows the depth (number of called codons) at each site in the gene, after restricting to barcodes for which there are enough reads. Most sites in most samples are sequenced to >400,000 codon counts. I sequenced the WT controls to about 1/3 depth, so the number of counts are lower as expected.

In [23]:

```
showPDF(countsplotprefix + '_depth.pdf')
```

The `summary_mutfreq.pdf` plot below shows the per-codon frequency of mutations at each site.
Generally, things are about as expected. In the mutant DNA library samples, there is a fairly high mutation frequency across all sites. In the post-selection mutant viruses, mutations remain prevalent, though at lower levels, suggesting that there is some selection against mutations. In the wildtype plasmid and wildtype virus, the mutation frequency is much lower.

I observe a prominent peak in all the DNA mutant libraries at around position 200, that seems to remain in the passaged virus samples. This is present in all replicate libraries. This is likely an issue with the process of mutagenesis:

- I had made my replicate libraries from three independent clones of the plasmid encoding WT PB2, so it is unlikely that three independent clones had issues at site ~200.
- The WT virus was rescued with plasmid that generated mutant Lib 4. WT virus does not show a peak at 200.
- The DNA-WT control was from plasmid that generated mutant lib 6. DNA-WT also does not show a peak at 200.

This peak should be accounted for during data analysis since we will compare passaged samples to the DNA library samples.

In [24]:

```
showPDF(countsplotprefix + '_mutfreq.pdf')
```

##### Completeness of mutation sampling¶

The `summary_cumulmutcounts.pdf` plot below shows the fraction of mutations that are found $\leq$ the indicated number of times.
This gives an idea of how completely the possible amino-acid and codon mutations were sampled in each sample.

In the plasmid mutant library, nearly all codon and amino-acid mutations were sampled at least once (and usually much more than once), indicating that my plasmid mutant libraries contained virtually all these mutations as desired.

For mutant libraries passaged in A549 and CCL141, many but not all amino acid mutations were sampled at least once. This suggests that mutations are being lost from sampling, consistent with some mutations being lost through selection.

The wildtype controls sampled only a relatively small fraction of the possible mutations, consistent with mutations/errors in passaging and sequencing being rare.

In [25]:

```
showPDF(countsplotprefix + '_cumulmutcounts.pdf')
```

##### Average mutation frequencies¶

The `summary_codonntchanges.pdf` plot shows per-codon frequency across the entire gene of codon mutations that change different numbers of nucleotides (e.g., `ATG` to `AAG` changes 1 nucleotide, `ATG` to `AAC` changes 2 nucleotides, and `ATG` to `CAC` changes 3 nucleotides).
Since we are dealing with codon-mutant libraries in this experiments, we see all three types of mutations in the mutagenized plasmid DNA and virus samples.
The "mutations" in the wildtype controls are sequencing errors, and those are almost all single-nucleotide changes since it is very rare for two nucleotides in the same codon to both experience a sequencing error.

The `summary_codonmuttypes.pdf` plot shows the average per-codon frequency of nonsynonymous, synonymous, and stop codon mutations across the entire gene.
As expected, we see some selection against stop codons and moderate selection against nonsynonymous mutations in the post-selection samples relative to plasmid DNA samples, indicating purifying selection against deleterious changes.

In [26]:

```
showPDF([countsplotprefix + '_codonntchanges.pdf', countsplotprefix + '_codonmuttypes.pdf'])
```

In [27]:

```
codonmuttypes = pd.read_csv(countsplotprefix + '_codonmuttypes.csv').sort_index(axis=1)
display(HTML(codonmuttypes.to_html(index=False, float_format=lambda x: '{0:.3g}'.format(x))))
```

| name | nonsynonymous | stop | synonymous |
| --- | --- | --- | --- |
| A549-Lib4 | 0.00103 | 7.86e-05 | 0.000329 |
| A549-Lib5 | 0.00114 | 7.63e-05 | 0.000315 |
| A549-Lib6 | 0.00109 | 8.92e-05 | 0.000329 |
| A549-WT | 0.000507 | 7.46e-05 | 0.000202 |
| CCL141-Lib4 | 0.00112 | 6.46e-05 | 0.000315 |
| CCL141-Lib5 | 0.0012 | 9.08e-05 | 0.00033 |
| CCL141-Lib6 | 0.00119 | 8.33e-05 | 0.000326 |
| CCL141-WT | 0.000724 | 0.000128 | 0.000276 |
| DNA-Lib4 | 0.00231 | 0.0002 | 0.0003 |
| DNA-Lib5 | 0.00223 | 0.000187 | 0.000287 |
| DNA-Lib6 | 0.00227 | 0.0002 | 0.000295 |
| DNA-WT | 0.000544 | 0.000103 | 0.000194 |

Now I calculate the corrected mutation average mutation frequencies by subtracting the appropriate error control for each sample:

- The pre-selection frequency is mutDNA - wtDNA
- The post-selection frequency is mutvirus - wtvirus

When I use the respective WT controls, I find 2-15% stop codons remaining in A549. In CCL141 cells, there is a very high rate of stop mutations in the wtvirus control, so there is <0% stop codons remaining.

Because the wtvirus control passaged in CCL141 appears to have rather high error rate, I will also try using DNA-WT as the error control:

- The pre-selection frequency (2) is mutDNA - wtDNA
- The post-selection frequency (2) is mutvirus - wtDNA

When I use wtDNA as the error control, the both A549 and CCL141 have <0% stop codons remaining, as the mutation rate in the DNA-WT control is quite high.

I suspect that I am seeing high rate of mutations due to oxidative damage. To eliminate the effect of mutations due to oxidative damage, I can make these estimates considering only 2 and 3 nucleotide mutations (see below).

In [28]:

```
mutvirus = (pd.concat([codonmuttypes[codonmuttypes['name'].str.contains('A549-Lib')], 
                     codonmuttypes[codonmuttypes['name'].str.contains('CCL141-Lib')]
                     ])
            .assign(experiment=lambda x: x.name,
                  sample='mutvirus'))
wtvirus = (pd.concat(
    [codonmuttypes[codonmuttypes['name'].str.contains('A549-WT')]\
    .assign(experiment='A549-{0}'.format(i),
           sample='wtvirus')
    for i in ['Lib4','Lib5','Lib6']
    ] +\
    [codonmuttypes[codonmuttypes['name'].str.contains('CCL141-WT')]\
    .assign(experiment='CCL141-{0}'.format(i),
           sample='wtvirus')
    for i in ['Lib4','Lib5','Lib6']
    ]))
mutDNA = (pd.concat(
    [codonmuttypes[codonmuttypes['name'].str.contains('DNA-Lib')]\
    .assign(experiment=lambda x:x.name.str.replace('DNA', i),
            sample='mutDNA')
     for i in ['A549', 'CCL141']]))
wtDNA = (pd.concat(
    [codonmuttypes[codonmuttypes['name'].str.contains('DNA-WT')]\
    .assign(experiment='{0}-{1}'.format(cell, i),
            sample='wtDNA')
    for cell in ['A549', 'CCL141']
    for i in ['Lib4','Lib5','Lib6']]))

codonmuttypes_corr = (pd.concat([wtDNA, mutDNA, wtvirus, mutvirus])
                      .rename(columns={'nonsynonymous':'nonsyn', 'synonymous':'syn'})
                      .melt(id_vars=['experiment', 'sample'],
                            value_vars=['stop', 'nonsyn', 'syn'],
                            var_name='muttype',
                            value_name='mutfreq')
                      .pivot_table(index=['experiment', 'muttype'], columns='sample', values='mutfreq')
                      .assign(pre=lambda x: x.mutDNA - x.wtDNA,
                              post=lambda x: x.mutvirus - x.wtvirus)
                      .assign(percent=lambda x: 100 * x.post / x.pre)
                      .assign(pre2=lambda x: x.mutDNA - x.wtDNA,
                              post2=lambda x: x.mutvirus - x.wtDNA)
                      .assign(percent2=lambda x: 100 * x.post2 / x.pre2)
                     )
display(HTML(codonmuttypes_corr.to_html(float_format=lambda x: '{0:.3g}'.format(x))))
```

|  | sample | mutDNA | mutvirus | wtDNA | wtvirus | pre | post | percent | pre2 | post2 | percent2 |
| --- | --- | --- | --- | --- | --- | --- | --- | --- | --- | --- | --- |
| experiment | muttype |  |  |  |  |  |  |  |  |  |  |
| A549-Lib4 | nonsyn | 0.00231 | 0.00103 | 0.000544 | 0.000507 | 0.00176 | 0.000519 | 29.5 | 0.00176 | 0.000483 | 27.4 |
|  | stop | 0.0002 | 7.86e-05 | 0.000103 | 7.46e-05 | 9.72e-05 | 3.96e-06 | 4.08 | 9.72e-05 | -2.47e-05 | -25.5 |
|  | syn | 0.0003 | 0.000329 | 0.000194 | 0.000202 | 0.000106 | 0.000126 | 119 | 0.000106 | 0.000135 | 128 |
| A549-Lib5 | nonsyn | 0.00223 | 0.00114 | 0.000544 | 0.000507 | 0.00168 | 0.000636 | 37.7 | 0.00168 | 0.000599 | 35.6 |
|  | stop | 0.000187 | 7.63e-05 | 0.000103 | 7.46e-05 | 8.38e-05 | 1.68e-06 | 2 | 8.38e-05 | -2.7e-05 | -32.2 |
|  | syn | 0.000287 | 0.000315 | 0.000194 | 0.000202 | 9.36e-05 | 0.000112 | 120 | 9.36e-05 | 0.000121 | 129 |
| A549-Lib6 | nonsyn | 0.00227 | 0.00109 | 0.000544 | 0.000507 | 0.00173 | 0.000581 | 33.6 | 0.00173 | 0.000544 | 31.5 |
|  | stop | 0.0002 | 8.92e-05 | 0.000103 | 7.46e-05 | 9.65e-05 | 1.46e-05 | 15.1 | 9.65e-05 | -1.41e-05 | -14.6 |
|  | syn | 0.000295 | 0.000329 | 0.000194 | 0.000202 | 0.000102 | 0.000127 | 125 | 0.000102 | 0.000136 | 133 |
| CCL141-Lib4 | nonsyn | 0.00231 | 0.00112 | 0.000544 | 0.000724 | 0.00176 | 0.000397 | 22.5 | 0.00176 | 0.000577 | 32.7 |
|  | stop | 0.0002 | 6.46e-05 | 0.000103 | 0.000128 | 9.72e-05 | -6.32e-05 | -65.1 | 9.72e-05 | -3.87e-05 | -39.8 |
|  | syn | 0.0003 | 0.000315 | 0.000194 | 0.000276 | 0.000106 | 3.84e-05 | 36.2 | 0.000106 | 0.000121 | 114 |
| CCL141-Lib5 | nonsyn | 0.00223 | 0.0012 | 0.000544 | 0.000724 | 0.00168 | 0.000472 | 28 | 0.00168 | 0.000652 | 38.7 |
|  | stop | 0.000187 | 9.08e-05 | 0.000103 | 0.000128 | 8.38e-05 | -3.7e-05 | -44.2 | 8.38e-05 | -1.25e-05 | -14.9 |
|  | syn | 0.000287 | 0.00033 | 0.000194 | 0.000276 | 9.36e-05 | 5.33e-05 | 56.9 | 9.36e-05 | 0.000136 | 145 |
| CCL141-Lib6 | nonsyn | 0.00227 | 0.00119 | 0.000544 | 0.000724 | 0.00173 | 0.000462 | 26.7 | 0.00173 | 0.000642 | 37.1 |
|  | stop | 0.0002 | 8.33e-05 | 0.000103 | 0.000128 | 9.65e-05 | -4.45e-05 | -46.2 | 9.65e-05 | -2e-05 | -20.7 |
|  | syn | 0.000295 | 0.000326 | 0.000194 | 0.000276 | 0.000102 | 4.97e-05 | 48.9 | 0.000102 | 0.000132 | 130 |

##### Check for oxidative damage¶

The `summary_singlentchanges.pdf` plot below shows the frequency of each type of nucleotide change among only codon mutations with **one** nucleotide change. This plot is mostly useful to check if there is a large bias in which mutations appear.

Oxidative damage is characterized by an excess of C -> A ot G -> T mutations. In particular, if you are getting oxidative damage (which causes G to T mutations) during the library preparation process, you will see a large excess of C to A or G to T mutations (or both). For instance, in the case of influenza, when we get bad oxidative damage, then we see an excess of C to A mutations in the final cDNA since the damage is occurring to a ssRNA genome. If you are sequencing something without polarity, you might see both types of mutations. I see quite significant excess of these mutations, particularly in the CCL141-WT and even in the DNA-WT control. I think this should still be able to be corrected for during analysis.

In [29]:

```
showPDF(countsplotprefix + '_singlentchanges.pdf', width=600)
```

##### Investigate and isolate the effect of oxidative damage on average mutation frequencies¶

The above plot shows my data exhibit quite significant oxidative damage. To see if my high retention of stop codons is due to high oxidative damage, I subsetted mutations on ones which were accessible by one nucleotide mutations or not, and repeated plotting mutation types. To do so, I modified plotCodonMutTypes. The script that does this, `plotCodonMutTypes_accessible`, is imported from `pymodules.utils`.

In [ ]:

```
samples['countsfiles'] = samples.apply(lambda row: './results/codoncounts/{0}_codoncounts.csv'.format(row['name']), axis=1)
accessibilities = [(True, 'accessible'), (False, 'notaccessible')]
classifications = [('n_ntchanges', 'codonntchanges'), 
                   ('aachange', 'codonmuttypes')]
for accessibility, isaccessible in accessibilities:
    for classificationtype, classificationname in classifications:
        print("Annotating and plotting mutation type: {0}, {1}".format(isaccessible, classificationname))
        plotCodonMutTypes_accessible(samples['name'], samples['countsfiles'], 
                                     '{0}_{1}_{2}.pdf'.format(countsplotprefix, classificationname, isaccessible), 
                                     accessibility, classification=classificationtype, 
                                     csvfile='{0}_{1}_{2}.csv'.format(countsplotprefix, classificationname, isaccessible))
```

When I subset on non-accessible mutations (2 or 3 nucleotide substitutions), I observe much better purging of stop codons (2-7% remaining). Thus, if we eliminate spurious stop codons remaining due to oxidative damage/sequencing error, we actually get observe good selection.

In [31]:

```
for accl, acc in [('All mutations', ''), 
                  ('Accessible by 1-nucleotide substitution', '_accessible'), 
                  ('Not accessible by 1-nucleotide substitution', '_notaccessible')]:
    print(accl)
    showPDF(['{0}_{1}{2}.pdf'.format(countsplotprefix, 'codonntchanges', acc),
             '{0}_{1}{2}.pdf'.format(countsplotprefix, 'codonmuttypes', acc)])
```

```
All mutations
```

```
Accessible by 1-nucleotide substitution
```

```
Not accessible by 1-nucleotide substitution
```

In [32]:

```
codonmuttypes = pd.read_csv(countsplotprefix + '_codonmuttypes_notaccessible.csv').sort_index(axis=1)
display(HTML(codonmuttypes.to_html(index=False, float_format=lambda x: '{0:.3g}'.format(x))))
```

| name | nonsynonymous | stop | synonymous |
| --- | --- | --- | --- |
| A549-Lib4 | 0.000405 | 2.44e-06 | 1.16e-05 |
| A549-Lib5 | 0.000526 | 6.77e-06 | 1.43e-05 |
| A549-Lib6 | 0.000431 | 4.01e-06 | 1.32e-05 |
| A549-WT | 2.45e-06 | 2.59e-08 | 2.59e-08 |
| CCL141-Lib4 | 0.000523 | 3.8e-06 | 1.23e-05 |
| CCL141-Lib5 | 0.000517 | 6.26e-06 | 1.35e-05 |
| CCL141-Lib6 | 0.000528 | 4.95e-06 | 1.31e-05 |
| CCL141-WT | 6.06e-07 | 2.75e-08 | 0 |
| DNA-Lib4 | 0.00153 | 9.57e-05 | 1.71e-05 |
| DNA-Lib5 | 0.00148 | 9.16e-05 | 1.63e-05 |
| DNA-Lib6 | 0.0015 | 9.44e-05 | 1.64e-05 |
| DNA-WT | 7.06e-08 | 0 | 0 |

Again, I calculate the corrected mutation average mutation frequencies by subtracting the appropriate error control for each sample as above.

In [33]:

```
mutvirus = (pd.concat([codonmuttypes[codonmuttypes['name'].str.contains('A549-Lib')], 
                     codonmuttypes[codonmuttypes['name'].str.contains('CCL141-Lib')]
                     ])
            .assign(experiment=lambda x: x.name,
                  sample='mutvirus'))
wtvirus = (pd.concat(
    [codonmuttypes[codonmuttypes['name'].str.contains('A549-WT')]\
    .assign(experiment='A549-{0}'.format(i),
           sample='wtvirus')
    for i in ['Lib4','Lib5','Lib6']
    ] +\
    [codonmuttypes[codonmuttypes['name'].str.contains('CCL141-WT')]\
    .assign(experiment='CCL141-{0}'.format(i),
           sample='wtvirus')
    for i in ['Lib4','Lib5','Lib6']
    ]))
mutDNA = (pd.concat(
    [codonmuttypes[codonmuttypes['name'].str.contains('DNA-Lib')]\
    .assign(experiment=lambda x:x.name.str.replace('DNA', i),
            sample='mutDNA')
     for i in ['A549', 'CCL141']]))
wtDNA = (pd.concat(
    [codonmuttypes[codonmuttypes['name'].str.contains('DNA-WT')]\
    .assign(experiment='{0}-{1}'.format(cell, i),
            sample='wtDNA')
    for cell in ['A549', 'CCL141']
    for i in ['Lib4','Lib5','Lib6']]))

codonmuttypes_corr = (pd.concat([wtDNA, mutDNA, wtvirus, mutvirus])
                      .rename(columns={'nonsynonymous':'nonsyn', 'synonymous':'syn'})
                      .melt(id_vars=['experiment', 'sample'],
                            value_vars=['stop', 'nonsyn', 'syn'],
                            var_name='muttype',
                            value_name='mutfreq')
                      .pivot_table(index=['experiment', 'muttype'], columns='sample', values='mutfreq')
                      .assign(pre=lambda x: x.mutDNA - x.wtDNA,
                              post=lambda x: x.mutvirus - x.wtvirus)
                      .assign(percent=lambda x: 100 * x.post / x.pre)
                      .assign(pre2=lambda x: x.mutDNA - x.wtDNA,
                              post2=lambda x: x.mutvirus - x.wtDNA)
                      .assign(percent2=lambda x: 100 * x.post2 / x.pre2)
                     )
display(HTML(codonmuttypes_corr.to_html(float_format=lambda x: '{0:.3g}'.format(x))))
```

|  | sample | mutDNA | mutvirus | wtDNA | wtvirus | pre | post | percent | pre2 | post2 | percent2 |
| --- | --- | --- | --- | --- | --- | --- | --- | --- | --- | --- | --- |
| experiment | muttype |  |  |  |  |  |  |  |  |  |  |
| A549-Lib4 | nonsyn | 0.00153 | 0.000405 | 7.06e-08 | 2.45e-06 | 0.00153 | 0.000403 | 26.4 | 0.00153 | 0.000405 | 26.5 |
|  | stop | 9.57e-05 | 2.44e-06 | 0 | 2.59e-08 | 9.57e-05 | 2.42e-06 | 2.53 | 9.57e-05 | 2.44e-06 | 2.55 |
|  | syn | 1.71e-05 | 1.16e-05 | 0 | 2.59e-08 | 1.71e-05 | 1.16e-05 | 68 | 1.71e-05 | 1.16e-05 | 68.2 |
| A549-Lib5 | nonsyn | 0.00148 | 0.000526 | 7.06e-08 | 2.45e-06 | 0.00148 | 0.000523 | 35.4 | 0.00148 | 0.000526 | 35.5 |
|  | stop | 9.16e-05 | 6.77e-06 | 0 | 2.59e-08 | 9.16e-05 | 6.74e-06 | 7.36 | 9.16e-05 | 6.77e-06 | 7.39 |
|  | syn | 1.63e-05 | 1.43e-05 | 0 | 2.59e-08 | 1.63e-05 | 1.43e-05 | 87.6 | 1.63e-05 | 1.43e-05 | 87.7 |
| A549-Lib6 | nonsyn | 0.0015 | 0.000431 | 7.06e-08 | 2.45e-06 | 0.0015 | 0.000428 | 28.6 | 0.0015 | 0.000431 | 28.8 |
|  | stop | 9.44e-05 | 4.01e-06 | 0 | 2.59e-08 | 9.44e-05 | 3.98e-06 | 4.22 | 9.44e-05 | 4.01e-06 | 4.24 |
|  | syn | 1.64e-05 | 1.32e-05 | 0 | 2.59e-08 | 1.64e-05 | 1.32e-05 | 80.6 | 1.64e-05 | 1.32e-05 | 80.8 |
| CCL141-Lib4 | nonsyn | 0.00153 | 0.000523 | 7.06e-08 | 6.06e-07 | 0.00153 | 0.000522 | 34.2 | 0.00153 | 0.000523 | 34.2 |
|  | stop | 9.57e-05 | 3.8e-06 | 0 | 2.75e-08 | 9.57e-05 | 3.78e-06 | 3.95 | 9.57e-05 | 3.8e-06 | 3.98 |
|  | syn | 1.71e-05 | 1.23e-05 | 0 | 0 | 1.71e-05 | 1.23e-05 | 72.3 | 1.71e-05 | 1.23e-05 | 72.3 |
| CCL141-Lib5 | nonsyn | 0.00148 | 0.000517 | 7.06e-08 | 6.06e-07 | 0.00148 | 0.000517 | 34.9 | 0.00148 | 0.000517 | 35 |
|  | stop | 9.16e-05 | 6.26e-06 | 0 | 2.75e-08 | 9.16e-05 | 6.24e-06 | 6.81 | 9.16e-05 | 6.26e-06 | 6.84 |
|  | syn | 1.63e-05 | 1.35e-05 | 0 | 0 | 1.63e-05 | 1.35e-05 | 82.6 | 1.63e-05 | 1.35e-05 | 82.6 |
| CCL141-Lib6 | nonsyn | 0.0015 | 0.000528 | 7.06e-08 | 6.06e-07 | 0.0015 | 0.000528 | 35.2 | 0.0015 | 0.000528 | 35.3 |
|  | stop | 9.44e-05 | 4.95e-06 | 0 | 2.75e-08 | 9.44e-05 | 4.93e-06 | 5.22 | 9.44e-05 | 4.95e-06 | 5.25 |
|  | syn | 1.64e-05 | 1.31e-05 | 0 | 0 | 1.64e-05 | 1.31e-05 | 80 | 1.64e-05 | 1.31e-05 | 80 |

#### Copy files to paper figures directory¶

In [34]:

```
myfiguresdir = os.path.join(figuresdir, 'Fig1/')
if not os.path.isdir(myfiguresdir):
    os.mkdir(myfiguresdir)

files = [countsplotprefix + '_codonmuttypes.pdf',
         countsplotprefix + '_codonmuttypes_notaccessible.pdf',
        countsplotprefix + '_cumulmutcounts.pdf']
for f in files:
    shutil.copy(f, myfiguresdir)
```
