## Supplementary material for "Comprehensive mapping of avian influenza polymerase adaptation to the human host": File S2: Fig2_InferPreferences.html


### Infer Preferences¶

##### Import modules, define directories¶

In [2]:

```
import os
import re
import glob
import shutil
import subprocess
import string
import warnings
warnings.simplefilter('ignore') # don't print warnings to avoid clutter in notebook

import pandas as pd
import numpy as np
from scipy import stats
import matplotlib as mpl
import matplotlib.pyplot as plt
%matplotlib inline

from Bio import SeqIO

import dms_tools2
import dms_tools2.sra
import dms_tools2.plot
import dms_tools2.prefs
import dms_tools2.compareprefs
import dms_tools2.rplot
from dms_tools2.ipython_utils import showPDF
from dms_tools2.plot import plotCorrMatrix
from pymodules.utils import plotCorrMatrix2
from dms_tools2.prefs import prefsToMutEffects
from dms_tools2 import AAS

from IPython.display import display, HTML, Markdown, Image

from pymodules.utils import annotateCodonCounts_accessible 
from pymodules.utils import plotCodonMutTypes_accessible

print("Using dms_tools2 version {0}".format(dms_tools2.__version__))

# CPUs to use, should not exceed the number you request with slurm
ncpus = 14

# do we use existing results or generate everything new?
use_existing = 'yes'

# Directories
fastqdir = './fastq/'
datadir = './data'
resultsdir = './results/' 
countsdir = os.path.join(resultsdir, 'codoncounts/')
countsplotprefix = os.path.join(countsdir, 'summary')

cells = ['A549', 'CCL141']
prefsmethod = [('Bayes2err', 'prefs1'), ('Bayes2errA549', 'prefs')]
prefsdir = {cell: os.path.join(resultsdir, 'prefs_{0}'.format(cell)) for cell in cells}
prefsdir1 = {cell: os.path.join(resultsdir, 'prefs1_{0}'.format(cell)) for cell in cells}
prefsmethoddir = {'Bayes2errA549': prefsdir, #method Bayesian, with A549-WT as the post-err control, because CCL-141WT has high mutation rate
                  'Bayes2err': prefsdir1} #method Bayesian, with respective host cell WT post-err ctrls
phydmsdir = os.path.join(resultsdir, 'phydms_analysis/')
prefs_summaryplotsdir= os.path.join(resultsdir, 'prefs_summaryplots/')

paperdir = './paper'
figuresdir = os.path.join(paperdir, 'figures/')

for xdir in ([fastqdir, datadir, resultsdir, countsdir, paperdir, figuresdir]
             + list(prefsdir.values()) + list(prefsdir1.values())
             + [phydmsdir, prefs_summaryplotsdir]):
    if not os.path.isdir(xdir):
        os.mkdir(xdir)
```

```
Using dms_tools2 version 2.3.0
```

```
/fh/fast/bloom_j/computational_notebooks/yqsoh/2018/PB2-DMS/pymodules/utils.py:14: UserWarning: matplotlib.pyplot as already been imported, this call will have no effect.
  matplotlib.use('pdf')
```

In [3]:

```
## Set plotting parameters/preferences
# Set colors to use throughout
palette ={"None":"k",
          "Human":"#d95f02", "A549":"#d95f02", "Known - Human":"#d95f02", "H7N9 - Human":"#d95f02",
          "Bird":"#1b9e77", "Avian":"#1b9e77", "CCL141":"#1b9e77", 
          "Both":"#7570b3", "Known - Both":"#7570b3",
         }
```

##### Define samples¶

In [8]:

```
countsbatchfile = os.path.join(countsdir, 'batch.csv')
samples = pd.read_csv(countsbatchfile)
display(HTML(samples.to_html(index=False)))
```

| name | R1 |
| --- | --- |
| A549-Lib4 | A549-MutVirusLib-Rep1\_R1.fastq.gz |
| A549-Lib5 | A549-MutVirusLib-Rep2\_R1.fastq.gz |
| A549-Lib6 | A549-MutVirusLib-Rep3\_R1.fastq.gz |
| A549-WT | A549-WT\_R1.fastq.gz |
| CCL141-Lib4 | CCL141-MutVirusLib-Rep1\_R1.fastq.gz |
| CCL141-Lib5 | CCL141-MutVirusLib-Rep2\_R1.fastq.gz |
| CCL141-Lib6 | CCL141-MutVirusLib-Rep3\_R1.fastq.gz |
| CCL141-WT | CCL141-WT\_R1.fastq.gz |
| DNA-Lib4 | DNA-Lib-Rep1\_R1.fastq.gz |
| DNA-Lib5 | DNA-Lib-Rep2\_R1.fastq.gz |
| DNA-Lib6 | DNA-Lib-Rep3\_R1.fastq.gz |
| DNA-WT | DNA-WT\_R1.fastq.gz |

Codon counts generated in `Fig1_ProcessDMSdata.ipynb`:

In [9]:

```
!ls {countsdir}/*_codoncounts.csv
```

```
./results/codoncounts//A549-Lib4_codoncounts.csv
./results/codoncounts//A549-Lib5_codoncounts.csv
./results/codoncounts//A549-Lib6_codoncounts.csv
./results/codoncounts//A549-WT_codoncounts.csv
./results/codoncounts//CCL141-Lib4_codoncounts.csv
./results/codoncounts//CCL141-Lib5_codoncounts.csv
./results/codoncounts//CCL141-Lib6_codoncounts.csv
./results/codoncounts//CCL141-WT_codoncounts.csv
./results/codoncounts//DNA-Lib4_codoncounts.csv
./results/codoncounts//DNA-Lib5_codoncounts.csv
./results/codoncounts//DNA-Lib6_codoncounts.csv
./results/codoncounts//DNA-WT_codoncounts.csv
```

#### Amino-acid preferences¶

##### Estimate the amino-acid preferences¶

We would like to analyze the counts of each mutation pre- and post-selection to estimate the preference of each site for each amino acid.
We do this using the dms2\_batch\_prefs program.

Because of the high amount of oxidative damage especially in the CCL141-WT control and DNA-WT control, I therefore tried a couple of ways of error correction. I used `--method bayesian`, and tried using both the respective species WT for post-error controls, as well as A549-WT as the post-error control for both A549 and CCL141 passaged samples. These results turn out to be almost identical. As such, subsequent analyses will use A549-WT as the error control for both A549 and CCL141 passaged samples.

These conditions as summarized:

| Method | Error control | Cell | outdir | Result | notes |
| --- | --- | --- | --- | --- | --- |
| Bayesian | DNA-WT, A549-WT | A549 | prefs\_A549 | Success | *Should be same as prefs1\_A549* |
| Bayesian | DNA-WT, A549-WT | CCL141 | prefs\_CCL141 | Success | *Used A549-WT instead of CCL141 for post-error control* |
| Bayesian | DNA-WT, A549-WT | A549 | prefs1\_A549 | Success |  |
| Bayesian | DNA-WT, CCL141-WT | CCL141 | prefs1\_CCL141 | Success |  |

In [10]:

```
# Create batch file for dms2_batch_prefs, for each cell type, using A549-WT for both post-err controls.
prefsbatchA549 = {}
prefsbatchA549['A549'] = pd.DataFrame(
        columns=['name', 'pre', 'post', 'errpre', 'errpost'],
        data=[('A549-Lib4', 'DNA-Lib4', 'A549-Lib4', 'DNA-WT', 'A549-WT'),
              ('A549-Lib5', 'DNA-Lib5', 'A549-Lib5', 'DNA-WT', 'A549-WT'),
              ('A549-Lib6', 'DNA-Lib6', 'A549-Lib6', 'DNA-WT', 'A549-WT'),
              ])
prefsbatchA549['CCL141'] = pd.DataFrame(
        columns=['name', 'pre', 'post', 'errpre', 'errpost'],
        data=[('CCL141-Lib4', 'DNA-Lib4', 'CCL141-Lib4', 'DNA-WT', 'A549-WT'),
              ('CCL141-Lib5', 'DNA-Lib5', 'CCL141-Lib5', 'DNA-WT', 'A549-WT'),
              ('CCL141-Lib6', 'DNA-Lib6', 'CCL141-Lib6', 'DNA-WT', 'A549-WT'),
              ])
# Create batch file for dms2_batch_prefs, for each cell type, using post-selection error controls from respective cells
prefsbatch = {}
prefsbatch['A549'] = pd.DataFrame(
        columns=['name', 'pre', 'post', 'errpre', 'errpost'],
        data=[('A549-Lib4', 'DNA-Lib4', 'A549-Lib4', 'DNA-WT', 'A549-WT'),
              ('A549-Lib5', 'DNA-Lib5', 'A549-Lib5', 'DNA-WT', 'A549-WT'),
              ('A549-Lib6', 'DNA-Lib6', 'A549-Lib6', 'DNA-WT', 'A549-WT'),
              ])
prefsbatch['CCL141'] = pd.DataFrame(
        columns=['name', 'pre', 'post', 'errpre', 'errpost'],
        data=[('CCL141-Lib4', 'DNA-Lib4', 'CCL141-Lib4', 'DNA-WT', 'CCL141-WT'),
              ('CCL141-Lib5', 'DNA-Lib5', 'CCL141-Lib5', 'DNA-WT', 'CCL141-WT'),
              ('CCL141-Lib6', 'DNA-Lib6', 'CCL141-Lib6', 'DNA-WT', 'CCL141-WT'),
              ])

prefsarg = {'Bayes2errA549': ('bayesianA549', prefsbatchA549),
            'Bayes2err': ('bayesian', prefsbatch),
           }
```

In [11]:

```
for methoderr, (method, prefsbatchi) in prefsarg.items():
    for cell in cells:
        outdir = prefsmethoddir[methoderr][cell]
        print('\n*Infer preferences for Cell {1}; Method {0}; Prefsdir {2}'.format(methoderr, cell, outdir))
        prefsbatchfile = os.path.join(outdir, 'batch.csv')
        display(HTML(prefsbatchi[cell].to_html(index=False)))
        prefsbatchi[cell].to_csv(prefsbatchfile, index=False)
        cmd = 'dms2_batch_prefs --indir {0} --batchfile {1} --outdir {2} --summaryprefix summary --use_existing {3} --method {4}'\
            .format(countsdir, prefsbatchfile, outdir, use_existing, 'bayesian')
        print(cmd + '\n')
        !$cmd
```

```
*Infer preferences for Cell A549; Method Bayes2errA549; Prefsdir ./results/prefs_A549
```

| name | pre | post | errpre | errpost |
| --- | --- | --- | --- | --- |
| A549-Lib4 | DNA-Lib4 | A549-Lib4 | DNA-WT | A549-WT |
| A549-Lib5 | DNA-Lib5 | A549-Lib5 | DNA-WT | A549-WT |
| A549-Lib6 | DNA-Lib6 | A549-Lib6 | DNA-WT | A549-WT |

```
dms2_batch_prefs --indir ./results/codoncounts/ --batchfile ./results/prefs_A549/batch.csv --outdir ./results/prefs_A549 --summaryprefix summary --use_existing yes --method bayesian

Output summary files already exist and '--use_existing' is 'yes', so exiting with no further action.

*Infer preferences for Cell CCL141; Method Bayes2errA549; Prefsdir ./results/prefs_CCL141
```

| name | pre | post | errpre | errpost |
| --- | --- | --- | --- | --- |
| CCL141-Lib4 | DNA-Lib4 | CCL141-Lib4 | DNA-WT | A549-WT |
| CCL141-Lib5 | DNA-Lib5 | CCL141-Lib5 | DNA-WT | A549-WT |
| CCL141-Lib6 | DNA-Lib6 | CCL141-Lib6 | DNA-WT | A549-WT |

```
dms2_batch_prefs --indir ./results/codoncounts/ --batchfile ./results/prefs_CCL141/batch.csv --outdir ./results/prefs_CCL141 --summaryprefix summary --use_existing yes --method bayesian

Output summary files already exist and '--use_existing' is 'yes', so exiting with no further action.

*Infer preferences for Cell A549; Method Bayes2err; Prefsdir ./results/prefs1_A549
```

| name | pre | post | errpre | errpost |
| --- | --- | --- | --- | --- |
| A549-Lib4 | DNA-Lib4 | A549-Lib4 | DNA-WT | A549-WT |
| A549-Lib5 | DNA-Lib5 | A549-Lib5 | DNA-WT | A549-WT |
| A549-Lib6 | DNA-Lib6 | A549-Lib6 | DNA-WT | A549-WT |

```
dms2_batch_prefs --indir ./results/codoncounts/ --batchfile ./results/prefs1_A549/batch.csv --outdir ./results/prefs1_A549 --summaryprefix summary --use_existing yes --method bayesian

Output summary files already exist and '--use_existing' is 'yes', so exiting with no further action.

*Infer preferences for Cell CCL141; Method Bayes2err; Prefsdir ./results/prefs1_CCL141
```

| name | pre | post | errpre | errpost |
| --- | --- | --- | --- | --- |
| CCL141-Lib4 | DNA-Lib4 | CCL141-Lib4 | DNA-WT | CCL141-WT |
| CCL141-Lib5 | DNA-Lib5 | CCL141-Lib5 | DNA-WT | CCL141-WT |
| CCL141-Lib6 | DNA-Lib6 | CCL141-Lib6 | DNA-WT | CCL141-WT |

```
dms2_batch_prefs --indir ./results/codoncounts/ --batchfile ./results/prefs1_CCL141/batch.csv --outdir ./results/prefs1_CCL141 --summaryprefix summary --use_existing yes --method bayesian

Output summary files already exist and '--use_existing' is 'yes', so exiting with no further action.
```

This creates files containing the estimated amino-acid preferences for each replicate. These preferences are in the directory specified by --outdir in the call to dms2\_batch\_prefs above:

In [12]:

```
for methoderr, i in prefsmethod:
    print(methoderr+":")
    for cell in cells:
        !ls {prefsmethoddir[methoderr][cell]}/{cell}*_prefs.csv
```

```
Bayes2err:
./results/prefs1_A549/A549-Lib4_prefs.csv
./results/prefs1_A549/A549-Lib5_prefs.csv
./results/prefs1_A549/A549-Lib6_prefs.csv
./results/prefs1_CCL141/CCL141-Lib4_prefs.csv
./results/prefs1_CCL141/CCL141-Lib5_prefs.csv
./results/prefs1_CCL141/CCL141-Lib6_prefs.csv
Bayes2errA549:
./results/prefs_A549/A549-Lib4_prefs.csv
./results/prefs_A549/A549-Lib5_prefs.csv
./results/prefs_A549/A549-Lib6_prefs.csv
./results/prefs_CCL141/CCL141-Lib4_prefs.csv
./results/prefs_CCL141/CCL141-Lib5_prefs.csv
./results/prefs_CCL141/CCL141-Lib6_prefs.csv
```

We can look at the summary files created by dms2\_batch\_prefs.
These files (for each cell type) are in the directory specified by `--outdir` and have the prefix specified by `--summaryprefix`.

To look at all correlations at once, I generated summary correlation plots comparing across cell types. These are in the directory `results/prefs_summaryplots`.

In [13]:

```
def corrplottypes(datatype):
    print('Datatype: ' + datatype)
#     sns.set_style("ticks")
    names = ['{0}-Lib{1}'.format(cell, i) for cell in cells for i in ['4', '5', '6']]
    print('Samples:\n' + '\n'.join(names))
    for methoderr, i in prefsmethod:
        print ('* Prefs method:' + methoderr)
        if datatype in ['prefs', 'logprefs']:
            infiles = ['{0}/{1}-Lib{2}_prefs.csv'.format(prefsmethoddir[methoderr][cell], cell, i) for cell in cells for i in ['4', '5', '6']]
            print('Files:\n' + '\n'.join(infiles))
        elif datatype in ['effect', 'log2effect']:
            infiles = ['{0}/{1}-Lib{2}_muteffectwt.csv'.format(prefsmethoddir[methoderr][cell], cell, i) for cell in cells for i in ['4', '5', '6']]
            print('Files:\n' + '\n'.join(infiles))
        plotfile = os.path.join(prefs_summaryplotsdir, '{0}{1}_{2}corr.pdf'.format(i, methoderr, datatype))
        print('Plotfile: ' + plotfile)
        colors=[palette['A549']]*3 + [palette['None']]*6 + [palette['CCL141']] + [palette['None']]*3 + [palette['CCL141']]*2
        if (datatype == 'prefs'):
            plotCorrMatrix(names, infiles, plotfile, datatype, colors=colors)
        elif datatype in ['logprefs', 'effect', 'log2effect']:
            plotCorrMatrix2(names, infiles, plotfile, datatype, colors=colors)
            
def plotcorrplots(datatype):
    fs = {}
    for methoderr, i in prefsmethod:
        fs[methoderr] = os.path.join(prefs_summaryplotsdir, '{0}{1}_{2}corr.pdf'.format(i, methoderr, datatype))
    print('{0} by method: {1} ; {2}'.format(datatype, 'Bayesian with 2err controls', 'Bayesian with 2 err controls, but using A549-WT for all post-err'))
    print('Files: {0} ; {1}'.format(fs['Bayes2err'], fs['Bayes2errA549']))
    showPDF([fs['Bayes2err'], fs['Bayes2errA549']], width=1000)
```

In [14]:

```
corrplottypes('prefs')
```

```
Datatype: prefs
Samples:
A549-Lib4
A549-Lib5
A549-Lib6
CCL141-Lib4
CCL141-Lib5
CCL141-Lib6
* Prefs method:Bayes2err
Files:
./results/prefs1_A549/A549-Lib4_prefs.csv
./results/prefs1_A549/A549-Lib5_prefs.csv
./results/prefs1_A549/A549-Lib6_prefs.csv
./results/prefs1_CCL141/CCL141-Lib4_prefs.csv
./results/prefs1_CCL141/CCL141-Lib5_prefs.csv
./results/prefs1_CCL141/CCL141-Lib6_prefs.csv
Plotfile: ./results/prefs_summaryplots/prefs1Bayes2err_prefscorr.pdf
* Prefs method:Bayes2errA549
Files:
./results/prefs_A549/A549-Lib4_prefs.csv
./results/prefs_A549/A549-Lib5_prefs.csv
./results/prefs_A549/A549-Lib6_prefs.csv
./results/prefs_CCL141/CCL141-Lib4_prefs.csv
./results/prefs_CCL141/CCL141-Lib5_prefs.csv
./results/prefs_CCL141/CCL141-Lib6_prefs.csv
Plotfile: ./results/prefs_summaryplots/prefsBayes2errA549_prefscorr.pdf
```

Both ways of inferring preferences were very similar (either using the respective post-err control, or using the A549-WT as the post-err control for both cell types). In general, we observe good correlations between biological replicates within both cell lines. Correlations are a little worse when comparing between cell lines, as expected.

In [15]:

```
plotcorrplots('prefs')
```

```
prefs by method: Bayesian with 2err controls ; Bayesian with 2 err controls, but using A549-WT for all post-err
Files: ./results/prefs_summaryplots/prefs1Bayes2err_prefscorr.pdf ; ./results/prefs_summaryplots/prefsBayes2errA549_prefscorr.pdf
```

##### Mutational effects¶

We can also represent the effects of mutations by their preference relative to the wild-type amino acid. If we take the log of this mutational effect, negative values indicate deleterious mutations and positive values indicate favorable mutations relative to the wild-type amino acid.

Here we calculate the mutational effects and then plot their log2 values on a logo plot.

In [16]:

```
# Get wild-type amino acids
wtoverlayfile = os.path.join(datadir, 'wildtypeoverlayfile.csv')
aacounts = dms_tools2.utils.codonToAACounts(
        pd.read_csv(os.path.join(countsdir, 'DNA-WT_codoncounts.csv')))
aacounts.query('wildtype != "*"')[['site', 'wildtype']].to_csv(wtoverlayfile, index=False)

# Calculate and save mutational effects
wt = pd.read_csv(wtoverlayfile)
for methoderr, i in prefsmethod:
    for cell in cells:
        prefsfileprefix = ['{0}-Lib{1}_'.format(cell, lib) for lib in ['4','5','6']]\
                + ['summary_avg']
        for pre in prefsfileprefix:
            prefsfile = os.path.join(prefsmethoddir[methoderr][cell], '{0}prefs.csv'.format(pre))
            outfile = os.path.join(prefsmethoddir[methoderr][cell], '{0}muteffect.csv'.format(pre))
            outfilewt = os.path.join(prefsmethoddir[methoderr][cell], '{0}muteffectwt.csv'.format(pre))
            print(prefsfile+' --> '+outfile+', '+outfilewt)
            prefs = pd.read_csv(prefsfile)
            if not os.path.exists(outfile):
                muteffect = prefsToMutEffects(prefs, dms_tools2.AAS)
                muteffectwt = pd.merge(wt, muteffect, how='left', left_on=['site', 'wildtype'], right_on=['site', 'initial'], sort=False)
                muteffect.to_csv(outfile, index=False)
                muteffectwt.to_csv(outfilewt, index=False)
            else:
                print(outfile+' exists!')
```

```
./results/prefs1_A549/A549-Lib4_prefs.csv --> ./results/prefs1_A549/A549-Lib4_muteffect.csv, ./results/prefs1_A549/A549-Lib4_muteffectwt.csv
./results/prefs1_A549/A549-Lib4_muteffect.csv exists!
./results/prefs1_A549/A549-Lib5_prefs.csv --> ./results/prefs1_A549/A549-Lib5_muteffect.csv, ./results/prefs1_A549/A549-Lib5_muteffectwt.csv
./results/prefs1_A549/A549-Lib5_muteffect.csv exists!
./results/prefs1_A549/A549-Lib6_prefs.csv --> ./results/prefs1_A549/A549-Lib6_muteffect.csv, ./results/prefs1_A549/A549-Lib6_muteffectwt.csv
./results/prefs1_A549/A549-Lib6_muteffect.csv exists!
./results/prefs1_A549/summary_avgprefs.csv --> ./results/prefs1_A549/summary_avgmuteffect.csv, ./results/prefs1_A549/summary_avgmuteffectwt.csv
./results/prefs1_A549/summary_avgmuteffect.csv exists!
./results/prefs1_CCL141/CCL141-Lib4_prefs.csv --> ./results/prefs1_CCL141/CCL141-Lib4_muteffect.csv, ./results/prefs1_CCL141/CCL141-Lib4_muteffectwt.csv
./results/prefs1_CCL141/CCL141-Lib4_muteffect.csv exists!
./results/prefs1_CCL141/CCL141-Lib5_prefs.csv --> ./results/prefs1_CCL141/CCL141-Lib5_muteffect.csv, ./results/prefs1_CCL141/CCL141-Lib5_muteffectwt.csv
./results/prefs1_CCL141/CCL141-Lib5_muteffect.csv exists!
./results/prefs1_CCL141/CCL141-Lib6_prefs.csv --> ./results/prefs1_CCL141/CCL141-Lib6_muteffect.csv, ./results/prefs1_CCL141/CCL141-Lib6_muteffectwt.csv
./results/prefs1_CCL141/CCL141-Lib6_muteffect.csv exists!
./results/prefs1_CCL141/summary_avgprefs.csv --> ./results/prefs1_CCL141/summary_avgmuteffect.csv, ./results/prefs1_CCL141/summary_avgmuteffectwt.csv
./results/prefs1_CCL141/summary_avgmuteffect.csv exists!
./results/prefs_A549/A549-Lib4_prefs.csv --> ./results/prefs_A549/A549-Lib4_muteffect.csv, ./results/prefs_A549/A549-Lib4_muteffectwt.csv
./results/prefs_A549/A549-Lib4_muteffect.csv exists!
./results/prefs_A549/A549-Lib5_prefs.csv --> ./results/prefs_A549/A549-Lib5_muteffect.csv, ./results/prefs_A549/A549-Lib5_muteffectwt.csv
./results/prefs_A549/A549-Lib5_muteffect.csv exists!
./results/prefs_A549/A549-Lib6_prefs.csv --> ./results/prefs_A549/A549-Lib6_muteffect.csv, ./results/prefs_A549/A549-Lib6_muteffectwt.csv
./results/prefs_A549/A549-Lib6_muteffect.csv exists!
./results/prefs_A549/summary_avgprefs.csv --> ./results/prefs_A549/summary_avgmuteffect.csv, ./results/prefs_A549/summary_avgmuteffectwt.csv
./results/prefs_A549/summary_avgmuteffect.csv exists!
./results/prefs_CCL141/CCL141-Lib4_prefs.csv --> ./results/prefs_CCL141/CCL141-Lib4_muteffect.csv, ./results/prefs_CCL141/CCL141-Lib4_muteffectwt.csv
./results/prefs_CCL141/CCL141-Lib4_muteffect.csv exists!
./results/prefs_CCL141/CCL141-Lib5_prefs.csv --> ./results/prefs_CCL141/CCL141-Lib5_muteffect.csv, ./results/prefs_CCL141/CCL141-Lib5_muteffectwt.csv
./results/prefs_CCL141/CCL141-Lib5_muteffect.csv exists!
./results/prefs_CCL141/CCL141-Lib6_prefs.csv --> ./results/prefs_CCL141/CCL141-Lib6_muteffect.csv, ./results/prefs_CCL141/CCL141-Lib6_muteffectwt.csv
./results/prefs_CCL141/CCL141-Lib6_muteffect.csv exists!
./results/prefs_CCL141/summary_avgprefs.csv --> ./results/prefs_CCL141/summary_avgmuteffect.csv, ./results/prefs_CCL141/summary_avgmuteffectwt.csv
./results/prefs_CCL141/summary_avgmuteffect.csv exists!
```

In [17]:

```
corrplottypes('log2effect')
```

```
Datatype: log2effect
Samples:
A549-Lib4
A549-Lib5
A549-Lib6
CCL141-Lib4
CCL141-Lib5
CCL141-Lib6
* Prefs method:Bayes2err
Files:
./results/prefs1_A549/A549-Lib4_muteffectwt.csv
./results/prefs1_A549/A549-Lib5_muteffectwt.csv
./results/prefs1_A549/A549-Lib6_muteffectwt.csv
./results/prefs1_CCL141/CCL141-Lib4_muteffectwt.csv
./results/prefs1_CCL141/CCL141-Lib5_muteffectwt.csv
./results/prefs1_CCL141/CCL141-Lib6_muteffectwt.csv
Plotfile: ./results/prefs_summaryplots/prefs1Bayes2err_log2effectcorr.pdf
* Prefs method:Bayes2errA549
Files:
./results/prefs_A549/A549-Lib4_muteffectwt.csv
./results/prefs_A549/A549-Lib5_muteffectwt.csv
./results/prefs_A549/A549-Lib6_muteffectwt.csv
./results/prefs_CCL141/CCL141-Lib4_muteffectwt.csv
./results/prefs_CCL141/CCL141-Lib5_muteffectwt.csv
./results/prefs_CCL141/CCL141-Lib6_muteffectwt.csv
Plotfile: ./results/prefs_summaryplots/prefsBayes2errA549_log2effectcorr.pdf
```

In [18]:

```
plotcorrplots('log2effect')
```

```
log2effect by method: Bayesian with 2err controls ; Bayesian with 2 err controls, but using A549-WT for all post-err
Files: ./results/prefs_summaryplots/prefs1Bayes2err_log2effectcorr.pdf ; ./results/prefs_summaryplots/prefsBayes2errA549_log2effectcorr.pdf
```

##### Visualize preferences¶

Our run of dms2\_batch\_prefs also creates a file with the **average** preferences across replicates, `summary_avgprefs.csv`.
Let's use dms2\_logoplot to visualize these average preferences in the form of a logo plot, where the height of each letter is proportional to the preference for that amino acid.

In [19]:

```
# We already have wtoverlayfile
wtoverlayfile = os.path.join(datadir, 'wildtypeoverlayfile.csv')
# Get domains annotation of PB2
domainsFile = os.path.join(datadir, 'domains.csv')
```

In [20]:

```
for methoderr, i in prefsmethod:
    for cell in cells:
        outdir = prefsmethoddir[methoderr][cell]
        avgprefs = os.path.join(outdir, 'summary_avgprefs.csv')
        logoname = 'logoAvgprefs'
        print('** ' + methoderr + ', ' + cell+':')
        print('avgprefs: {0}'.format(avgprefs))
        print('logoname: {0}'.format(logoname))
        print('outdir: {0}'.format(outdir))
        print('overlay1: {0}'.format(wtoverlayfile))
        print('overlay2: {0}'.format(domainsFile))

        log = !dms2_logoplot \
                --prefs {avgprefs} \
                --name {logoname} \
                --outdir {outdir} \
                --overlay1 {wtoverlayfile} wildtype wildtype \
                --overlay2 {domainsFile} D "Domains"\
                --nperline 100 \
                --letterheight 1 \
                --numberevery 10 \
                --use_existing {use_existing}
#         print("\n".join(log))

        logoplot = os.path.join(outdir, '{0}_prefs.pdf'.format(logoname))
        print('{0}, {1}: {2}'.format(methoderr, cell, logoplot))
        showPDF(logoplot, width=1000)
```

```
** Bayes2err, A549:
avgprefs: ./results/prefs1_A549/summary_avgprefs.csv
logoname: logoAvgprefs
outdir: ./results/prefs1_A549
overlay1: ./data/wildtypeoverlayfile.csv
overlay2: ./data/domains.csv
Bayes2err, A549: ./results/prefs1_A549/logoAvgprefs_prefs.pdf
```

```
** Bayes2err, CCL141:
avgprefs: ./results/prefs1_CCL141/summary_avgprefs.csv
logoname: logoAvgprefs
outdir: ./results/prefs1_CCL141
overlay1: ./data/wildtypeoverlayfile.csv
overlay2: ./data/domains.csv
Bayes2err, CCL141: ./results/prefs1_CCL141/logoAvgprefs_prefs.pdf
```

```
** Bayes2errA549, A549:
avgprefs: ./results/prefs_A549/summary_avgprefs.csv
logoname: logoAvgprefs
outdir: ./results/prefs_A549
overlay1: ./data/wildtypeoverlayfile.csv
overlay2: ./data/domains.csv
Bayes2errA549, A549: ./results/prefs_A549/logoAvgprefs_prefs.pdf
```

```
** Bayes2errA549, CCL141:
avgprefs: ./results/prefs_CCL141/summary_avgprefs.csv
logoname: logoAvgprefs
outdir: ./results/prefs_CCL141
overlay1: ./data/wildtypeoverlayfile.csv
overlay2: ./data/domains.csv
Bayes2errA549, CCL141: ./results/prefs_CCL141/logoAvgprefs_prefs.pdf
```

#### Phylogenetic analyses with experimentally informed codon models¶

Here, we will take the averaged amino-acid preferences and fit them to natural sequence evolution.
The goal here is to test the experimentally measured preferences are informative about natural sequence evolution, and if so obtain a *stringency parameter* that can be used to re-scale them to bring them into line with the stringency of natural selection.

##### Curate PB2 sequences¶

I had downloaded complete coding sequences from the Influenza Virus Resource Database (NCBI) for:

- Human seasonal influenza PB2 (on 20180330)
- Avian influenza PB2 (on 20180320)
- All human influenza PB2, including pdmH1N1 and recent zoonotic transfers (on 20180320)

I used the latter two sets of sequences to generate a combined influenza PB2 tree.

To obtain a subsampling of sequences for phylogenetic analyses, I first aligned, removed sequences with ambiguous sequence, gaps, and trimmed the sequences to my reference sequence using phydms\_prepalignment. For each set of sequences, I subsampled 1 sequence per year. This is done by the script `subsample_IVR_sequences.py`.

In [21]:

```
# Specify downloaded sequences, add reference sequence to each set of downloaded sequences
IVRseqs = ['Human-PB2', 'Avian-PB2', 'HumanSeasonal-PB2']
S009PB2seq = 'S009-PB2'
S009PB2header = 'S009-PB2'
# Add S009 PB2 sequence to each set of IVR sequences.
for species in IVRseqs:
    infile1 = os.path.join(datadir, S009PB2seq+'.fasta')
    infile2 = os.path.join(datadir, species+'.fasta')
    filewithref = '{0}/{1}_{2}.fasta'.format(datadir, species, S009PB2header)
    if os.path.isfile(filewithref):
        print('{0} already added to {1}. File: {2}'.format(infile1, infile2, filewithref))
    else:
        print('Adding {0} to {1}. File: {2}'.format(infile1, infile2, filewithref))
        log = !cat {infile1} {infile2} > {filewithref}

# Prep samples with phydms_prepalignment
for species in IVRseqs:
    filewithref = '{0}/{1}_{2}.fasta'.format(datadir, species, S009PB2header)
    print('Prepping {0}...'.format(filewithref))
    alignmentoutfile = os.path.join(datadir, species+'_alignment.fasta')
    if os.path.isfile(alignmentoutfile):
        print('{0} already aligned and prepped by phydms_prepalignment.'.format(alignmentoutfile))
    else:
        cmd = 'phydms_prepalignment {0} {1} {2} --mafft mafft --minuniqueness 2'\
        .format(filewithref, alignmentoutfile, S009PB2header)
        print(cmd)
        !$cmd
```

```
./data/S009-PB2.fasta already added to ./data/Human-PB2.fasta. File: ./data/Human-PB2_S009-PB2.fasta
./data/S009-PB2.fasta already added to ./data/Avian-PB2.fasta. File: ./data/Avian-PB2_S009-PB2.fasta
./data/S009-PB2.fasta already added to ./data/HumanSeasonal-PB2.fasta. File: ./data/HumanSeasonal-PB2_S009-PB2.fasta
Prepping ./data/Human-PB2_S009-PB2.fasta...
./data/Human-PB2_alignment.fasta already aligned and prepped by phydms_prepalignment.
Prepping ./data/Avian-PB2_S009-PB2.fasta...
./data/Avian-PB2_alignment.fasta already aligned and prepped by phydms_prepalignment.
Prepping ./data/HumanSeasonal-PB2_S009-PB2.fasta...
./data/HumanSeasonal-PB2_alignment.fasta already aligned and prepped by phydms_prepalignment.
```

In [22]:

```
# Subsample sequences
numberSubsamples = 1
for species in IVRseqs:
    alignmentfile = os.path.join(datadir, species+'_alignment.fasta')
    subsample_prefix = '{0}/{1}_subsample'.format(datadir, species)
    logfile = os.path.join(datadir, species+'_subsample.log')
    log = !python scripts/subsample_IVR_sequences.py {alignmentfile} {numberSubsamples} {subsample_prefix}
    with open(logfile, 'w') as f:
        f.write('\n'.join(log))
        print('Subsample complete.')

# Remove outlier sequence from ./data/HumanSeasonal-PB2_subsample_0.fa
shutil.move('./data/HumanSeasonal-PB2_subsample_0.fa', './data/HumanSeasonal-PB2_subsample_0.fa.original')
outlierid = 'cds_AGQ47881_A/New_Jersey/Swiss/1976_1976//_PB2'
records = (r for r in SeqIO.parse('./data/HumanSeasonal-PB2_subsample_0.fa.original', 'fasta') if r.id != outlierid)
SeqIO.write(records, './data/HumanSeasonal-PB2_subsample_0.fa', 'fasta')

# Made a combined set of sequences for both species by combining each Human and Avian subsample.
!cat data/Human-PB2_subsample_0.fa data/Avian-PB2_subsample_0.fa >data/HumanAvian-PB2_subsample_0.fa

# Final set of sequences to use in phydms
IVRseqs2 = ['HumanAvian-PB2', 'Avian-PB2', 'HumanSeasonal-PB2']
```

```
Subsample complete.
Subsample complete.
Subsample complete.
```

Note: I manually removed an outlier from the HumanSeasonal-PB2\_subsample\_0 set of sequences from the 1976 Swine flu. It is likely a descendant of the 1918 virus that had been evolving in pigs rather than humans since 1918 before causing a brief outbreak in 1976:
http://www.sciencemag.org/news/2009/04/retrospective-what-happened-swine-flu-1976.

##### Fit preferences to the evolution of natural PB2 sequences¶

I used the preferences estimated above and fit them to the natural evolution of human, avian, and combined influenza PB2 sequences. I do this using phydms, which will fit and compare an experimentally-informed codon substitution model using the preferences, to conventional codon substitution models.

In [23]:

```
# First, I had to rename prefs files so that the file name tells us which cell type the preferences came from.
pwd = !pwd #To make symbolic links, I need the full absolute path directory!
phydmsdirfull = os.path.join(pwd[0], phydmsdir)
prefsrenamedfiles = []
for methoderr, i in prefsmethod:
    for cell in cells:
        indir = os.path.join(pwd[0], prefsmethoddir[methoderr][cell])
        avgprefs = os.path.join(indir, 'summary_avgprefs.csv')
        renamedfile = os.path.join(phydmsdirfull, '{0}avgprefs_{1}.csv'.format(cell, methoderr))
        prefsrenamedfiles.append(renamedfile)
        if not os.path.isfile(renamedfile):
            print('Linking '+avgprefs)
            print('     to '+renamedfile)
            os.symlink(avgprefs, renamedfile)
        else:
            print('Already linked to {0}'.format(renamedfile))
```

```
Already linked to /fh/fast/bloom_j/computational_notebooks/yqsoh/2018/PB2-DMS/./results/phydms_analysis/A549avgprefs_Bayes2err.csv
Already linked to /fh/fast/bloom_j/computational_notebooks/yqsoh/2018/PB2-DMS/./results/phydms_analysis/CCL141avgprefs_Bayes2err.csv
Already linked to /fh/fast/bloom_j/computational_notebooks/yqsoh/2018/PB2-DMS/./results/phydms_analysis/A549avgprefs_Bayes2errA549.csv
Already linked to /fh/fast/bloom_j/computational_notebooks/yqsoh/2018/PB2-DMS/./results/phydms_analysis/CCL141avgprefs_Bayes2errA549.csv
```

In [24]:

```
# Run phydms
prefsrenamedfiles = [os.path.join(phydmsdir, '{0}avgprefs_{1}.csv'.format(cell, methoderr)) 
                     for cell in cells for methoderr,i in prefsmethod]
phydms_input_prefs = ' '.join(prefsrenamedfiles)
raxmlpath = 'raxmlHPC'

for species in IVRseqs2:
    for i in range(0, numberSubsamples):
        alignment = './data/{0}_subsample_{1}.fa'.format(species, i)
        sample_phydms_dir = os.path.join(phydmsdir, 'default_{0}_subsample_{1}'.format(species, i))+'/'
        if not os.path.isdir(sample_phydms_dir):
            os.mkdir(sample_phydms_dir)
        
        cmd_phydms = 'phydms_comprehensive {0} {1} {2} --raxml {3}'\
            .format(sample_phydms_dir, alignment, phydms_input_prefs, raxmlpath)
        
        modelcomparison = os.path.join(sample_phydms_dir, 'modelcomparison.md')
        if use_existing == 'yes' and os.path.isfile(modelcomparison):
            print('Results of phydms analysis already exist.')
        else:
            print(cmd_phydms)
            !$cmd_phydms
```

```
Results of phydms analysis already exist.
Results of phydms analysis already exist.
Results of phydms analysis already exist.
```

The table of results compares four different models:

1. The *ExpCM* model informed by the deep mutational scanning (in either A549 or CCL141 cells).
2. A standard Goldman-Yang style model that allows the dN/dS ratio to vary across sites (the M5 model of YNGKP).
3. A standard Goldman-Yang style model with a single dN/dS ratio (the M0 model of YNGKP).
4. A "control" *ExpCM* in which the amino-acid preferences are averaged across sites to eliminate all site-specific information. Note that this is different than averaging the replicates to the get the "average preferences" -- here, the preferences are being averaged across **sites**, not replicates, so there is no longer any site-specific information.

If the deep mutational scanning is informative about evolution, then we expect it to fit the natural sequence evolution much better than any of the other models.
That is in fact what we see, as it has a much lower AIC. There is no difference between using the respective host cell WT as post-error control (`Bayes2err`), or using A549-WT for both the post-error controls (`Bayes2errA549`). For both human and avian influenza PB2 trees, preferences from A549 perform slightly better, but it's not clear if that difference is meaningful.

Also of relevance is the *stringency parameter* ($\beta$) that re-scales the preferences to optimally fit natural evolution in an *ExpCM*.
When $\beta > 1$, it means that natural evolution prefers the same amino acids as the experiments but with greater stringency (see Hilton and Bloom (2017).

In [25]:

```
for species in IVRseqs2:
    for i in range(0, numberSubsamples):
        sample_phydms_dir = os.path.join(resultsdir, 'phydms_analysis/default_{0}_subsample_{1}'.format(species, i))+'/'
        modelcomparison = os.path.join(sample_phydms_dir, 'modelcomparison.md')
        print('Model comparison for {0}: {1}'.format(species, modelcomparison))
        display(Markdown(modelcomparison))
```

```
Model comparison for HumanAvian-PB2: ./results/phydms_analysis/default_HumanAvian-PB2_subsample_0/modelcomparison.md
```

| Model | deltaAIC | LogLikelihood | nParams | ParamValues |
| --- | --- | --- | --- | --- |
| ExpCM\_A549avgprefs\_Bayes2err | 0.00 | -30634.89 | 6 | beta=2.51, kappa=8.63, omega=0.15 |
| ExpCM\_A549avgprefs\_Bayes2errA549 | 0.00 | -30634.89 | 6 | beta=2.51, kappa=8.63, omega=0.15 |
| ExpCM\_CCL141avgprefs\_Bayes2err | 258.82 | -30764.30 | 6 | beta=2.63, kappa=8.65, omega=0.14 |
| ExpCM\_CCL141avgprefs\_Bayes2errA549 | 385.04 | -30827.41 | 6 | beta=2.57, kappa=8.58, omega=0.12 |
| YNGKP\_M5 | 3507.84 | -32382.81 | 12 | alpha\_omega=0.30, beta\_omega=6.07, kappa=7.77 |
| averaged\_ExpCM\_CCL141avgprefs\_Bayes2err | 3514.96 | -32392.37 | 6 | beta=0.52, kappa=8.67, omega=0.04 |
| averaged\_ExpCM\_CCL141avgprefs\_Bayes2errA549 | 3516.34 | -32393.06 | 6 | beta=0.44, kappa=8.63, omega=0.04 |
| averaged\_ExpCM\_A549avgprefs\_Bayes2err | 3516.92 | -32393.35 | 6 | beta=0.33, kappa=8.64, omega=0.04 |
| averaged\_ExpCM\_A549avgprefs\_Bayes2errA549 | 3516.92 | -32393.35 | 6 | beta=0.33, kappa=8.64, omega=0.04 |
| YNGKP\_M0 | 4131.20 | -32695.49 | 11 | kappa=7.74, omega=0.04 |

```
Model comparison for Avian-PB2: ./results/phydms_analysis/default_Avian-PB2_subsample_0/modelcomparison.md
```

| Model | deltaAIC | LogLikelihood | nParams | ParamValues |
| --- | --- | --- | --- | --- |
| ExpCM\_A549avgprefs\_Bayes2err | 0.00 | -19943.48 | 6 | beta=2.70, kappa=9.42, omega=0.11 |
| ExpCM\_A549avgprefs\_Bayes2errA549 | 0.00 | -19943.48 | 6 | beta=2.70, kappa=9.42, omega=0.11 |
| ExpCM\_CCL141avgprefs\_Bayes2err | 156.04 | -20021.50 | 6 | beta=2.95, kappa=9.46, omega=0.10 |
| ExpCM\_CCL141avgprefs\_Bayes2errA549 | 277.00 | -20081.98 | 6 | beta=2.86, kappa=9.39, omega=0.09 |
| averaged\_ExpCM\_CCL141avgprefs\_Bayes2err | 3168.14 | -21527.55 | 6 | beta=0.36, kappa=9.46, omega=0.03 |
| averaged\_ExpCM\_CCL141avgprefs\_Bayes2errA549 | 3169.00 | -21527.98 | 6 | beta=0.31, kappa=9.46, omega=0.03 |
| averaged\_ExpCM\_A549avgprefs\_Bayes2err | 3169.22 | -21528.09 | 6 | beta=0.31, kappa=9.46, omega=0.03 |
| averaged\_ExpCM\_A549avgprefs\_Bayes2errA549 | 3169.22 | -21528.09 | 6 | beta=0.31, kappa=9.46, omega=0.03 |
| YNGKP\_M5 | 3306.58 | -21590.77 | 12 | alpha\_omega=0.30, beta\_omega=9.96, kappa=8.48 |
| YNGKP\_M0 | 3537.90 | -21707.43 | 11 | kappa=8.45, omega=0.03 |

```
Model comparison for HumanSeasonal-PB2: ./results/phydms_analysis/default_HumanSeasonal-PB2_subsample_0/modelcomparison.md
```

| Model | deltaAIC | LogLikelihood | nParams | ParamValues |
| --- | --- | --- | --- | --- |
| ExpCM\_A549avgprefs\_Bayes2err | 0.00 | -11194.41 | 6 | beta=2.80, kappa=7.24, omega=0.29 |
| ExpCM\_A549avgprefs\_Bayes2errA549 | 0.00 | -11194.41 | 6 | beta=2.80, kappa=7.24, omega=0.29 |
| ExpCM\_CCL141avgprefs\_Bayes2err | 169.32 | -11279.07 | 6 | beta=3.05, kappa=7.20, omega=0.26 |
| ExpCM\_CCL141avgprefs\_Bayes2errA549 | 285.14 | -11336.98 | 6 | beta=2.98, kappa=7.11, omega=0.23 |
| YNGKP\_M5 | 3135.52 | -12756.17 | 12 | alpha\_omega=0.30, beta\_omega=3.60, kappa=6.58 |
| averaged\_ExpCM\_CCL141avgprefs\_Bayes2err | 3187.50 | -12788.16 | 6 | beta=0.14, kappa=7.20, omega=0.08 |
| averaged\_ExpCM\_CCL141avgprefs\_Bayes2errA549 | 3187.78 | -12788.30 | 6 | beta=0.09, kappa=7.20, omega=0.08 |
| averaged\_ExpCM\_A549avgprefs\_Bayes2err | 3187.88 | -12788.35 | 6 | beta=0.06, kappa=7.20, omega=0.08 |
| averaged\_ExpCM\_A549avgprefs\_Bayes2errA549 | 3187.88 | -12788.35 | 6 | beta=0.06, kappa=7.20, omega=0.08 |
| YNGKP\_M0 | 3378.92 | -12878.87 | 11 | kappa=6.52, omega=0.07 |

Get beta values for each set of preferences, from the corresponding species tree.

In [26]:

```
beta = {}
cell_species = [('A549', 'HumanAvian-PB2'),
                ('CCL141', 'HumanAvian-PB2')]
for methoderr, i in [('Bayes2errA549', 'prefs')]:
    for cell, species in cell_species:
        print('Method {0}, preferences in {1}, fit against {2}:'.format(methoderr, cell, species))
        b = 'None'
        sample_phydms_dir = os.path.join(resultsdir, 'phydms_analysis/default_{0}_subsample_0'.format(species))
        paramsfile = os.path.join(sample_phydms_dir, 'ExpCM_{0}avgprefs_{1}_modelparams.txt'.format(cell, methoderr))
        print('  Looking in {0}:'.format(paramsfile))
        with open(paramsfile) as infile:
            for line in infile:
                if re.search('beta = ', line):
                    w = line.rstrip().split(' ')
                    b = float(w[-1])
        print('  beta average = '+str(b))
        beta[(methoderr, cell, species)] = b
```

```
Method Bayes2errA549, preferences in A549, fit against HumanAvian-PB2:
  Looking in ./results/phydms_analysis/default_HumanAvian-PB2_subsample_0/ExpCM_A549avgprefs_Bayes2errA549_modelparams.txt:
  beta average = 2.50974
Method Bayes2errA549, preferences in CCL141, fit against HumanAvian-PB2:
  Looking in ./results/phydms_analysis/default_HumanAvian-PB2_subsample_0/ExpCM_CCL141avgprefs_Bayes2errA549_modelparams.txt:
  beta average = 2.57381
```

##### Rescale preferences¶

Rescale prefs for both A549, CCL141 cells inferred by Bayes2errA549 method, using beta calculated from fitting to combined Human and Avian sequences.

In [27]:

```
species = 'HumanAvian-PB2'
for methoderr, i in [('Bayes2errA549', 'prefs')]:
    for cell in cells:
        pdir = prefsmethoddir[methoderr][cell]
        prefsfiles = [os.path.join(pdir, '{0}-Lib{1}_prefs.csv'.format(cell, i)) for i in ['4', '5', '6']] + \
                        [os.path.join(pdir, 'summary_avgprefs.csv'.format(cell, i))]
        rescaled_prefsfiles = [os.path.join(pdir, '{0}-Lib{1}_prefs_rescaled.csv'.format(cell, i)) for i in ['4', '5', '6']] + \
                        [os.path.join(pdir, 'summary_avgprefs_rescaled.csv'.format(cell, i))]
        for prefsfile, rescaled_prefsfile in zip(prefsfiles, rescaled_prefsfiles):
            print('Rescaling preference {0}, using beta of {1}'.format(prefsfile, beta[(methoderr, cell, species)]))
            print('             outfile {0}'.format(rescaled_prefsfile))
            if not (os.path.isfile(rescaled_prefsfile)):
                print('Rescaling!')
                (dms_tools2.prefs.rescalePrefs(pd.read_csv(prefsfile), beta[(methoderr, cell, species)])
                    .to_csv(rescaled_prefsfile, index=False))
            else:
                print('Already rescaled!')
```

```
Rescaling preference ./results/prefs_A549/A549-Lib4_prefs.csv, using beta of 2.50974
             outfile ./results/prefs_A549/A549-Lib4_prefs_rescaled.csv
Already rescaled!
Rescaling preference ./results/prefs_A549/A549-Lib5_prefs.csv, using beta of 2.50974
             outfile ./results/prefs_A549/A549-Lib5_prefs_rescaled.csv
Already rescaled!
Rescaling preference ./results/prefs_A549/A549-Lib6_prefs.csv, using beta of 2.50974
             outfile ./results/prefs_A549/A549-Lib6_prefs_rescaled.csv
Already rescaled!
Rescaling preference ./results/prefs_A549/summary_avgprefs.csv, using beta of 2.50974
             outfile ./results/prefs_A549/summary_avgprefs_rescaled.csv
Already rescaled!
Rescaling preference ./results/prefs_CCL141/CCL141-Lib4_prefs.csv, using beta of 2.57381
             outfile ./results/prefs_CCL141/CCL141-Lib4_prefs_rescaled.csv
Already rescaled!
Rescaling preference ./results/prefs_CCL141/CCL141-Lib5_prefs.csv, using beta of 2.57381
             outfile ./results/prefs_CCL141/CCL141-Lib5_prefs_rescaled.csv
Already rescaled!
Rescaling preference ./results/prefs_CCL141/CCL141-Lib6_prefs.csv, using beta of 2.57381
             outfile ./results/prefs_CCL141/CCL141-Lib6_prefs_rescaled.csv
Already rescaled!
Rescaling preference ./results/prefs_CCL141/summary_avgprefs.csv, using beta of 2.57381
             outfile ./results/prefs_CCL141/summary_avgprefs_rescaled.csv
Already rescaled!
```

##### Visualize rescaled preferences¶

In [28]:

```
# Using rescaled preferences

for methoderr, i in [('Bayes2errA549', 'prefs')]:
    for cell in cells:
        outdir = prefsmethoddir[methoderr][cell]
        avgprefs = os.path.join(outdir, 'summary_avgprefs_rescaled.csv')
        logoname = 'logoAvgprefsRescaled'
        print(cell+':')
        print('avgprefs: {0}'.format(avgprefs))
        print('logoname: {0}'.format(logoname))
        print('outdir: {0}'.format(outdir))
        print('overlay1: {0}'.format(wtoverlayfile))
        print('overlay2: {0}'.format(domainsFile))

        log = !dms2_logoplot \
                --prefs {avgprefs} \
                --name {logoname} \
                --outdir {outdir} \
                --overlay1 {wtoverlayfile} wildtype wildtype \
                --overlay2 {domainsFile} D "Domains"\
                --nperline 100 \
                --letterheight 1 \
                --numberevery 10 \
                --use_existing yes

#         print("\n".join(log))
                
        logoplot = os.path.join(outdir, '{0}_prefs.pdf'.format(logoname))
        print('{0}, {1}: {2}'.format(methoderr, cell, logoplot))
        showPDF(logoplot, width=1000)
```

```
A549:
avgprefs: ./results/prefs_A549/summary_avgprefs_rescaled.csv
logoname: logoAvgprefsRescaled
outdir: ./results/prefs_A549
overlay1: ./data/wildtypeoverlayfile.csv
overlay2: ./data/domains.csv
Bayes2errA549, A549: ./results/prefs_A549/logoAvgprefsRescaled_prefs.pdf
```

```
CCL141:
avgprefs: ./results/prefs_CCL141/summary_avgprefs_rescaled.csv
logoname: logoAvgprefsRescaled
outdir: ./results/prefs_CCL141
overlay1: ./data/wildtypeoverlayfile.csv
overlay2: ./data/domains.csv
Bayes2errA549, CCL141: ./results/prefs_CCL141/logoAvgprefsRescaled_prefs.pdf
```

#### Mutational tolerance of PB2¶

I examined entropy across sites of PB2 as a measure of mutational tolerance. I compared entropy of PB2 as measured in A549 and CCL141 cells. I also examined the relationship between mutational tolerance at a site, and variation in natural influenza sequences. (These sequences are as described in jupyter notebook `Fig1_ProcessDMSdata.ipynb`.)

In [29]:

```
# Set matplotlib rcParams for figures
import seaborn as sns
sns.set(context='paper', style='ticks', palette='deep', font='Arial', font_scale=1.042, color_codes=True,
        rc = {'font.size': 10,
 'axes.labelsize': 10,
 'axes.titlesize': 10,
 'xtick.labelsize': 9,
 'ytick.labelsize': 9,
 'legend.fontsize': 9,
 'axes.linewidth': 1.0,
 'grid.linewidth': 0.8,
 'lines.linewidth': 1,
 'lines.markersize': 4.5,
 'patch.linewidth': 0.8,
 'xtick.major.width': 1.0,
 'ytick.major.width': 1.0,
 'xtick.minor.width': 0.8,
 'ytick.minor.width': 0.8,
 'xtick.major.size': 4.5,
 'ytick.major.size': 4.5,
 'xtick.minor.size': 3,
 'ytick.minor.size': 3}
       )
# sns.plotting_context()
```

##### Calculate entropy and neffective from preferences¶

In [30]:

```
entropy_prefsdf = {}
for methoderr, i in [('Bayes2errA549', 'prefs')]:
    for cell in cells:
        outdir = prefsmethoddir[methoderr][cell]
        rescaled_prefsdf_file = os.path.join(outdir, 'summary_avgprefs_rescaled.csv')
        entropy_prefsdf[cell] = dms_tools2.prefs.prefsEntropy(pd.read_csv(rescaled_prefsdf_file), dms_tools2.AAS)
        entropyprefs = os.path.join(outdir, 'summary_avgprefs_entropies_rescaled.csv')
        print(entropyprefs)
        entropy_prefsdf[cell].to_csv(entropyprefs, index=False)

entropydf = (pd.merge(entropy_prefsdf['A549'], entropy_prefsdf['CCL141'], 
                      how='outer', on='site',suffixes=('_A549', '_CCL141'))
            )[['site','entropy_A549','neffective_A549','entropy_CCL141','neffective_CCL141']]
entropydf.head()
```

```
./results/prefs_A549/summary_avgprefs_entropies_rescaled.csv
./results/prefs_CCL141/summary_avgprefs_entropies_rescaled.csv
```

Out[30]:

|  | site | entropy\_A549 | neffective\_A549 | entropy\_CCL141 | neffective\_CCL141 |
| --- | --- | --- | --- | --- | --- |
| 0 | 1 | 0.135966 | 1.145643 | 0.361451 | 1.435411 |
| 1 | 2 | 1.625650 | 5.081722 | 1.455075 | 4.284804 |
| 2 | 3 | 1.835373 | 6.267471 | 0.614140 | 1.848067 |
| 3 | 4 | 1.813682 | 6.132990 | 2.149284 | 8.578718 |
| 4 | 5 | 2.532167 | 12.580745 | 2.396898 | 10.989030 |

##### Calculate entropy and neffective of natural sequences¶

In [31]:

```
IVRseqs = ['Human-PB2', 'Avian-PB2']
naturalfreq = {}
naturalentropy = {}
for strain in IVRseqs:
    alignmentfile = 'data/{0}_subsample_0.fa'.format(strain)
    naturalfreq[strain] = dms_tools2.prefs.aafreqsFromAlignment(alignmentfile, codon_to_aa=True, ignore_gaps=True, ignore_stop=True)
    naturalentropy[strain] = dms_tools2.prefs.prefsEntropy(naturalfreq[strain], dms_tools2.AAS)
    naturalentropy[strain] = naturalentropy[strain][['site', 'entropy', 'neffective']]
naturalentropydf = pd.merge(naturalentropy['Human-PB2'], naturalentropy['Avian-PB2'],
                            how='inner', on='site', suffixes=['_HumanSeasonal','_Avian']
                           )
allentropydf = pd.merge(entropydf, naturalentropydf,
                       how='inner', on='site')
allentropydf.head()
```

Out[31]:

|  | site | entropy\_A549 | neffective\_A549 | entropy\_CCL141 | neffective\_CCL141 | entropy\_HumanSeasonal | neffective\_HumanSeasonal | entropy\_Avian | neffective\_Avian |
| --- | --- | --- | --- | --- | --- | --- | --- | --- | --- |
| 0 | 1 | 0.135966 | 1.145643 | 0.361451 | 1.435411 | 0.0 | 1.0 | 0.000000 | 1.000000 |
| 1 | 2 | 1.625650 | 5.081722 | 1.455075 | 4.284804 | 0.0 | 1.0 | 0.084766 | 1.088462 |
| 2 | 3 | 1.835373 | 6.267471 | 0.614140 | 1.848067 | 0.0 | 1.0 | 0.000000 | 1.000000 |
| 3 | 4 | 1.813682 | 6.132990 | 2.149284 | 8.578718 | 0.0 | 1.0 | 0.000000 | 1.000000 |
| 4 | 5 | 2.532167 | 12.580745 | 2.396898 | 10.989030 | 0.0 | 1.0 | 0.000000 | 1.000000 |

In [32]:

```
df = allentropydf
metrics = [('entropy', 'Entropy'), ('neffective', '$N_{eff}$')]
pairs = [('CCL141', 'A549'),
        ('CCL141', 'Avian'),
        ('A549', 'HumanSeasonal')]
axlabel = {'A549': 'A549 preferences',
          'CCL141': 'CCL141 preferences',
          'Avian': 'Avian sequences',
          'HumanSeasonal': 'Human sequences'}
axesticks = {(('CCL141', 'A549'), 'entropy'): [[0, 1, 2, 3], [0, 1, 2, 3]],
          (('CCL141', 'Avian'), 'entropy'): [[0, 1, 2, 3], [0, 0.25, 0.5, 0.75, 1]],
          (('A549', 'HumanSeasonal'), 'entropy'): [[0, 1, 2, 3], [0, 0.25, 0.5, 0.75, 1]],
          (('CCL141', 'A549'), 'neffective'): [[0, 5, 10, 15, 20], [0, 5, 10, 15, 20]],
          (('CCL141', 'Avian'), 'neffective'): [[0, 5, 10, 15, 20], [1, 2, 3]],
          (('A549', 'HumanSeasonal'), 'neffective'): [[0, 5, 10, 15, 20], [1, 2, 3]],
         }
for (x, y) in pairs:
    for (metric, metriclabel) in metrics:
        g = sns.relplot(x='{0}_{1}'.format(metric, x), y='{0}_{1}'.format(metric, y), data=df,
                        alpha=0.5, height=2.5, aspect=1
                       )
        g.set(xlabel='{0} in {1}'.format(metriclabel, axlabel[x]), 
              ylabel='{0} in {1}'.format(metriclabel, axlabel[y]),
              xticks=axesticks[(x,y),metric][0],
              yticks=axesticks[(x,y),metric][1], 
              )
        r, p = stats.pearsonr(df['{0}_{1}'.format(metric, x)], df['{0}_{1}'.format(metric, y)])
        plt.text(0.05, 1, '$R$ = {0:.2f}'.format(r), horizontalalignment='left',verticalalignment='top', transform=g.ax.transAxes)
        plt.savefig('results/prefs_summaryplots/{0}_{1}_{2}.pdf'.format(metric, x, y), dpi=300, bbox_inches='tight')
```

What is the outlier site that has >0.75 entropy in natural avian sequences? It appears that site 478 has two major variants (V, I) and a minor variant (M).

In [33]:

```
allentropydf[allentropydf['entropy_Avian']>0.75]
```

Out[33]:

|  | site | entropy\_A549 | neffective\_A549 | entropy\_CCL141 | neffective\_CCL141 | entropy\_HumanSeasonal | neffective\_HumanSeasonal | entropy\_Avian | neffective\_Avian |
| --- | --- | --- | --- | --- | --- | --- | --- | --- | --- |
| 477 | 478 | 0.716477 | 2.047209 | 1.231393 | 3.426 | 0.669587 | 1.95343 | 0.765089 | 2.149186 |

In [34]:

```
freqdf = (pd.melt(naturalfreq['Avian-PB2'], id_vars=['site'], 
        value_vars=['A', 'C', 'D', 'E', 'F', 'G', 'H', 'I', 'K', 'L', 'M', 'N', 'P',
       'Q', 'R', 'S', 'T', 'V', 'W', 'Y']))
freqdf[freqdf['site']==478]
```

Out[34]:

|  | site | variable | value |
| --- | --- | --- | --- |
| 477 | 478 | A | 0.000000 |
| 1236 | 478 | C | 0.000000 |
| 1995 | 478 | D | 0.000000 |
| 2754 | 478 | E | 0.000000 |
| 3513 | 478 | F | 0.000000 |
| 4272 | 478 | G | 0.000000 |
| 5031 | 478 | H | 0.000000 |
| 5790 | 478 | I | 0.466667 |
| 6549 | 478 | K | 0.000000 |
| 7308 | 478 | L | 0.000000 |
| 8067 | 478 | M | 0.016667 |
| 8826 | 478 | N | 0.000000 |
| 9585 | 478 | P | 0.000000 |
| 10344 | 478 | Q | 0.000000 |
| 11103 | 478 | R | 0.000000 |
| 11862 | 478 | S | 0.000000 |
| 12621 | 478 | T | 0.000000 |
| 13380 | 478 | V | 0.516667 |
| 14139 | 478 | W | 0.000000 |
| 14898 | 478 | Y | 0.000000 |

#### Examine preferences at sites with known critical residues¶

In [35]:

```
wt_aas = pd.read_csv('data/wildtypeoverlayfile.csv')
criticalresidues = ([1, #Start codon
                     323, 325, 330, 363, 404, 357, 361, 376] + #5' methyl cap
                    list(range(736, 760))) #NLS
criticalresiduesorder = ([1, 2, #Add False site between sets of residues to add space
                     323, 325, 330, 363, 404, 357, 361, 376, 377] +
                    list(range(736, 760)))
rescaledprefsforplot = {}
for cell in cells:
    inprefs = pd.read_csv('./results/prefs_{0}/summary_avgprefs_rescaled.csv'.format(cell))
    rescaledprefsforplot[cell] = (wt_aas.merge(inprefs)
                                  .assign(labeledsite=lambda x: x.wildtype + x.site.astype('str'),
                                          show=lambda x: x['site'].isin(criticalresidues),
                                          siteonly=lambda x: x.site,
                                          site=lambda x: x.labeledsite)
                                 )
    rescaledprefsforplot[cell].siteonly = rescaledprefsforplot[cell].siteonly.astype("category")
    rescaledprefsforplot[cell].siteonly.cat.set_categories(criticalresiduesorder, inplace=True)
    rescaledprefsforplot[cell].sort_values(['siteonly'], inplace=True)
```

In [36]:

```
for cell in cells:
    outfile = os.path.join(prefs_summaryplotsdir, 'criticalresidues_rescaledprefs{0}.pdf'.format(cell))
    dms_tools2.rplot.siteSubsetGGSeqLogo(
            rescaledprefsforplot[cell],
            AAS,
            outfile,
            width=0.25 * len(criticalresidues),
            height=1.75)
    print('Rescaled preferences as measured in {0}, at sites with critical residues:'.format(cell))
    showPDF(outfile, width=500)
```

```
Rescaled preferences as measured in A549, at sites with critical residues:
```

```
Rescaled preferences as measured in CCL141, at sites with critical residues:
```

#### Copy files to paper figures directory¶

In [37]:

```
myfiguresdir = os.path.join(figuresdir, 'Fig2/')
if not os.path.isdir(myfiguresdir):
    os.mkdir(myfiguresdir)

files = (['./results/prefs_summaryplots/prefsBayes2errA549_prefscorr.pdf',
         './results/prefs_A549/logoAvgprefsRescaled_prefs.pdf',
         './results/prefs_CCL141/logoAvgprefsRescaled_prefs.pdf'] +
         ['results/prefs_summaryplots/{0}_{1}_{2}.pdf'.format(metric, x, y) 
          for metric in ['entropy', 'neffective'] for (x,y) in pairs] +
         ['results/prefs_summaryplots/criticalresidues_rescaledprefs{0}.pdf'.format(cell)
         for cell in cells] +
         ['./results/phydms_analysis/default_HumanAvian-PB2_subsample_0/modelcomparison.md']
        )
for f in files:
    shutil.copy(f, myfiguresdir)
    
filestorename = [('./results/prefs_A549/logoAvgprefsRescaled_prefs.pdf', 'A549_logoAvgprefsRescaled_prefs.pdf'),
                 ('./results/prefs_CCL141/logoAvgprefsRescaled_prefs.pdf', 'CCL141_logoAvgprefsRescaled_prefs.pdf'),
                 ('./results/prefs_A549/summary_avgprefs_entropies_rescaled.csv', 'A549_summary_avgprefs_entropies_rescaled.csv'),
                 ('./results/prefs_CCL141/summary_avgprefs_entropies_rescaled.csv', 'CCL141_summary_avgprefs_entropies_rescaled.csv')
                ]
for f1, f2 in filestorename:
    shutil.copy(f1, os.path.join(myfiguresdir, f2))
```
