## Supplementary material for "Comprehensive mapping of avian influenza polymerase adaptation to the human host": File S2: Fig3_IdentifyAdaptiveMutations.html


### Infer differential selection¶

##### Import modules, define directories¶

In [2]:

```
import os
import re
import glob
import shutil
import subprocess
import string
import warnings
warnings.simplefilter('ignore') # don't print warnings to avoid clutter in notebook

import pandas as pd
import numpy as np
import matplotlib as mpl
import matplotlib.pyplot as plt
%matplotlib inline

import dms_tools2
import dms_tools2.sra
import dms_tools2.plot
import dms_tools2.prefs
# import dms_tools2.compareprefs
import dms_tools2.rplot
print(dms_tools2.rplot.versionInfo())
from dms_tools2.ipython_utils import showPDF
from dms_tools2 import AAS

from pymodules.utils import permtest

from IPython.display import display, HTML, Markdown, Image

print("Using dms_tools2 version {0}".format(dms_tools2.__version__))

# CPUs to use, should not exceed the number you request with slurm
ncpus = 14

# do we use existing results or generate everything new?
use_existing = 'yes'

# Directories

fastqdir = './fastq/'
datadir = './data'
resultsdir = './results/' 
countsdir = os.path.join(resultsdir, 'codoncounts/')
countsplotprefix = os.path.join(countsdir, 'summary')

cells = ['A549', 'CCL141']
prefsmethod = [('Bayes2err', 'prefs1'), ('Bayes2errA549', 'prefs')]
prefsdir = {cell: os.path.join(resultsdir, 'prefs_{0}'.format(cell)) for cell in cells}
prefsdir1 = {cell: os.path.join(resultsdir, 'prefs1_{0}'.format(cell)) for cell in cells}
prefsmethoddir = {'Bayes2errA549': prefsdir, #method Bayesian, with A549-WT as the post-err control, because CCL-141WT has high mutation rate
                  'Bayes2err': prefsdir1} #method Bayesian, with respective host cell WT post-err ctrls
# phydmsdir = os.path.join(resultsdir, 'phydms_analysis/')
# prefs_summaryplotsdir= os.path.join(resultsdir, 'prefs_summaryplots/')
diffseldir = os.path.join(resultsdir, 'diffsel')
diffselprefix = os.path.join(diffseldir, 'summary_')
# prefsdistdir = os.path.join(resultsdir, 'prefsdist')

paperdir = './paper'
figuresdir = os.path.join(paperdir, 'figures/')

for xdir in ([fastqdir, datadir, resultsdir, countsdir] +
             list(prefsdir.values()) + list(prefsdir1.values()) +
#              [phydmsdir, prefs_summaryplotsdir] +
             [diffseldir] +
             [paperdir, figuresdir]
             ):
    if not os.path.isdir(xdir):
        os.mkdir(xdir)
```

```
Using `rpy2` 2.9.1 with ``R`` 4.3
Using dms_tools2 version 2.3.0
```

##### Define samples¶

In [5]:

```
countsbatchfile = os.path.join(countsdir, 'batch.csv')
samples = pd.read_csv(countsbatchfile)
display(HTML(samples.to_html(index=False)))
```

| name | R1 |
| --- | --- |
| A549-Lib4 | A549-MutVirusLib-Rep1\_R1.fastq.gz |
| A549-Lib5 | A549-MutVirusLib-Rep2\_R1.fastq.gz |
| A549-Lib6 | A549-MutVirusLib-Rep3\_R1.fastq.gz |
| A549-WT | A549-WT\_R1.fastq.gz |
| CCL141-Lib4 | CCL141-MutVirusLib-Rep1\_R1.fastq.gz |
| CCL141-Lib5 | CCL141-MutVirusLib-Rep2\_R1.fastq.gz |
| CCL141-Lib6 | CCL141-MutVirusLib-Rep3\_R1.fastq.gz |
| CCL141-WT | CCL141-WT\_R1.fastq.gz |
| DNA-Lib4 | DNA-Lib-Rep1\_R1.fastq.gz |
| DNA-Lib5 | DNA-Lib-Rep2\_R1.fastq.gz |
| DNA-Lib6 | DNA-Lib-Rep3\_R1.fastq.gz |
| DNA-WT | DNA-WT\_R1.fastq.gz |

#### Differential selection¶

##### Calculate differential selection¶

We will quantify the differential selection on mutations using the dms2\_diffsel program. First, we first create a batch file to use with dms2\_batch\_diffsel. By grouping replicates for the same timepoint in the batch file, we tell dms2\_batch\_diffsel to analyze these together and take their mean and median.

Note: dms2\_diffsel takes only a single error control. I used the `A549-WT` sample as error control here, since it should correct for both RT-PCR and PCR errors.

In [6]:

```
diffselbatchfile = os.path.join(diffseldir, 'batch.csv')

diffselbatch = pd.DataFrame(
        columns=['group', 'name', 'sel', 'mock', 'err'],
        data=[('A549vCCL141', 'Lib4', 'A549-Lib4', 'CCL141-Lib4', 'A549-WT'),
              ('A549vCCL141', 'Lib5', 'A549-Lib5', 'CCL141-Lib5', 'A549-WT'),
              ('A549vCCL141', 'Lib6', 'A549-Lib6', 'CCL141-Lib6', 'A549-WT')]
)

print("Here is the batch file that we write to CSV format to use as input:")
display(HTML(diffselbatch.to_html(index=False)))

diffselbatch.to_csv(diffselbatchfile, index=False)
```

```
Here is the batch file that we write to CSV format to use as input:
```

| group | name | sel | mock | err |
| --- | --- | --- | --- | --- |
| A549vCCL141 | Lib4 | A549-Lib4 | CCL141-Lib4 | A549-WT |
| A549vCCL141 | Lib5 | A549-Lib5 | CCL141-Lib5 | A549-WT |
| A549vCCL141 | Lib6 | A549-Lib6 | CCL141-Lib6 | A549-WT |

Run dms2\_batch\_diffsel. Input counts are as calculated by dms2\_batch\_bcsubamp.

In [7]:

```
cmd = 'dms2_batch_diffsel --summaryprefix summary --batchfile {0} --outdir {1} --indir {2} --use_existing {3}'\
    .format(diffselbatchfile, diffseldir, countsdir, use_existing)
print(cmd)
!$cmd
```

```
dms2_batch_diffsel --summaryprefix summary --batchfile ./results/diffsel/batch.csv --outdir ./results/diffsel --indir ./results/codoncounts/ --use_existing yes
INFO:dms2_batch_diffsel:Beginning execution of dms2_batch_diffsel in directory /fh/fast/bloom_j/computational_notebooks/yqsoh/2018/PB2-DMS

INFO:dms2_batch_diffsel:Progress is being logged to ./results/diffsel/summary.log
INFO:dms2_batch_diffsel:Version information:
	Time and date: Fri Dec 28 10:52:10 2018
	Platform: Linux-3.13.0-143-generic-x86_64-with-debian-jessie-sid
	Python version: 3.6.5 |Anaconda, Inc.| (default, Apr 29 2018, 16:14:56)  [GCC 7.2.0]
	dms_tools2 version: 2.3.0
	Bio version: 1.72
	HTSeq version: 0.11.0
	pandas version: 0.23.4
	numpy version: 1.15.1
	IPython version: 6.5.0
	jupyter version: 1.0.0
	matplotlib version: 3.0.0
	plotnine version: 0.5.0
	natsort version: 5.4.1
	pystan version: 2.16.0.0
	scipy version: 1.1.0
	seaborn version: 0.9.0
	phydmslib version: 2.3.1
	statsmodels version: 0.9.0
	rpy2 version: 2.9.1
	regex version: 2.4.151
	umi_tools version unknown

INFO:dms2_batch_diffsel:Parsed the following arguments:
	outdir = ./results/diffsel
	ncpus = -1
	use_existing = yes
	indir = ./results/codoncounts/
	chartype = codon_to_aa
	excludestop = yes
	pseudocount = 5
	mincount = 0
	batchfile = ./results/diffsel/batch.csv
	summaryprefix = summary

INFO:dms2_batch_diffsel:Parsing info from ./results/diffsel/batch.csv
INFO:dms2_batch_diffsel:Read the following sample information:
group,name,sel,mock,err
A549vCCL141,Lib4,A549-Lib4,CCL141-Lib4,A549-WT
A549vCCL141,Lib5,A549-Lib5,CCL141-Lib5,A549-WT
A549vCCL141,Lib6,A549-Lib6,CCL141-Lib6,A549-WT


INFO:dms2_batch_diffsel:Output summary files already exist and '--use_existing' is 'yes', so exiting with no further action.
```

We obtain output files giving the mutation and site differential selection values in the formats of the `mutdiffsel.csv` and `sitediffsel.csv` files created by dms2\_diffsel. Specifically, there is a file for each individual sample, with a file name that gives the `group` and `name` specified for this sample in `--batchfile`:

In [8]:

```
!ls {diffseldir}/A549vCCL141*.csv
```

```
./results/diffsel/A549vCCL141-Lib4_mutdiffsel.csv
./results/diffsel/A549vCCL141-Lib4_sitediffsel.csv
./results/diffsel/A549vCCL141-Lib5_mutdiffsel.csv
./results/diffsel/A549vCCL141-Lib5_sitediffsel.csv
./results/diffsel/A549vCCL141-Lib6_mutdiffsel.csv
./results/diffsel/A549vCCL141-Lib6_sitediffsel.csv
```

There are also files giving the mean and median differential selection for **all** samples in each group.
These files are prefixed with the name given by `--summaryprefix` (in this case, `summary`), and also indicate the group:

In [9]:

```
!ls {diffseldir}/summary*A549vCCL141*.csv
```

```
./results/diffsel/summary_A549vCCL141-meanmutdiffsel.csv
./results/diffsel/summary_A549vCCL141-meansitediffsel.csv
./results/diffsel/summary_A549vCCL141-medianmutdiffsel.csv
./results/diffsel/summary_A549vCCL141-mediansitediffsel.csv
```

We can look at the correlation of samples **within** the group for mutation differential selection, positive site differential selection, absolute site differential selection, and maximum mutation differential selection for a site.

In [10]:

```
seltypes = ['mutdiffsel', 'positivesitediffsel', 'absolutesitediffsel', 'maxmutdiffsel']
groups = diffselbatch['group'].unique()
plots = []
for seltype in seltypes:
    for g in groups:
        plot = diffselprefix + g + '-' + seltype + 'corr.pdf'
        plots.append(plot)
print('\t'+ '\t\t'.join(seltypes))
showPDF(plots, width=1000)
```

```
	mutdiffsel		positivesitediffsel		absolutesitediffsel		maxmutdiffsel
```

Now we look at the differential selection along the primary sequence.
Several plots showing this kind of selection are made by dms2\_batch\_diffsel.
These plots show either the **mean** or the **median** differential selection among all samples in each group (CSV files have been created containing this differential selection as described above).

These files all have the prefix indicated by `--summaryprefix`, and show the positive differential selection, total site differential selection (both positive and negative), the maximum differential selection, and the minimum and maximum differential selection.
Here are all the files created by this analysis:

In [11]:

```
!ls {diffseldir}/summary*diffsel.pdf
```

```
./results/diffsel/summary_meanmaxdiffsel.pdf
./results/diffsel/summary_meanminmaxdiffsel.pdf
./results/diffsel/summary_meanpositivediffsel.pdf
./results/diffsel/summary_meantotaldiffsel.pdf
./results/diffsel/summary_medianmaxdiffsel.pdf
./results/diffsel/summary_medianminmaxdiffsel.pdf
./results/diffsel/summary_medianpositivediffsel.pdf
./results/diffsel/summary_mediantotaldiffsel.pdf
```

##### Visualize preferences¶

Below are the mean and median side-by-side for the total site differential selection: total differential selection for **all mutations combined at a given site**. For this experiment, the median and mean values look very similar.

In [12]:

```
seltypes = ['totaldiffsel', 'positivediffsel', 'minmaxdiffsel', 'maxdiffsel']
for seltype in seltypes:
    print(seltype + ': mean, median')
    showPDF(['{0}mean{1}.pdf'.format(diffselprefix, seltype), '{0}median{1}.pdf'.format(diffselprefix, seltype)], width=1000)
```

```
totaldiffsel: mean, median
```

```
positivediffsel: mean, median
```

```
minmaxdiffsel: mean, median
```

```
maxdiffsel: mean, median
```

We can visualize the differential selection values for each mutation in the form of logo plots, that can be created with dms2\_logoplot. We make those logo plots using the **median** and **mean** mutation differential selection values returned by dms2\_batch\_diffsel.

In [13]:

```
wtoverlayfile = os.path.join(datadir, 'wildtypeoverlayfile.csv')
```

In [14]:

```
diffseltype = ['median', 'mean']
for m in diffseltype:
    outdir = diffseldir
    diffselfile = '{0}A549vCCL141-{1}mutdiffsel.csv'.format(diffselprefix, m)
    logoname = 'logoDiffsel{0}'.format(m)
    print('**Creating logoplot for {0}mutdiffsel'.format(m))
    print('diffsel: {0}'.format(diffselfile))
    print('logoname: {0}'.format(logoname))
    print('outdir: {0}'.format(diffseldir))
    print('overlay1: {0}'.format(wtoverlayfile))
    log = !dms2_logoplot \
            --diffsel {diffselfile} \
            --name {logoname} \
            --outdir {diffseldir} \
            --restrictdiffsel all \
            --sepline yes \
            --nperline 100 \
            --letterheight 1 \
            --mapmetric functionalgroup \
            --overlay1 {wtoverlayfile} wildtype wildtype \
            --underlay yes \
            --scalebar 10 diffsel=10 \
            --use_existing {use_existing} 
#     print("\n".join(log))
    logoplot = os.path.join(diffseldir, '{0}_diffsel.pdf'.format(logoname))
    print(logoplot)
    showPDF(logoplot, width=1000)
```

```
**Creating logoplot for medianmutdiffsel
diffsel: ./results/diffsel/summary_A549vCCL141-medianmutdiffsel.csv
logoname: logoDiffselmedian
outdir: ./results/diffsel
overlay1: ./data/wildtypeoverlayfile.csv
./results/diffsel/logoDiffselmedian_diffsel.pdf
```

```
**Creating logoplot for meanmutdiffsel
diffsel: ./results/diffsel/summary_A549vCCL141-meanmutdiffsel.csv
logoname: logoDiffselmean
outdir: ./results/diffsel
overlay1: ./data/wildtypeoverlayfile.csv
./results/diffsel/logoDiffselmean_diffsel.pdf
```

#### Identify top adaptive mutations¶

In [15]:

```
## Set plotting parameters/preferences
# Set colors to use throughout
palette ={"None":"gray",
          "Human":"#d95f02", "A549":"#d95f02", "Known - Human":"#d95f02", "H7N9 - Human":"#d95f02",
          "Bird":"#1b9e77", "Avian":"#1b9e77", "CCL141":"#1b9e77", 
          "Both":"#7570b3", "Known - Both":"#7570b3",
          "Yes": "#d95f02", "No": "gray" #KnownAdaptive
         }
# Set matplotlib rcParams for figures
import seaborn as sns
sns.set(context='paper', style='ticks', palette='deep', font='Arial', font_scale=1.042, color_codes=True,
        rc = {'font.size': 10,
 'axes.labelsize': 10,
 'axes.titlesize': 10,
 'xtick.labelsize': 9,
 'ytick.labelsize': 9,
 'legend.fontsize': 9,
 'axes.linewidth': 1.0,
 'grid.linewidth': 0.8,
 'lines.linewidth': 1,
 'lines.markersize': 4.5,
 'patch.linewidth': 0.8,
 'xtick.major.width': 1.0,
 'ytick.major.width': 1.0,
 'xtick.minor.width': 0.8,
 'ytick.minor.width': 0.8,
 'xtick.major.size': 4.5,
 'ytick.major.size': 4.5,
 'xtick.minor.size': 3,
 'ytick.minor.size': 3}
       )
# sns.plotting_context()
```

##### Gather prior annotations and experimental data¶

In [16]:

```
# Gather prior annotations for known human/mammalian adaptive mutations
mutsAdaptive = pd.read_table('data/Muts_HumanAvian.txt')
mutsAdaptive['KnownAdaptive-ExptOrComp'] = 'Yes'
mutsAdaptive['aa_Avian'] = mutsAdaptive['Mutation'].str.extract('(^[A-Z])')
mutsAdaptive['aa_Human'] = mutsAdaptive['Mutation'].str.extract('([A-Z]$)')
mutsAdaptive.rename(columns={'ExptVerified': 'KnownAdaptive'}, inplace=True)
mutsAdaptive_mut = (mutsAdaptive[['Site', 'KnownAdaptive', 'KnownAdaptive-ExptOrComp', 
                                 'aa_Avian', 'aa_Human']]
                    .drop_duplicates() )
display(HTML(mutsAdaptive_mut.head().to_html(index=False)))
```

| Site | KnownAdaptive | KnownAdaptive-ExptOrComp | aa\_Avian | aa\_Human |
| --- | --- | --- | --- | --- |
| 9 | Yes | Yes | D | N |
| 44 | No | Yes | A | S |
| 64 | No | Yes | M | T |
| 67 | No | Yes | I | V |
| 74 | Yes | Yes | G | S |

In [17]:

```
x = mutsAdaptive_mut[mutsAdaptive['KnownAdaptive']=='Yes']['Site']
print(set(list(x)))
print(len(set(list(x))))
```

```
{256, 9, 526, 271, 660, 534, 535, 158, 286, 684, 701, 189, 702, 192, 199, 74, 714, 588, 591, 740, 489, 627, 636, 253}
24
```

In [18]:

```
# Gather experimental measurements of 
# preferences, mutational effect, and differential selection,
# and merge them into a dataframe for downstream plotting.

# Preferences
pfiles = [os.path.join(prefsmethoddir['Bayes2errA549'][cell], 'summary_avgprefs.csv') 
          for cell in cells]
plist = []
for i, f in enumerate(pfiles):
    df = pd.read_csv(f)
    df = pd.melt(df, id_vars=['site'], value_vars=dms_tools2.AAS, 
                var_name='mutation', value_name='pref{0}'.format(cells[i]))
    df['log2pref{0}'.format(cells[i])] = np.log2(df['pref{0}'.format(cells[i])])
    plist.append(df)
pref = pd.merge(plist[0], plist[1], how='left', on=['site', 'mutation'], sort=False)

# Mutational effect
efiles = [os.path.join(prefsmethoddir['Bayes2errA549'][cell], 'summary_avgmuteffectwt.csv') 
          for cell in cells]
elist = []
for i, f in enumerate(efiles):
    df = pd.read_csv(f)
    df['mutation'] = df['final']
    df.drop(columns=['wildtype','initial','final'], inplace=True)
    elist.append(df)
effect = pd.merge(elist[0], elist[1], how='left', on=['site', 'mutation'], sort=False, 
                  suffixes=(cells[0], cells[1]))

# Differential selection
diffsel_mut = pd.read_csv(diffselprefix+'A549vCCL141-meanmutdiffsel.csv')

dfmut = ((((pd.merge(pref, effect, how='left', on=['site', 'mutation'], sort=False) )
          .merge(diffsel_mut, how='left', on=['site', 'mutation'], sort=False) )
         .merge(mutsAdaptive_mut, how='left', 
                 left_on=['site', 'mutation'], right_on=['Site', 'aa_Human'], sort=False)
         .drop(columns=['Site', 'aa_Avian', 'aa_Human', 'KnownAdaptive-ExptOrComp']) )
         .fillna({'KnownAdaptive': 'No'}) )

dfmut = dfmut[['site', 'wildtype', 'mutation', 
               'prefA549', 'prefCCL141', 'log2prefA549', 'log2prefCCL141', 
               'effectA549', 'effectCCL141', 'log2effectA549', 'log2effectCCL141', 
               'mutdiffsel', 'KnownAdaptive']]

dfmut.head()
```

Out[18]:

|  | site | wildtype | mutation | prefA549 | prefCCL141 | log2prefA549 | log2prefCCL141 | effectA549 | effectCCL141 | log2effectA549 | log2effectCCL141 | mutdiffsel | KnownAdaptive |
| --- | --- | --- | --- | --- | --- | --- | --- | --- | --- | --- | --- | --- | --- |
| 0 | 1 | M | A | 0.018441 | 0.035779 | -5.760954 | -4.804750 | 0.037634 | 0.098789 | -4.731825 | -3.339500 | -0.596582 | No |
| 1 | 2 | E | A | 0.017338 | 0.030416 | -5.849903 | -5.039008 | 0.093657 | 0.195126 | -3.416476 | -2.357521 | -0.642891 | No |
| 2 | 3 | R | A | 0.029029 | 0.022867 | -5.106354 | -5.450617 | 0.172319 | 0.077507 | -2.536850 | -3.689531 | 0.629427 | No |
| 3 | 4 | I | A | 0.033297 | 0.049470 | -4.908455 | -4.337306 | 0.269836 | 0.486289 | -1.889844 | -1.040116 | -0.681969 | No |
| 4 | 5 | K | A | 0.062883 | 0.041191 | -3.991193 | -4.601543 | 0.766084 | 0.390159 | -0.384426 | -1.357867 | 0.610305 | No |

##### Do known adaptive mutations have higher differential selection compared to other mutations?¶

Do known adaptive mutations have higher differential selection values than all other mutations? Yes. but they are not the highest I found in my experiments.

Note: p-value calculated by a permutation test (using `permtest` imported from `pymodules/utils.py`). The p value is then defined as, the fraction of times the difference in medians is as large as observed, for randomly shuffled mutdiffsel values.

In [19]:

```
dataframe = dfmut[pd.notnull(dfmut['mutdiffsel'])] #Need to remove NAs where mutated aa = wt aa
group = 'KnownAdaptive'
(metric, name) = ('mutdiffsel', 'Mutation diff sel')

# Calculate p-value
(p, eq) = permtest(dataframe, metric, group, 10000)
print('{0} / {1}: '.format(metric, name))
print('  p {0} {1}'.format(eq, p))
```

```
mutdiffsel / Mutation diff sel: 
  p <= 0.0001
```

In [20]:

```
sns.set_style("ticks")
fig, axes = plt.subplots(1, 1, figsize=(2.5,3))
g = sns.boxplot(x=group, y=metric, data=dataframe, #color='gray',
                palette=palette, 
                ax=axes, order=['Yes', 'No'])
g.set_ylabel(name)
g.set_xlabel('Known adaptive mutation')
sns.despine()
# Adding p value annotation
x1, x2 = 0, 1   # first column: 0, see plt.xticks()
yrange = dataframe[metric].max() - dataframe[metric].min()
y, h, col = dataframe[metric].max() + yrange/10, yrange/50, 'k'
g.plot([x1, x1, x2, x2], [y, y+h, y+h, y], lw=0.5, c=col)
g.text((x1+x2)*.5, y+h*2, 'p {0} {1}'.format(eq, p), 
         ha='center', va='bottom', color=col) 
# plt.tight_layout(pad=1, w_pad=1, h_pad=1.0)
plt.tight_layout()
fig.savefig('results/diffsel/knownadapt_mutdiffsel_boxplot.pdf', dpi=300, bbox_inches='tight')
```

##### Are known adaptive mutations the top most differentially selected mutations?¶

I plotted the top 50 most differentially selected mutations and only 2 of those are known. i.e. many top differentially selected mutations are novel.

In [21]:

```
sns.set_style("whitegrid")
metric, name = ('mutdiffsel', 'Mutation diff sel')
sub = dfmut.sort_values(by=metric, ascending=False).head(n=50)
sub['rank'] = range(len(sub))
sub['sitemut'] = sub['wildtype'] + sub['site'].astype(str) + ' ' + sub['mutation']
fig, axes = plt.subplots(1, 1, figsize=(8,3))
sub1 = sub[sub['KnownAdaptive']=='No']
sub2 = sub[sub['KnownAdaptive']=='Yes']
axes.scatter(x=sub1['rank'], y=sub1[metric], 
             s=20, c=palette['No'], marker='o')
axes.scatter(x=sub2['rank'], y=sub2[metric], 
             s=50, c=palette['Yes'], marker='x')
axes.set_ylabel(name)
axes.set_xlabel('')
axes.grid(False)
axes.set_xticks(sub['rank'])
axes.set_xticklabels(sub['sitemut']);
axes.tick_params(axis='x',rotation=90,)
plt.tight_layout()
fig.savefig('results/diffsel/knownadapt_mutdiffsel_rankplot.pdf', dpi=300, bbox_inches='tight')
```

##### Identify novel top adaptive mutations¶

I will identify novel adaptive mutations by a combination of criteria:

- Human adaptive mutations: high positive differential selection, high mutational effect in A549
- Avian adaptive mutations: high negative differential selection, high mutational effect in CCL141
- Adaptive in both: high mutational effect in both A549 and CCL141

I required that in addition to high differential selection, mutational effect is also high (perform well compared to the wild-type residue), because the mutations that fix in nature are likely the ones that are *both* host adaptive and confer high fitness.

The thresholds I have set for each of these criteria are arbitrary -- they are mostly to get the number of mutations for each category down to a manageable number.

In [22]:

```
def select_muts(row):
    if row['log2effectA549']>1 and row['log2effectCCL141']>1:
        return 'Both'
    elif row['log2effectA549']>1 and row['mutdiffsel']>1.5:
        return 'A549'
    elif row['log2effectCCL141']>1 and row['mutdiffsel']<-2:
        return 'CCL141'
    else:
        return 'None'
dfmut['Experimentally adaptive in'] = dfmut.apply(lambda row: select_muts(row), axis=1)
dfmut.rename(columns={'KnownAdaptive': 'Known human adaptive'}, inplace=True)
dfmut.to_csv('results/diffsel/summary_prefs_effects_diffsel.csv', index=False)
display(HTML(dfmut.head().to_html()))
display(HTML(dfmut.groupby('Experimentally adaptive in').describe().to_html()))
```

|  | site | wildtype | mutation | prefA549 | prefCCL141 | log2prefA549 | log2prefCCL141 | effectA549 | effectCCL141 | log2effectA549 | log2effectCCL141 | mutdiffsel | Known human adaptive | Experimentally adaptive in |
| --- | --- | --- | --- | --- | --- | --- | --- | --- | --- | --- | --- | --- | --- | --- |
| 0 | 1 | M | A | 0.018441 | 0.035779 | -5.760954 | -4.804750 | 0.037634 | 0.098789 | -4.731825 | -3.339500 | -0.596582 | No | None |
| 1 | 2 | E | A | 0.017338 | 0.030416 | -5.849903 | -5.039008 | 0.093657 | 0.195126 | -3.416476 | -2.357521 | -0.642891 | No | None |
| 2 | 3 | R | A | 0.029029 | 0.022867 | -5.106354 | -5.450617 | 0.172319 | 0.077507 | -2.536850 | -3.689531 | 0.629427 | No | None |
| 3 | 4 | I | A | 0.033297 | 0.049470 | -4.908455 | -4.337306 | 0.269836 | 0.486289 | -1.889844 | -1.040116 | -0.681969 | No | None |
| 4 | 5 | K | A | 0.062883 | 0.041191 | -3.991193 | -4.601543 | 0.766084 | 0.390159 | -0.384426 | -1.357867 | 0.610305 | No | None |

|  | effectA549 | | | | | | | | effectCCL141 | | | | | | | | log2effectA549 | | | | | | | | log2effectCCL141 | | | | | | | | log2prefA549 | | | | | | | | log2prefCCL141 | | | | | | | | mutdiffsel | | | | | | | | prefA549 | | | | | | | | prefCCL141 | | | | | | | | site | | | | | | | |
| --- | --- | --- | --- | --- | --- | --- | --- | --- | --- | --- | --- | --- | --- | --- | --- | --- | --- | --- | --- | --- | --- | --- | --- | --- | --- | --- | --- | --- | --- | --- | --- | --- | --- | --- | --- | --- | --- | --- | --- | --- | --- | --- | --- | --- | --- | --- | --- | --- | --- | --- | --- | --- | --- | --- | --- | --- | --- | --- | --- | --- | --- | --- | --- | --- | --- | --- | --- | --- | --- | --- | --- | --- | --- | --- | --- | --- | --- | --- | --- | --- |
|  | count | mean | std | min | 25% | 50% | 75% | max | count | mean | std | min | 25% | 50% | 75% | max | count | mean | std | min | 25% | 50% | 75% | max | count | mean | std | min | 25% | 50% | 75% | max | count | mean | std | min | 25% | 50% | 75% | max | count | mean | std | min | 25% | 50% | 75% | max | count | mean | std | min | 25% | 50% | 75% | max | count | mean | std | min | 25% | 50% | 75% | max | count | mean | std | min | 25% | 50% | 75% | max | count | mean | std | min | 25% | 50% | 75% | max |
| Experimentally adaptive in |  |  |  |  |  |  |  |  |  |  |  |  |  |  |  |  |  |  |  |  |  |  |  |  |  |  |  |  |  |  |  |  |  |  |  |  |  |  |  |  |  |  |  |  |  |  |  |  |  |  |  |  |  |  |  |  |  |  |  |  |  |  |  |  |  |  |  |  |  |  |  |  |  |  |  |  |  |  |  |  |
| A549 | 34.0 | 2.769773 | 0.740277 | 2.019931 | 2.230899 | 2.460975 | 3.158564 | 4.677888 | 34.0 | 0.506684 | 0.252159 | 0.098527 | 0.333353 | 0.506312 | 0.700736 | 1.117994 | 34.0 | 1.425838 | 0.350339 | 1.014306 | 1.157530 | 1.299219 | 1.659010 | 2.225857 | 34.0 | -1.228273 | 0.961324 | -3.343339 | -1.584886 | -0.982106 | -0.513206 | 0.160912 | 34.0 | -3.048108 | 0.586045 | -4.047004 | -3.547981 | -3.001045 | -2.689285 | -1.798817 | 34.0 | -4.521246 | 0.784575 | -6.117086 | -4.779421 | -4.395836 | -3.966782 | -3.143583 | 34.0 | 2.165375 | 0.554717 | 1.539068 | 1.754101 | 2.004553 | 2.390083 | 3.603195 | 34.0 | 0.131361 | 0.057855 | 0.060497 | 0.085544 | 0.124941 | 0.155045 | 0.287410 | 34.0 | 0.049739 | 0.025308 | 0.014407 | 0.036413 | 0.047517 | 0.064004 | 0.113158 | 34.0 | 394.205882 | 235.131141 | 9.0 | 177.5 | 521.0 | 603.25 | 701.0 |
| Both | 7.0 | 3.044897 | 0.477527 | 2.231670 | 2.749195 | 3.327202 | 3.383925 | 3.489166 | 7.0 | 2.419446 | 0.366403 | 2.001495 | 2.257697 | 2.281910 | 2.494686 | 3.147953 | 7.0 | 1.589763 | 0.242014 | 1.158124 | 1.457814 | 1.734309 | 1.758698 | 1.802882 | 7.0 | 1.261588 | 0.205964 | 1.001078 | 1.174843 | 1.190242 | 1.317849 | 1.654414 | 7.0 | -2.919792 | 0.289175 | -3.288854 | -3.174998 | -2.806211 | -2.716303 | -2.560880 | 7.0 | -3.310580 | 0.364543 | -3.953988 | -3.476981 | -3.252818 | -3.058549 | -2.896194 | 7.0 | 0.239691 | 0.326200 | -0.245348 | 0.078703 | 0.187155 | 0.396075 | 0.786476 | 7.0 | 0.134402 | 0.026252 | 0.102319 | 0.110864 | 0.142970 | 0.152164 | 0.169472 | 7.0 | 0.103455 | 0.024485 | 0.064525 | 0.089810 | 0.104907 | 0.120402 | 0.134326 | 7.0 | 140.571429 | 88.988496 | 60.0 | 77.5 | 82.0 | 197.00 | 293.0 |
| CCL141 | 42.0 | 0.399170 | 0.153844 | 0.152611 | 0.287264 | 0.397510 | 0.447976 | 0.848517 | 42.0 | 3.527602 | 2.003748 | 2.022795 | 2.526134 | 2.796926 | 4.119619 | 14.150783 | 42.0 | -1.430785 | 0.571774 | -2.712068 | -1.799853 | -1.331055 | -1.158524 | -0.236985 | 42.0 | 1.689004 | 0.556303 | 1.016350 | 1.336931 | 1.483785 | 2.042374 | 3.822810 | 42.0 | -4.628683 | 0.649272 | -6.284928 | -4.976986 | -4.516553 | -4.292647 | -3.214398 | 42.0 | -3.702863 | 0.629184 | -4.780023 | -4.148971 | -3.896101 | -3.263103 | -2.253953 | 42.0 | -2.570830 | 0.437346 | -3.913665 | -2.786036 | -2.445689 | -2.266236 | -2.002067 | 42.0 | 0.044313 | 0.018848 | 0.012825 | 0.031753 | 0.043693 | 0.051025 | 0.107738 | 42.0 | 0.084770 | 0.041597 | 0.036397 | 0.056373 | 0.067168 | 0.104168 | 0.209649 | 42.0 | 359.214286 | 71.829587 | 60.0 | 360.0 | 377.0 | 380.00 | 472.0 |
| None | 15097.0 | 0.342297 | 0.381177 | 0.008920 | 0.074567 | 0.176643 | 0.485445 | 5.516999 | 15097.0 | 0.393223 | 0.396989 | 0.006728 | 0.091110 | 0.237625 | 0.595687 | 5.043287 | 15097.0 | -2.401766 | 1.625693 | -6.808795 | -3.745327 | -2.501093 | -1.042619 | 2.463884 | 15097.0 | -2.132660 | 1.613767 | -7.215523 | -3.456249 | -2.073243 | -0.747373 | 2.334364 | 15097.0 | -4.869102 | 1.195911 | -10.265982 | -5.685631 | -4.877189 | -4.164855 | -0.572787 | 15097.0 | -4.807451 | 1.165324 | -10.803930 | -5.574908 | -4.724838 | -4.091779 | -0.609374 | 14338.0 | -0.158069 | 0.682177 | -3.636944 | -0.515365 | -0.099973 | 0.183754 | 3.229887 | 15097.0 | 0.049793 | 0.058197 | 0.000812 | 0.019429 | 0.034027 | 0.055751 | 0.672317 | 15097.0 | 0.049879 | 0.052352 | 0.000559 | 0.020979 | 0.037817 | 0.058648 | 0.655481 | 15097.0 | 380.136848 | 219.334264 | 1.0 | 190.0 | 381.0 | 570.00 | 759.0 |

Plot where these top mutations fall on metrics of diffsel and muteffect:

In [23]:

```
sns.set_style("ticks")
sub = dfmut

hues = ['Known human adaptive', 'Experimentally adaptive in']
hue_order = {'Known human adaptive': ['No', 'Yes'],
             'Experimentally adaptive in': ['A549', 'CCL141', 'Both', 'None'],
            }
hue_kws = {'Known human adaptive': dict(alpha=[0.2,1], facecolors=[None,None], s=[10,30], linewidths=[1,1], marker=[".","x"]),
           'Experimentally adaptive in': dict(alpha=[1,1,1,0.2], facecolors=[None,None,None,None], s=[10,10,10,10], linewidths=[1,1,1,1], marker=["o","o","o","."]),
          }

# get log2effect limits
lowlim = min(sub['log2effectCCL141'].values.min(), sub['log2effectA549'].values.min())
highlim = max(sub['log2effectCCL141'].values.max(), sub['log2effectA549'].values.max())
(lowlim, highlim) = ((lowlim-(highlim-lowlim)*0.05), (highlim+(highlim-lowlim)*0.05))

for hue in hues:
    g = sns.FacetGrid(sub, height=2.5, aspect=1,
                      hue=hue, hue_order=hue_order[hue], palette=palette, hue_kws=hue_kws[hue],
                      despine=False,
                     )
    g = (g.map(plt.scatter, 'log2effectA549', 'log2effectCCL141')
         .set_axis_labels("Mutation effect in A549", "Mutation effect in CCL141")
         .set(xlim=(lowlim, highlim), ylim=(lowlim, highlim), 
             xticks=[-6, -3, 0, 3], yticks=[-6, -3, 0, 3])
        )
    plt.plot([lowlim, highlim], [lowlim, highlim], linewidth=1, color='b', alpha=0.3)
    plt.plot([0, 0], [lowlim, highlim], linewidth=1, color='b', alpha=0.3)
    plt.plot([lowlim, highlim], [0, 0], linewidth=1, color='b', alpha=0.3)
    plt.savefig('results/diffsel/scatter_muteffects_{0}.pdf'.format(hue), dpi=300, bbox_inches='tight')
    for cell in ['A549', 'CCL141']:
        g = sns.FacetGrid(sub, height=2.5, aspect=1,
                          hue=hue, hue_order=hue_order[hue], palette=palette, hue_kws=hue_kws[hue],
                          despine=False,
                         )
        g = (g.map(plt.scatter, 'log2effect{0}'.format(cell), 'mutdiffsel')
             .set_axis_labels('Mutation effect in {0}'.format(cell), "Mutation diff sel")
             .set(xlim=(lowlim, highlim), 
                  xticks=[-6, -3, 0, 3])
            )
        plt.plot([0, 0], [lowlim, highlim], linewidth=1, color='b', alpha=0.3)
        plt.plot([lowlim, highlim], [0, 0], linewidth=1, color='b', alpha=0.3)
        plt.savefig('results/diffsel/scatter_diffsel_muteffects{0}_{1}.pdf'.format(cell, hue), dpi=300, bbox_inches='tight')
```

##### Visualize top adaptive sites¶

I will plot preferences for the top adaptive sites, so as to see how reproducible they are across replicates. I will then pick the most reproducible ones to validate.

In [24]:

```
def unique(sequence):
    seen = set()
    return [x for x in sequence if not (x in seen or seen.add(x))]

# dfmut.sort_values('site')

selectedsitelist = {}
selectedsitelist['A549'] = unique(list(
        dfmut[dfmut['Experimentally adaptive in']=='A549']
        .sort_values('mutdiffsel', ascending=False)
        ['site']))
selectedsitelist['CCL141'] = unique(list(
        dfmut[dfmut['Experimentally adaptive in']=='CCL141']
        .sort_values('mutdiffsel', ascending=True)
        ['site']))
selectedsitelist['Both'] = unique(list(
        dfmut[dfmut['Experimentally adaptive in']=='Both']
        .sort_values('log2effectA549', ascending=False)
        ['site']))
print(selectedsitelist)
```

```
{'A549': [521, 163, 669, 69, 183, 701, 627, 182, 698, 292, 684, 190, 676, 158, 156, 522, 355, 176, 532, 169, 82, 9], 'CCL141': [360, 382, 377, 378, 380, 112, 342, 202, 472, 60, 344], 'Both': [60, 293, 81, 199, 82, 195, 74]}
```

In [25]:

```
libs = ['4', '5', '6']

for cell in ['A549', 'CCL141', 'Both']:
    selectedsites = [str(x) for x in selectedsitelist[cell]]
    print('Selected in {0}:'.format(cell))

    # plot prefs
    prefsA549 = [os.path.join(prefsmethoddir['Bayes2errA549']['A549'], '{0}-Lib{1}_prefs_rescaled.csv'.format('A549', i)) for i in libs]
    prefsCCL141 = [os.path.join(prefsmethoddir['Bayes2errA549']['CCL141'], '{0}-Lib{1}_prefs_rescaled.csv'.format('CCL141', i)) for i in libs]
    selectedprefs = (
            pd.concat(
                [pd.read_csv(f, dtype={'site': str}).assign(cell=cells[i], replicate=j + 1) for 
                (i, iprefs) in enumerate([prefsA549, prefsCCL141]) for (j, f) in enumerate(iprefs)])
            .query('site in @selectedsites')
            .assign(stacklabel=lambda x: x['replicate'],
                    facetlabel=lambda x: x['cell'] + '\nsite ' + x['site'].astype(str),
                    cell=lambda x: pd.Categorical(x['cell'], cells),
                    site=lambda x: pd.Categorical(x['site'], selectedsites))
            .sort_values(['site', 'cell'], ascending=[True, True])
            [['facetlabel', 'stacklabel'] + dms_tools2.AAS]
            )
    selectedprefsplot = os.path.join(diffseldir, 'selectedlogo_{0}_rescaledprefs.pdf'.format(cell))
    dms_tools2.rplot.facetedGGSeqLogo(selectedprefs, dms_tools2.AAS, selectedprefsplot, 
                                      width=2.2, height=1.8 * len(selectedsites), 
                                      ncol=2, xlabelsrotate=False)

    #plot diffsel
    selecteddiffsel = (
        pd.read_csv(os.path.join(diffseldir, 'summary_A549vCCL141-meanmutdiffsel.csv'), dtype={'site': str})
        .pivot(index='site', columns='mutation', values='mutdiffsel')
        .reset_index()
        .query('site in @selectedsites')
        [['site'] + dms_tools2.AAS]
        .assign(stacklabel='',
                facetlabel=lambda x: 'dsel\n' + x['site'].astype(str),
                site=lambda x: pd.Categorical(x['site'], selectedsites))
        .sort_values('site')
        [['facetlabel', 'stacklabel'] + dms_tools2.AAS]
        )
    selecteddiffselplot = os.path.join(diffseldir, 'selectedlogo_{0}_diffsel.pdf'.format(cell))
    dms_tools2.rplot.facetedGGSeqLogo(selecteddiffsel, dms_tools2.AAS, selecteddiffselplot, 
                                      width=0.8, height=1.8 * len(selectedsites),
                                      ncol=1)
    
    dfsub = (dfmut[dfmut['Experimentally adaptive in']==cell]
             .sort_values('mutdiffsel', ascending=False)
             [['site', 'wildtype', 'mutation', 
               'log2effectA549', 'log2effectCCL141', 'mutdiffsel',
               'Known human adaptive', 'Experimentally adaptive in']]
            )
    display(HTML(dfsub.to_html(index=False)))
    showPDF([selectedprefsplot, selecteddiffselplot], width=200)
```

```
Selected in A549:
```

| site | wildtype | mutation | log2effectA549 | log2effectCCL141 | mutdiffsel | Known human adaptive | Experimentally adaptive in |
| --- | --- | --- | --- | --- | --- | --- | --- |
| 521 | T | V | 1.139283 | -3.343339 | 3.603195 | No | A549 |
| 521 | T | L | 1.337027 | -3.179853 | 3.425011 | No | A549 |
| 521 | T | F | 2.178947 | -2.593936 | 3.350285 | No | A549 |
| 521 | T | I | 1.237696 | -2.981343 | 3.165122 | No | A549 |
| 521 | T | Y | 1.061322 | -3.340716 | 2.821094 | No | A549 |
| 163 | I | C | 1.413918 | -1.618942 | 2.577371 | No | A549 |
| 669 | G | S | 1.101390 | -1.656856 | 2.566507 | No | A549 |
| 69 | E | K | 1.119720 | -0.105373 | 2.545517 | No | A549 |
| 521 | T | W | 1.304727 | -2.729416 | 2.420922 | No | A549 |
| 183 | L | D | 1.948999 | -1.053630 | 2.297568 | No | A549 |
| 701 | D | N | 1.193828 | -1.523811 | 2.240536 | Yes | A549 |
| 627 | E | C | 1.564604 | -0.639235 | 2.177474 | No | A549 |
| 182 | Q | N | 1.267788 | -1.587876 | 2.166410 | No | A549 |
| 69 | E | R | 1.147984 | -1.006468 | 2.142959 | No | A549 |
| 701 | D | M | 1.611496 | -1.575917 | 2.039116 | No | A549 |
| 698 | G | K | 2.225857 | 0.160912 | 2.038426 | No | A549 |
| 292 | I | T | 1.958468 | -0.678898 | 2.014002 | No | A549 |
| 684 | A | K | 1.535872 | -0.818438 | 1.995105 | No | A549 |
| 190 | K | Y | 1.759519 | -0.353815 | 1.990902 | No | A549 |
| 676 | T | Y | 1.674848 | -0.501244 | 1.944572 | No | A549 |
| 158 | E | L | 1.843315 | -0.549091 | 1.912780 | No | A549 |
| 156 | A | M | 2.030228 | -0.406435 | 1.874789 | No | A549 |
| 183 | L | N | 1.138063 | -1.162777 | 1.853388 | No | A549 |
| 701 | D | C | 1.044505 | -1.220379 | 1.796576 | No | A549 |
| 522 | Q | I | 1.014306 | -1.037564 | 1.790193 | No | A549 |
| 355 | R | G | 1.186167 | -0.774579 | 1.742070 | No | A549 |
| 521 | T | Q | 1.740830 | -0.441232 | 1.730401 | No | A549 |
| 627 | E | S | 1.380895 | -0.448657 | 1.708193 | No | A549 |
| 176 | I | E | 1.020005 | -0.957745 | 1.704337 | No | A549 |
| 532 | S | D | 1.204100 | -0.865743 | 1.699329 | No | A549 |
| 182 | Q | Y | 1.199057 | -1.318473 | 1.601512 | No | A549 |
| 169 | P | E | 1.235869 | -0.588357 | 1.589386 | No | A549 |
| 82 | N | W | 1.364159 | -0.464585 | 1.558646 | No | A549 |
| 9 | D | K | 1.293712 | -0.397477 | 1.539068 | No | A549 |

```
Selected in CCL141:
```

| site | wildtype | mutation | log2effectA549 | log2effectCCL141 | mutdiffsel | Known human adaptive | Experimentally adaptive in |
| --- | --- | --- | --- | --- | --- | --- | --- |
| 360 | Y | C | -1.437267 | 1.206288 | -2.002067 | No | CCL141 |
| 344 | V | A | -1.181167 | 1.016350 | -2.082354 | No | CCL141 |
| 380 | R | E | -1.416508 | 1.214195 | -2.084150 | No | CCL141 |
| 360 | Y | N | -0.944306 | 1.656373 | -2.099354 | No | CCL141 |
| 380 | R | Q | -0.924997 | 1.640853 | -2.167365 | No | CCL141 |
| 360 | Y | I | -0.879686 | 1.382261 | -2.180377 | No | CCL141 |
| 377 | A | V | -1.178649 | 1.401660 | -2.195263 | No | CCL141 |
| 377 | A | T | -0.371594 | 1.998542 | -2.215582 | No | CCL141 |
| 377 | A | I | -1.308090 | 1.336271 | -2.217317 | No | CCL141 |
| 380 | R | S | -0.601422 | 1.657432 | -2.232334 | No | CCL141 |
| 380 | R | I | -1.349442 | 1.153572 | -2.265061 | No | CCL141 |
| 472 | E | I | -1.816822 | 1.096028 | -2.269760 | No | CCL141 |
| 380 | R | V | -1.172698 | 1.287392 | -2.276490 | No | CCL141 |
| 382 | I | E | -0.236985 | 2.161193 | -2.285283 | No | CCL141 |
| 60 | D | R | -1.075849 | 1.594977 | -2.313749 | No | CCL141 |
| 360 | Y | L | -1.125766 | 1.385466 | -2.351218 | No | CCL141 |
| 382 | I | Q | -1.154371 | 1.860766 | -2.351320 | No | CCL141 |
| 472 | E | K | -1.170984 | 1.058766 | -2.359661 | No | CCL141 |
| 380 | R | F | -1.607821 | 1.464260 | -2.362344 | No | CCL141 |
| 382 | I | H | -1.568570 | 1.338911 | -2.378713 | No | CCL141 |
| 377 | A | E | -1.008668 | 3.822810 | -2.439841 | No | CCL141 |
| 202 | M | I | -1.823291 | 2.082615 | -2.451537 | No | CCL141 |
| 380 | R | Y | -1.506141 | 1.496656 | -2.467415 | No | CCL141 |
| 377 | A | M | -1.201826 | 2.053863 | -2.490044 | No | CCL141 |
| 360 | Y | P | -1.312668 | 1.230845 | -2.553633 | No | CCL141 |
| 342 | E | W | -2.461563 | 1.178997 | -2.563885 | No | CCL141 |
| 377 | A | K | -1.748946 | 2.142287 | -2.577349 | No | CCL141 |
| 377 | A | R | -1.386154 | 1.470913 | -2.587443 | No | CCL141 |
| 112 | P | W | -1.544874 | 1.796545 | -2.590450 | No | CCL141 |
| 382 | I | Y | -2.168926 | 1.446427 | -2.593108 | No | CCL141 |
| 380 | R | T | -0.509821 | 2.413105 | -2.729927 | No | CCL141 |
| 360 | Y | T | -1.895285 | 1.102965 | -2.804739 | No | CCL141 |
| 380 | R | L | -1.863148 | 1.556991 | -2.936576 | No | CCL141 |
| 378 | T | R | -2.162609 | 1.373550 | -3.011598 | No | CCL141 |
| 360 | Y | D | -1.229828 | 2.433986 | -3.134726 | No | CCL141 |
| 382 | I | D | -1.455739 | 2.399931 | -3.138905 | No | CCL141 |
| 360 | Y | Q | -2.472410 | 1.464094 | -3.145067 | No | CCL141 |
| 360 | Y | K | -2.665158 | 2.007907 | -3.178295 | No | CCL141 |
| 377 | A | Q | -1.180051 | 2.576383 | -3.208844 | No | CCL141 |
| 377 | A | H | -1.278295 | 2.535692 | -3.264574 | No | CCL141 |
| 382 | I | R | -2.712068 | 1.347478 | -3.503473 | No | CCL141 |
| 360 | Y | R | -1.982503 | 2.092585 | -3.913665 | No | CCL141 |

```
Selected in Both:
```

| site | wildtype | mutation | log2effectA549 | log2effectCCL141 | mutdiffsel | Known human adaptive | Experimentally adaptive in |
| --- | --- | --- | --- | --- | --- | --- | --- |
| 60 | D | M | 1.802882 | 1.169637 | 0.786476 | No | Both |
| 293 | R | C | 1.759293 | 1.001078 | 0.398511 | No | Both |
| 81 | T | W | 1.758102 | 1.180048 | 0.393639 | No | Both |
| 199 | A | S | 1.734309 | 1.654414 | 0.187155 | Yes | Both |
| 82 | N | K | 1.516544 | 1.371821 | 0.097212 | No | Both |
| 195 | D | K | 1.399084 | 1.263876 | 0.060194 | No | Both |
| 74 | G | C | 1.158124 | 1.190242 | -0.245348 | No | Both |

I selected the following final set of sites for validation:

In [26]:

```
selectedsitesfinal = ([9, 82, 163, 176, 182, 183, 292, 355, 521, 522, 532] + #A549
                      [534] + #H7N9
                      [627, 669, 701] + #A549
                      [112, 378, 382, ] #CCL141
                     )
selectedsitesfinalboth = [82, 195, 293] #Both
selectedsitesfinalall = unique(selectedsitesfinal + selectedsitesfinalboth)
```

In [27]:

```
# Get wt residues to label logoplots
df = dfmut[['site', 'wildtype']].drop_duplicates()
wtresidue = dict(zip(df['site'].astype(str), df['wildtype']))
# Specify colors for characters in logoplots
char_colors = {}
chars = ['*', 'A', 'C', 'D', 'E', 'F', 'G', 'H', 'I', 'K', 'L', 'M', 'N', 'P', 'Q', 'R', 'S', 'T', 'V', 'W', 'Y']
for char in chars:
    char_colors[char] = 'gray'
```

In [28]:

```
selectedsites = [str(x) for x in selectedsitesfinalall]

prefsA549 = [os.path.join(prefsmethoddir['Bayes2errA549']['A549'], 'summary_avgprefs_rescaled.csv')]
prefsCCL141 = [os.path.join(prefsmethoddir['Bayes2errA549']['CCL141'], 'summary_avgprefs_rescaled.csv')]

selectedprefs = (
        pd.concat(
            [pd.read_csv(f, dtype={'site': str}).assign(cell=cells[i], replicate=j + 1) for 
            (i, iprefs) in enumerate([prefsA549, prefsCCL141]) for (j, f) in enumerate(iprefs)])
        .query('site in @selectedsites')
        .assign(stacklabel='',
                facetlabel=lambda x: x['site'].astype(str) + '\n' + x['cell'],
                cell=lambda x: pd.Categorical(x['cell'], cells),
                site=lambda x: pd.Categorical(x['site'], selectedsites))
        .sort_values(['cell', 'site'])
        [['facetlabel', 'stacklabel'] + dms_tools2.AAS]
        )
selectedprefsplot = os.path.join(diffseldir, 'selectedlogofinal_rescaledprefs.pdf')
dms_tools2.rplot.facetedGGSeqLogo(selectedprefs, dms_tools2.AAS, selectedprefsplot, 
                                  width=0.8 * len(selectedsites), height=5, 
                                  ncol=len(selectedsites), char_colors=char_colors)

#plot diffsel
selecteddiffsel = (
    pd.read_csv(os.path.join(diffseldir, 'summary_A549vCCL141-meanmutdiffsel.csv'), dtype={'site': str})
    .pivot(index='site', columns='mutation', values='mutdiffsel')
    .reset_index()
    .query('site in @selectedsites')
    [['site'] + dms_tools2.AAS]
    .assign(stacklabel='',
            facetlabel=lambda x: x['site'].astype(str), #'dsel\n' + 
            site=lambda x: pd.Categorical(x['site'], selectedsites))
    .sort_values('site')
    [['facetlabel', 'stacklabel'] + dms_tools2.AAS]
    )
selecteddiffsel['facetlabel'] = selecteddiffsel.apply(
    lambda row: row['facetlabel'] + '\n' + wtresidue[row['facetlabel']], axis=1)
selecteddiffselplot = os.path.join(diffseldir, 'selectedlogofinal_diffsel.pdf')
dms_tools2.rplot.facetedGGSeqLogo(selecteddiffsel, dms_tools2.AAS, selecteddiffselplot, 
                                  width=0.8* len(selectedsites), height=6,
                                  ncol=len(selectedsites), char_colors=char_colors)

showPDF([selectedprefsplot], width=1000)
showPDF([selecteddiffselplot], width=1000)
```

#### Copy files to paper figures directory¶

In [29]:

```
myfiguresdir = os.path.join(figuresdir, 'Fig3/')
if not os.path.isdir(myfiguresdir):
    os.mkdir(myfiguresdir)

files = (['results/diffsel/logoDiffselmean_diffsel.pdf', 
          'results/diffsel/knownadapt_mutdiffsel_boxplot.pdf',
          'results/diffsel/knownadapt_mutdiffsel_rankplot.pdf',
          'results/diffsel/summary_prefs_effects_diffsel.csv'] +
         ['results/diffsel/scatter_muteffects_{0}.pdf'.format(hue) 
          for hue in hues] +
         ['results/diffsel/scatter_diffsel_muteffects{0}_{1}.pdf'.format(cell, hue) 
          for hue in hues for cell in cells] +
         ['results/diffsel/selectedlogofinal_rescaledprefs.pdf',
          'results/diffsel/selectedlogofinal_diffsel.pdf']
        )
for f in files:
    shutil.copy(f, myfiguresdir)
```
