## Supplementary material for "Comprehensive mapping of avian influenza polymerase adaptation to the human host": File S2: Fig4_Validation.html


### Analyze and plot results from minigenome and competition validation assays¶

##### Import modules, define directories¶

In [1]:

```
import os
import glob
import shutil
import pandas as pd
import numpy as np
import matplotlib as mpl
import matplotlib.pyplot as plt
%matplotlib inline

import seaborn as sns

from IPython.display import display, HTML, Markdown, Image

fastqdir = './fastq/'
paperdir = './paper'
figuresdir = os.path.join(paperdir, 'figures/')

for xdir in [fastqdir, paperdir, figuresdir]:
    if not os.path.isdir(xdir):
        os.mkdir(xdir)
```

In [2]:

```
## Set plotting parameters/preferences
# Set colors to use throughout
palette ={"None":"gray",
          "Human":"#d95f02", "A549":"#d95f02", "Known - Human":"#d95f02", "H7N9 - Human":"#d95f02",
          "Bird":"#1b9e77", "Avian":"#1b9e77", "CCL141":"#1b9e77", 
          "Both":"#7570b3", "Known - Both":"#7570b3",
          "Yes": "#d95f02", "No": "gray" #KnownAdaptive
         }
# Set matplotlib rcParams for figures
sns.set(context='paper', style='ticks', palette='deep', font='Arial', font_scale=1.042, color_codes=True,
        rc = {'font.size': 10,
 'axes.labelsize': 10,
 'axes.titlesize': 10,
 'xtick.labelsize': 8, #9,
 'ytick.labelsize': 8, #9,
 'legend.fontsize': 9,
 'axes.linewidth': 1.0,
 'grid.linewidth': 0.8,
 'lines.linewidth': 1,
 'lines.markersize': 4.5,
 'patch.linewidth': 0.8,
 'xtick.major.width': 1.0,
 'ytick.major.width': 1.0,
 'xtick.minor.width': 0.8,
 'ytick.minor.width': 0.8,
 'xtick.major.size': 4.5,
 'ytick.major.size': 4.5,
 'xtick.minor.size': 3,
 'ytick.minor.size': 3}
       )
# sns.plotting_context()
```

#### Minigenome assays¶

In [3]:

```
## Annotated raw flow data, write annotated data to file.
## Do not need to run again. Code block here for record only.

hostcells = ['A549', '293T']
welltoplasmidfile = 'validation/minigenome/Annot_WelltoPlasmid.txt'
plasmidtomutfile = 'validation/minigenome/Annot_PlasmidtoMutatation.txt'

renamecol = dict(zip(['Sample:', 'PE-mCherry-A+ | Freq. of Parent', 'PE-mCherry-A+/GFP-A+ | Freq. of Parent'], 
                     ['Sample', '% mCherry+', '% GFP+|mCherry+']))

for cell in hostcells:
    flowfile = 'validation/minigenome/{0}_flow.txt'.format(cell)
    flowannotfile = 'validation/minigenome/{0}_flow_withannot.txt'.format(cell)

    df = (pd.read_table(flowfile)
          .rename(columns=lambda x: x.replace('Cells/', ''))
          .rename(columns=renamecol)
          .assign(Well=lambda x: x['Sample'].str.split('_').str[2])
         )

    WellToPlasmid = (pd.read_table(welltoplasmidfile)
                     .assign(Plasmid=lambda x:x['Sample ID'].str.split(' ').str[0]) )
    PlasmidToMutation = pd.read_table(plasmidtomutfile, dtype={'Plasmid':object})
    df = pd.merge(df, WellToPlasmid, how='left', on='Well')
    df = pd.merge(df, PlasmidToMutation, how='left', on='Plasmid')

    df = df[['Site', 'Mutation', 'Selected in', 'Plasmid',
           'Plasmid name', 'Sample', '% mCherry+',
           '% GFP+|mCherry+', 'Well', 'Sample ID']].sort_values('Site')
    print('Write annotated flow data to {0}'.format(flowannotfile))
    df.to_csv(flowannotfile.format(cell), index=False)
    display(HTML(df.head().to_html(index=False)))
```

```
Write annotated flow data to validation/minigenome/A549_flow_withannot.txt
```

| Site | Mutation | Selected in | Plasmid | Plasmid name | Sample | % mCherry+ | % GFP+\|mCherry+ | Well | Sample ID |
| --- | --- | --- | --- | --- | --- | --- | --- | --- | --- |
| 0 | WT | None | 2150 | 2150\_HDM-S009-PB2 | Specimen\_001\_A1\_A01\_001.fcs | 22.9 | 1.75 | A1 | 2150 1 |
| 0 | WT | None | 2150 | 2150\_HDM-S009-PB2 | Specimen\_001\_A2\_A02\_002.fcs | 27.4 | 2.74 | A2 | 2150 3 |
| 0 | WT | None | 2150 | 2150\_HDM-S009-PB2 | Specimen\_001\_A3\_A03\_003.fcs | 24.3 | 3.31 | A3 | 2150 4 |
| 9 | D9K | A549 | 2191 | 2191\_HDM-S009-PB2-D9K | Specimen\_001\_F2\_F02\_062.fcs | 22.5 | 4.57 | F2 | 2191 3 |
| 9 | D9K | A549 | 2191 | 2191\_HDM-S009-PB2-D9K | Specimen\_001\_F1\_F01\_061.fcs | 24.0 | 4.33 | F1 | 2191 2 |

```
Write annotated flow data to validation/minigenome/293T_flow_withannot.txt
```

| Site | Mutation | Selected in | Plasmid | Plasmid name | Sample | % mCherry+ | % GFP+\|mCherry+ | Well | Sample ID |
| --- | --- | --- | --- | --- | --- | --- | --- | --- | --- |
| 0 | WT | None | 2150 | 2150\_HDM-S009-PB2 | Specimen\_001\_A1\_A01\_001.fcs | 36.4 | 36.3 | A1 | 2150 1 |
| 0 | WT | None | 2150 | 2150\_HDM-S009-PB2 | Specimen\_001\_A2\_A02\_002.fcs | 39.0 | 36.7 | A2 | 2150 3 |
| 0 | WT | None | 2150 | 2150\_HDM-S009-PB2 | Specimen\_001\_A3\_A03\_003.fcs | 40.9 | 38.8 | A3 | 2150 4 |
| 9 | D9K | A549 | 2191 | 2191\_HDM-S009-PB2-D9K | Specimen\_001\_F2\_F02\_062.fcs | 45.8 | 49.2 | F2 | 2191 3 |
| 9 | D9K | A549 | 2191 | 2191\_HDM-S009-PB2-D9K | Specimen\_001\_F1\_F01\_061.fcs | 42.8 | 47.5 | F1 | 2191 2 |

In [4]:

```
hostcells = ['A549', '293T']
xaxis = 'Mutation'
ymetric = '% GFP+|mCherry+'
categories = [('None', ''), 
              ('Known - Human', 'Known\nhuman\nadaptive'), 
              ('A549', 'Top adaptive in A549'), 
              ('H7N9 - Human', 'Adaptive\nin A549,\nobserved\nin H7N9'), 
              ('CCL141', 'Top\nadaptive\nin\nCCL141'), 
              ('Both', 'Top\nadaptive\nin\nboth')]
ylim = {'A549': [0.1, 100],
       '293T': [10, 100]}

for cell in hostcells:
    flowannotfile = 'validation/minigenome/{0}_flow_withannot.txt'.format(cell)
    df = pd.read_csv(flowannotfile)
    ylabel = 'Minigenome\nActivity\nin {0}:\n% GFP+'.format(cell)

    WTmean = df[df['Mutation']=='WT']['% GFP+|mCherry+'].mean()

    fig, axes = plt.subplots(1,6, 
                             figsize=(17*0.5,2.5),
                             gridspec_kw = {'width_ratios':[1, 1, 11, 2, 2, 1]})
    plt.subplots_adjust(wspace=0.05)
    for i, (cat, catlab) in enumerate(categories):
        ax = axes.flat[i]
        dftemp = df[df['Selected in']==cat]
        c = palette[cat]
        ax.set_yscale("log", basey=10)
        ax.set_ylim(ylim[cell])

        if cat=='Known - Human': #i==1:
            g = sns.swarmplot(x=xaxis, y=ymetric, data=dftemp, 
                          ax=ax, s=6, color=c, marker='X',
                     )
        elif cat=='H7N9 - Human': #i==3:
            g = sns.swarmplot(x=xaxis, y=ymetric, data=dftemp, 
                          ax=ax, s=4, color=c, marker='s',
                     )
        else:
            g = sns.swarmplot(x=xaxis, y=ymetric, data=dftemp, 
                          ax=ax, s=4, color=c,
                     )
        sns.despine()
        ax.set_xlabel('')
        ax.set_xticklabels(ax.get_xticklabels(), rotation=90, ha='center')
        ax.set_title(catlab, fontsize=10)
        ax.axhline(WTmean, color='gray', linewidth=1)
        ax.set_yticklabels([], minor=True)
        if i==0:
            ax.set_ylabel(ylabel, 
                         rotation='horizontal', va='center', labelpad=30)
        else:
            ax.set_ylabel('')
            ax.set_yticklabels([])
    fig.savefig('validation/minigenome/minigenome_{0}.pdf'.format(cell), bbox_inches="tight", dpi=300)
```

#### Competition assays¶

##### Read in sample sheet and download data¶

In [5]:

```
comptfolder = 'validation/competition'

samples = (pd.read_table('validation/competition/SraRunTable_compt.txt', sep='\t')
           [['Sample_Name', 'lab_host', 'Timepoint', 'Mutant', 'replicate', 'Run', 'Library_Name', ]]
           .sort_values('Library_Name')
           .rename(columns={'replicate':'BioRep'})
          )

samples['Cell'] = samples.apply(lambda x:x['lab_host'].split(sep=' ')[0],axis=1)
samples['file'] = samples.apply(lambda x:x['Library_Name']+'.fastq.gz',axis=1) ## TEMP added *

display(HTML(samples.head().to_html(index=False)))
```

| Sample\_Name | lab\_host | Timepoint | Mutant | BioRep | Run | Library\_Name | Cell | file |
| --- | --- | --- | --- | --- | --- | --- | --- | --- |
| compt\_R355G\_1\_A549\_10hpi | A549 cells | 10 | R355G | 1 | SRR8375145 | A-10-10 | A549 | A-10-10.fastq.gz |
| compt\_S532D\_1\_A549\_10hpi | A549 cells | 10 | S532D | 1 | SRR8375144 | A-10-11 | A549 | A-10-11.fastq.gz |
| compt\_G669S\_1\_A549\_10hpi | A549 cells | 10 | G669S | 1 | SRR8375051 | A-10-12 | A549 | A-10-12.fastq.gz |
| compt\_N82K\_1\_A549\_10hpi | A549 cells | 10 | N82K | 1 | SRR8375054 | A-10-13 | A549 | A-10-13.fastq.gz |
| compt\_T378R\_1\_A549\_10hpi | A549 cells | 10 | T378R | 1 | SRR8375053 | A-10-14 | A549 | A-10-14.fastq.gz |

In [6]:

```
use_existing = True # Whether to use existing files

downloadfiles = dict(zip(list(samples['Run']), list(samples['file'])))
print('Downloading fastq files', end=' ')
for run, file in downloadfiles.items():
    filenameold = os.path.join(fastqdir, '{0}.fastq.gz'.format(run))
    filenamenew = os.path.join(fastqdir, file)
    if os.path.isfile(filenamenew) and use_existing:
        print('x', end='')
    else:
        print('.', end='')
        log = !fastq-dump --outdir $fastqdir --gzip $run
        log = !mv $filenameold $filenamenew
print('Done')
```

```
Downloading fastq files xxxxxxxxxxxxxxxxxxxxxxxxxxxxxxxxxxxxxxxxxxxxxxxxxxxxxxxxxxxxxxxxxxxxxxxxxxxxxxxxxxxxxxxxxxxxxxxxxxxxxxxxxxxxxxxxxxxxxxxxxxxxxxxxxxxxxxxxDone
```

##### Count number of reads mapping to WT/mutant sequence in fastq files¶

In [7]:

```
# read in reference sequences to match to
RefSeq_df = pd.read_table(os.path.join(comptfolder, 'MutantRefSeq.txt') )\
    .set_index('Mutant')
RefSeq = RefSeq_df.to_dict()

# Count reads mapping to WT/mutant sequences, save to dataframe/file
SampleSheet = samples[['Sample_Name', 'Library_Name', 'Cell', 'Timepoint', 'BioRep', 'Mutant', 'file']]
countsdf = SampleSheet.reindex( columns = SampleSheet.columns.tolist() + ['TotalReads','WtReads','MutReads'])
for i, row in SampleSheet.iterrows():
    fastq = os.path.join(fastqdir, row['file'])
    seq_wt = RefSeq['seq_wt'][row['Mutant']]
    seq_mut = RefSeq['seq_mut'][row['Mutant']]
    TotalReads = !zcat $fastq | wc -l
    WtReads = !zgrep -c $seq_wt $fastq
    MutReads = !zgrep -c $seq_mut $fastq
    countsdf.loc[i,'TotalReads'] = int(TotalReads[0])/4
    countsdf.loc[i,'WtReads'] = int(WtReads[0])
    countsdf.loc[i,'MutReads'] = int(MutReads[0])
countsdf.to_csv(path_or_buf='validation/competition/competition_counts.txt', sep='\t', index=False)
```

In [8]:

```
countsdf.head()
```

Out[8]:

|  | Sample\_Name | Library\_Name | Cell | Timepoint | BioRep | Mutant | file | TotalReads | WtReads | MutReads |
| --- | --- | --- | --- | --- | --- | --- | --- | --- | --- | --- |
| 89 | compt\_R355G\_1\_A549\_10hpi | A-10-10 | A549 | 10 | 1 | R355G | A-10-10.fastq.gz | 378336.0 | 225854.0 | 108892.0 |
| 90 | compt\_S532D\_1\_A549\_10hpi | A-10-11 | A549 | 10 | 1 | S532D | A-10-11.fastq.gz | 252360.0 | 160831.0 | 21663.0 |
| 7 | compt\_G669S\_1\_A549\_10hpi | A-10-12 | A549 | 10 | 1 | G669S | A-10-12.fastq.gz | 466655.0 | 325608.0 | 101072.0 |
| 4 | compt\_N82K\_1\_A549\_10hpi | A-10-13 | A549 | 10 | 1 | N82K | A-10-13.fastq.gz | 338068.0 | 209271.0 | 96922.0 |
| 5 | compt\_T378R\_1\_A549\_10hpi | A-10-14 | A549 | 10 | 1 | T378R | A-10-14.fastq.gz | 457516.0 | 381797.0 | 21834.0 |

##### Calculate enrichment and plot¶

I calculated enrichment in A549 over CCL141 cells for each mutant at each timepoint as follows:

$$
\begin{align}
Enrichment = \frac{\left(count\_{mut}^{A549}\right) /
\left(count\_{wt}^{A549}\right)}
{\left(count\_{mut}^{CCL141}\right) /
\left(count\_{wt}^{CCL141}\right)} \\
\end{align}
$$

In [9]:

```
# First get countmut/countwt for each respective cell
countsdf = countsdf[['Cell', 'Timepoint', 'BioRep', 'Mutant', 'WtReads', 'MutReads']].copy()
countsdf['Mut/WT'] = countsdf['MutReads']/countsdf['WtReads']
#Then calculate this enrichment ratio comparing cell types 
countsdf_A549 = countsdf[countsdf['Cell']=='A549']
countsdf_CCL141 = countsdf[countsdf['Cell']=='CCL141']
countsdf2 = pd.merge(countsdf_A549, countsdf_CCL141, how='inner', 
                     on=['Timepoint', 'BioRep', 'Mutant'], 
                     suffixes=['_A', '_C'])
countsdf2['EnrichmentInA549'] = countsdf2['Mut/WT_A']/countsdf2['Mut/WT_C']
countsdf2 = countsdf2[['Timepoint', 'BioRep', 'Mutant', 'EnrichmentInA549']]
countsdf2.head()
```

Out[9]:

|  | Timepoint | BioRep | Mutant | EnrichmentInA549 |
| --- | --- | --- | --- | --- |
| 0 | 10 | 1 | R355G | 1.421011 |
| 1 | 10 | 1 | S532D | 0.370057 |
| 2 | 10 | 1 | G669S | 0.522676 |
| 3 | 10 | 1 | N82K | 0.754000 |
| 4 | 10 | 1 | T378R | 0.133511 |

In [10]:

```
# Make dictionary of where mutations are selected in, for plotting downstream
AnnotateMut = pd.read_table('validation/competition/Annot_Mutation.txt')[['Mutation', 'Selected in']]
AnnotateMutDict = dict(zip(AnnotateMut['Mutation'], AnnotateMut['Selected in']))
```

In [11]:

```
## Dataframes for plotting
# Make a dataframe with NaN at timepoint 0 (for plotting actual data)
dffinal = countsdf2
dfaddzero = dffinal[dffinal['Timepoint']==10].copy()
dfaddzero['Timepoint'] = 0
dfaddzero['EnrichmentInA549']=np.NaN
dffinal = pd.concat([dfaddzero, dffinal])
# Make a dataframe with assumed enrichment (=1) for timepoint 0, 
# and NaN at timepoint 48 (for plotting assumed start at enrichment ratio of 1)
dffinal2 = dffinal.copy()
dffinal2.loc[dffinal2['Timepoint']==0, 'EnrichmentInA549']=1
dffinal2.loc[dffinal2['Timepoint']==48, 'EnrichmentInA549']=np.NaN
```

In [12]:

```
# Set ymin/max based on spread of data
ymetric = 'EnrichmentInA549'
print(dffinal[ymetric].min(), dffinal[ymetric].max())
ymin, ymax = 0.01, 250
# Order to plot
ordered = ['E627E',
           'E627K',
           'D9K', 'N82W', 'I163C', 'I292T', 'R355G', 'T521F', 'S532D', 'E627C', 'G669S', 'D701M', 
           'R355K', 'S534F',
           'T378R', 'E627D',
           'N82K',]
print(len(ordered))
```

```
0.012585635282365894 190.31403504324564
17
```

In [13]:

```
fig, axes = plt.subplots(1, len(ordered), figsize=(len(ordered)*0.5,2.5))
for i, mut in enumerate(ordered):
    ax = axes.flat[i]
    c = palette[AnnotateMutDict[mut]]
    dftempA = dffinal[dffinal['Mutant']==mut]
    dftempB = dffinal2[dffinal2['Mutant']==mut]
    dftempA1 = dftempA[dftempA['BioRep']==1]
    dftempA2 = dftempA[dftempA['BioRep']==2]
    dftempB1 = dftempB[dftempB['BioRep']==1]
    dftempB2 = dftempB[dftempB['BioRep']==2]
    ax.set_yscale("log", basey=10)
    ax.set_ylim([ymin, ymax])
    
    if i in [1]: #syn control
        g = sns.pointplot(x='Timepoint', y=ymetric, data=dftempA1, ax=ax, color=c, markers='x')
        g = sns.pointplot(x='Timepoint', y=ymetric, data=dftempA2, ax=ax, color=c, markers='x')
        g = sns.pointplot(x='Timepoint', y=ymetric, data=dftempB1, ax=ax, color=c, markers='x', linestyles=["--"])
        g = sns.pointplot(x='Timepoint', y=ymetric, data=dftempB2, ax=ax, color=c, markers='x', linestyles=["--"])
    elif i in [12, 13]: #H7N9
        g = sns.pointplot(x='Timepoint', y=ymetric, data=dftempA1, ax=ax, color=c, markers='s')
        g = sns.pointplot(x='Timepoint', y=ymetric, data=dftempA2, ax=ax, color=c, markers='s')
        g = sns.pointplot(x='Timepoint', y=ymetric, data=dftempB1, ax=ax, color=c, markers='s', linestyles=["--"])
        g = sns.pointplot(x='Timepoint', y=ymetric, data=dftempB2, ax=ax, color=c, markers='s', linestyles=["--"])
    else:
        g = sns.pointplot(x='Timepoint', y=ymetric, data=dftempA1, ax=ax, color=c)
        g = sns.pointplot(x='Timepoint', y=ymetric, data=dftempA2, ax=ax, color=c)
        g = sns.pointplot(x='Timepoint', y=ymetric, data=dftempB1, ax=ax, color=c, linestyles=["--"])
        g = sns.pointplot(x='Timepoint', y=ymetric, data=dftempB2, ax=ax, color=c, linestyles=["--"])    
    
    plt.setp(ax.lines, linewidth=1)
    plt.setp(ax.collections, sizes=[10])
    g.set_title(mut, fontsize=8)
    ax.axhline(1, color='gray', linewidth=1)
    ax.set_xticklabels(ax.get_xticklabels(), rotation=90, ha='left')
    if i==0:
        sns.despine(ax=ax)
        g.set_ylabel('Viral\ncompetition:\nEnrichment\nin A549 over\nCCL141', 
                     rotation='horizontal', va='center', labelpad=30)
    elif i in [0, 1, 2, 12, 14, 16]:
        sns.despine(ax=ax)
        g.set_ylabel('')
        g.set_yticklabels([])
    else:
        sns.despine(ax=ax, left=True)
        g.set_ylabel('')
        g.set_yticklabels([])
        g.set_yticks([], minor=True)
        g.set_yticks([], minor=False)
    if i == int(len(ordered)/2):
        g.set_xlabel('Hours post-infection')
    else:
        g.set_xlabel('')
        
fig.savefig('validation/competition/competition.pdf', bbox_inches="tight", dpi=300)
```

#### Copy files to paper figures directory¶

In [14]:

```
paperdir = './paper'
figuresdir = os.path.join(paperdir, 'figures/')
myfiguresdir = os.path.join(figuresdir, 'Fig4/')
if not os.path.isdir(myfiguresdir):
    os.mkdir(myfiguresdir)

files = (['validation/minigenome/{0}_flow_withannot.txt'.format(cell) 
          for cell in hostcells] +
         ['validation/minigenome/minigenome_{0}.pdf'.format(cell) 
          for cell in hostcells] +
         ['validation/competition/competition_counts.txt',
          'validation/competition/competition.pdf']
        )
for f in files:
    shutil.copy(f, myfiguresdir)
```
