## Supplementary material for "Comprehensive mapping of avian influenza polymerase adaptation to the human host": File S2: Fig5_MapDiffSelToStructure.html


### Map differential selection to structures¶

We generated pymol scripts to display positive differential selection values on various PB2 structures.

| Structure | pdb name | Input txt files | Output pymol scripts |
| --- | --- | --- | --- |
| Polymerase in transcription pre-initiation form | 4wsb | preamble   setview   colorfullstructure   topA549 | map\_fullstructure.py   map\_topA549.py   map\_positive\_diff\_seq.py |
| Polymerase in apo form | 59d8 | preamble   setview   colorfullstructure   topA549 | map\_fullstructure.py   map\_topA549.py   map\_positive\_diff\_seq.py |
| PB2 in complex with RNA pol II | 6f5o | preamble   setview   MartinResidues   topA549 | map\_MartinResidues.py   map\_positive\_diff\_seq.py |
| PB2 in complex with importin | 4uae | preamble   setview   PumroyResidues   topA549 | map\_PumroyResidues.py   map\_positive\_diff\_seq.py |

Import modules, define directories

In [1]:

```
import os
import shutil
import pandas as pd
import numpy as np
from colour import Color
```

Generate `topA549.txt`

In [3]:

```
# Gather prior annotations for known human/mammalian adaptive mutations
mutsAdaptive = pd.read_table('data/Muts_HumanAvian.txt')
knownAdaptive = (set(mutsAdaptive[mutsAdaptive['ExptVerified']=='Yes']['Site']))
# Gather top adaptive mutations in A549
dmssummarydf = pd.read_csv('results/diffsel/summary_prefs_effects_diffsel.csv')
topA549 = (set(dmssummarydf[dmssummarydf['Experimentally adaptive in']=='A549']['site']))
# Intersect
knownAdaptiveAndTopA549 = (knownAdaptive & topA549)

# Set colors for each set of mutations
mutsets = [((knownAdaptive, 'knownAdaptive'), 'blue'), 
           ((topA549, 'topA549'), 'red'), 
           ((knownAdaptiveAndTopA549, 'knownAdaptiveAndTopA549'), 'magenta')]

# Write to file
outfile = 'pymol/topA549.txt'
f = open(outfile, 'w')
f.write('metric = \'topA549\'\n')
f.write('cmd.color(\'white\', \'S009\')\n\n')
for ((mutlist, mutlistname), col) in mutsets:
    f.write('# {0}\n'.format(mutlistname))
    f.write('for r in [{0}]:\n'.format(', '.join([str(mut) for mut in mutlist])))
    f.write('\tcmd.color(\'{0}\', \'resi {{0}} and S009\'.format(r))\n'.format(col))
    f.write('\tcmd.show(\'spheres\', \'resi {{0}} and S009\'.format(r))\n\n')
f.close()
```

Generate pymol scripts

In [4]:

```
# Standard across all structures
postamble = 'pymol/postamble.txt'
fullstructure = 'pymol/colorfullstructure.txt'
topA549 = 'pymol/topA549.txt'
pumroy = 'pymol/PumroyResidues.txt'
martin = 'pymol/MartinResidues.txt'
specialmetricfiles = {'fullstructure': fullstructure,
                     'topA549': topA549, 
                     'pumroy': pumroy,
                     'martin': martin}
# For non-specialmetrics
diffsel_file = 'results/diffsel/summary_A549vCCL141-meansitediffsel.csv'
metricdf = pd.read_csv(diffsel_file).sort_values('site')
colspectrum = 'white_red'

def write_pymol_script(structure, metric, metricdf, colspectrum):
    outfile = 'pymol/{0}_map_{1}.py'.format(structure, metric)
    preamble = 'pymol/preamble_{0}.txt'.format(structure)
    setview = 'pymol/setviewamble_{0}.txt'.format(structure)

    f = open(outfile, 'w')

    with open(preamble, 'r') as addtext:
        for line in addtext.readlines():
            f.write(line)
            if metric=='fullstructure' and line=='### To get PB2 only\n': 
                #don't print lines that will remove non-PB2 subunits
                break
        f.write('\n\n')
    
    # Write commands to color structure
    if metric in specialmetricfiles:
        with open(specialmetricfiles[metric], 'r') as addtext:
            for line in addtext.readlines():
                f.write(line)
            f.write('\n\n')
    else:
        f.write("metric = \'{0}\'\n".format(metric))
        for row in metricdf.itertuples(index=True, name='Pandas'):
            site, met = (getattr(row, "site"), getattr(row, metric))
            f.write("cmd.alter(\'resi {0}\', \'b = {1}\')\n".format(site, met))
        f.write('\n')
        f.write("cmd.spectrum(\'b\', \'{0}\', \'S009\')\n".format(colspectrum))
        f.write('\n')

    with open(setview, 'r') as addtext:
        for line in addtext.readlines():
            f.write(line)
    f.write('\n')
    with open(postamble, 'r') as addtext:
        for line in addtext.readlines():
            f.write(line)

    f.close()
```

In [9]:

```
structures = ['4wsb', '5d98']
metrics = ['fullstructure', 'topA549', 'positive_diffsel']

for structure in structures:
    for metric in metrics:
        write_pymol_script(structure, metric, metricdf, colspectrum)
```

In [10]:

```
structures = ['6f5o']
metrics = ['martin', 'positive_diffsel', 'topA549']

for structure in structures:
    for metric in metrics:
        write_pymol_script(structure, metric, metricdf, colspectrum)
```

In [12]:

```
structures = ['4uad']
metrics = ['pumroy', 'positive_diffsel', 'topA549']

for structure in structures:
    for metric in metrics:
        write_pymol_script(structure, metric, metricdf, colspectrum)
```

#### Copy files to paper figures directory¶

In [13]:

```
paperdir = './paper'
figuresdir = os.path.join(paperdir, 'figures/')
myfiguresdir = os.path.join(figuresdir, 'Fig5/')
if not os.path.isdir(myfiguresdir):
    os.mkdir(myfiguresdir)

filespy = !ls pymol/*.py
filespdb = !ls pymol/*.pdb
for f in filespy + filespdb:
    shutil.copy(f, myfiguresdir)
```
