## Supplementary material for "Comprehensive mapping of avian influenza polymerase adaptation to the human host": File S2: Fig6_H7N9.html


### Compare the effect of mutations occurring during avian-to-human tranmission of influenza, vs mutations occuring within avian influenza¶

Are mutations occurring during avian-to-human transmission of influenza are more likely to be identified as beneficial in our deep mutational scan? We will compare mutations occuring during avian-to-human transmission of H7N9 influenza, to mutations occurring within avian influenza.

###### Import modules, define directories, define functions¶

In [12]:

```
import sys, subprocess, glob, os, shutil, re, importlib, csv, json, copy, argparse
from collections import Counter
import warnings
warnings.simplefilter('ignore') # don't print warnings to avoid clutter in notebook

import numpy as np
from scipy import stats
import pandas as pd
import matplotlib as mpl
import matplotlib.pyplot as plt
mpl.use("Agg")
from matplotlib import gridspec
from matplotlib.collections import LineCollection

import Bio.Phylo
from Bio.SeqRecord import SeqRecord
from Bio import SeqIO

from IPython.display import display, HTML, Markdown, Image

# Directories
resultsdir = './results/' 
h7n9dir = os.path.join(resultsdir, 'H7N9/')
if not os.path.isdir(h7n9dir):
    os.mkdir(h7n9dir)
```

In [13]:

```
# Set plotting parameters
import seaborn as sns
sns.set(context='paper', style='ticks', palette='deep', font='Arial', font_scale=1.042, color_codes=True,
        rc = {'font.size': 10,
 'axes.labelsize': 10,
 'axes.titlesize': 10,
 'xtick.labelsize': 9,
 'ytick.labelsize': 9,
 'legend.fontsize': 9,
 'axes.linewidth': 1.0,
 'grid.linewidth': 0.8,
 'lines.linewidth': 1,
 'lines.markersize': 4.5,
 'patch.linewidth': 0.8,
 'xtick.major.width': 1.0,
 'ytick.major.width': 1.0,
 'xtick.minor.width': 0.8,
 'ytick.minor.width': 0.8,
 'xtick.major.size': 4.5,
 'ytick.major.size': 4.5,
 'xtick.minor.size': 3,
 'ytick.minor.size': 3}
       )
# sns.plotting_context()
```

#### Classification of mutations on H7N9 PB2 tree¶

We started with the annotated tree of H7N9 PB2 sequences as available on `nextstrain`. This tree was generated by `nextstrain's` `augur` pipeline, in which sequences are aligned, a phylogeny is constructed from the aligned sequences, and the phylogeny is annotated with inferred ancestral pathogen dates, sequences, and traits including mutations.

##### Functions for reading jsons and generating BioPhylo tree object¶

These functions are copied/derived from nextstrain augur/base/io\_util.py.

In [14]:

```
# function to use the json module to read in a json file and store it as "data"                
def read_json(file_name):
    try:
        handle = open(file_name, 'r')
    except IOError:
        pass
    else:
        data = json.load(handle)
        handle.close()
    return data

# code for parsing through tree jsons and returning descendents
def all_descendants(node):
    """Take node, ie. dict, and return a flattened list of all nodes descending from this node"""
    yield node
    
    # this will recursively return all internal nodes (nodes with children)
    if 'children' in node:
        for child in node['children']:
            for desc in all_descendants(child):
                yield desc
                
# Biopython's trees don't store links to node parents, so we need to build
# a map of each node to its parent.
# Code from the Bio.Phylo cookbook: http://biopython.org/wiki/Phylo_cookbook
def all_parents(tree):
    parents = {}
    for clade in tree.find_clades(order='level'):
        for child in clade:
            parents[child] = clade
    return parents

def annotate_parents(tree):
    # Get all parent nodes by node.
    parents_by_node = all_parents(tree)

    # Next, annotate each node with its parent.
    for node in tree.find_clades():
        if node == tree.root:
            node.up = None
        else:
            node.up = parents_by_node[node]

    # Return the tree.
    return tree

# Get children of a node
def all_children(tree):
    children = {}
    for clade in tree.find_clades(order='level'):
        children[clade] = [child for child in clade]
    return children

def annotate_children(tree):
    # Get all children nodes by node.
    children_by_node = all_children(tree)

    # Next, annotate each node with its parent.
    for node in tree.find_clades():
        if node.is_terminal():
            node.down = None
        else:
            node.down = children_by_node[node]

    # Return the tree.
    return tree

def json_to_tree(json_dict, root=True):
    """Returns a Bio.Phylo tree corresponding to the given JSON dictionary exported
    by `tree_to_json`.

    Assigns links back to parent nodes for the root of the tree.

    >>> import json
    >>> json_fh = open("tests/data/json_tree_to_nexus/flu_h3n2_ha_3y_tree.json", "r")
    >>> json_dict = json.load(json_fh)
    >>> tree = json_to_tree(json_dict)
    >>> tree.name
    u'NODE_0002020'
    >>> len(tree.clades)
    2
    >>> tree.clades[0].name
    u'NODE_0001489'
    >>> hasattr(tree, "attr")
    True
    >>> "dTiter" in tree.attr
    True
    """
    node = Bio.Phylo.Newick.Clade()
    node.name = json_dict["strain"]

    if "children" in json_dict:
        # Recursively add children to the current node.
        node.clades = [json_to_tree(child, root=False) for child in json_dict["children"]]

    # Assign all non-children attributes.
    for attr, value in json_dict.items():
        if attr != "children":
            setattr(node, attr, value)

    node.numdate = node.attr.get("num_date")
    node.branch_length = node.attr.get("div")

    if "translations" in node.attr:
        node.translations = node.attr["translations"]

    if root:
        node = annotate_parents(node)
        node = annotate_children(node)

    return node
```

##### Parse H7N9 PB2 tree¶

This code will loop through each clade in the tree (starting from the root out to the terminal nodes). For each clade, gather all terminal nodes that fall within that clade. If the current clade includes only human sequences, then it will be labelled human to human or bird to human as follows:

**1. Human to human:** If the current clade includes only human sequences, look at its parent node. If all terminal branches stemming from its parent node are human, then the current clade falls within a monophyletic human clade. This branch is labelled "human to human." SNPs on this branch are added to the "human\_to\_human" list.

**2. Bird to human:** If the current clade includes only human sequences, look at its parent node. If the terminal branches stemming from its parent node include both human and nonhuman sequences, then the current clade represents the branch leading to a monophyletic human clade. This branch is labelled "bird to human." SNPs on this branch are added to the "bird\_to\_human" list.

If the current clade includes a mixture of human and nonhuman sequences, the branch leading to that clade is labelled as "bird to bird". Any SNPs that occur along this branch are added to the "bird\_to\_bird" list.

**Tree tips** are treated as clades themselves, and are categorized as above. For example, a human tip that falls within a bird clade will have the branch leading to it labelled as bird to human, and those SNPs will be added to the bird\_to\_human list. A human tip that falls within a human-only clade will have its branch labelled as human to human, and its SNPs will be added to the human\_to\_human list.

Note that since H7N9 human influenza is thought to typically arise from avian\_to\_human transmissions, what we identify as human\_to\_human transmissions likely actually arise from avian\_to\_human transmissions. However, we are unable to accurately assign them as avian\_to\_human transitions because of insufficient sampling of avian sequences.

We classified as human mutations those occurring on bird\_to\_human and human\_to\_human branches. We classified as avian mutations those occurring on bird\_to\_bird branches.

In [15]:

```
# Read in PB2 json tree file and convert to BioPhylo object

# tree I want to use for this analysis 
PB2_tree = "data/flu_avian_h7n9_pb2_tree_current_2018-11-20.json"

# read in json and convert to a BioPhylo tree object
tree = read_json(PB2_tree)
tree = json_to_tree(tree)
# tree
```

In [16]:

```
# lists and dicts to store amino acid mutations 
bird_to_bird = []
bird_to_human = []
human_to_human = []
bird_to_bird_d = []
bird_to_human_d = []
human_to_human_d = []
host = {}
mutations = {}

# loop through all the nodes in the tree, going from root to tip 
for clade in tree.find_clades(): 
    tips_list = []
    hosts_list = []
    mutlist = []
    
    # for each clade, output all terminal nodes that fall within that clade 
    for terminal in clade.get_terminals():
        hosts_list.append(terminal.attr['host'])
        tips_list.append(terminal.name)
        
    # if all terminal nodes are human, print the clade name and the list of terminal branches it entails
    if set(hosts_list) == {'human'}:
        
        # now check to see if the node above includes terminal nodes that are not human 
        up1_hosts_list = []
        up1_tips_list = []
        
        # get terminal nodes for the node up one (parent node)
        for x in clade.up.get_terminals():   
            up1_hosts_list.append(x.attr['host'])
            up1_tips_list.append(x.name)
        
        # if the node above contains not just human samples, then clade should be set to bird_to_human
        if len(set(up1_hosts_list)) > 1:
            host[clade.name] = 'bird_to_human'
            
            if hasattr(clade, "aa_muts") and len(clade.aa_muts['PB2'])!=0:
                for mut in clade.aa_muts['PB2']:
                    bird_to_human.append(mut)
                    d = {}
                    d = {'clade': clade.name}
                    d['transition'] = 'bird_to_human'
                    d['mut'] = mut
                    bird_to_human_d.append(d)
                    mutlist.append(mut)
                mutations[clade.name] = mutlist
            else:
                d = {}
                d = {'clade': clade.name}
                d['transition'] = 'bird_to_human_d'
                d['mut'] = np.NaN
                bird_to_human_d.append(d)
                
        
        # if including the node above still gives you an entirely human clade, then add these mutations to the 
        # human to human count 
        elif set(up1_hosts_list) == {'human'}: 
            host[clade.name] = 'human_to_human'
            
            if hasattr(clade, "aa_muts") and len(clade.aa_muts['PB2'])!=0:
                for mut in clade.aa_muts['PB2']:
                    human_to_human.append(mut)
                    d = {}
                    d = {'clade': clade.name}
                    d['transition'] = 'human_to_human'
                    d['mut'] = mut
                    human_to_human_d.append(d)
                    mutlist.append(mut)
                mutations[clade.name] = mutlist
            else:
                d = {}
                d = {'clade': clade.name}
                d['transition'] = 'human_to_human'
                d['mut'] = np.NaN
                human_to_human_d.append(d)
        
        else:
            print('some other option I havent thought of but need to address')
            print(clade.up, up1_tips_list, up1_hosts_list)
            
    else:
        host[clade.name] = 'bird_to_bird'
        
        if hasattr(clade, "aa_muts") and len(clade.aa_muts['PB2'])!=0:
            for mut in clade.aa_muts['PB2']:
                bird_to_bird.append(mut)
                d = {}
                d = {'clade': clade.name}
                d['transition'] = 'bird_to_bird'
                d['mut'] = mut
                bird_to_bird_d.append(d)
                mutlist.append(mut)
            mutations[clade.name] = mutlist
        else:
            d = {}
            d = {'clade': clade.name}
            d['transition'] = 'bird_to_bird'
            d['mut'] = np.NaN
            bird_to_bird_d.append(d)
```

In [17]:

```
# use Counter to count the number of times each amino acid change is detected in each list; print total unique SNPs
bird_to_bird_count = Counter(bird_to_bird)
bird_to_human_count = Counter(bird_to_human)
human_to_human_count = Counter(human_to_human)
print(len(bird_to_bird_count), len(bird_to_human_count), len(human_to_human_count))

# get a complete list of all of the SNPs identified
all_SNPs = set(bird_to_human + bird_to_bird + human_to_human)

# loop through and count how many times each SNP occurs in each dataset 
all_counts = {}

for a in all_SNPs: 
    if a in bird_to_bird_count:
        b_to_b = bird_to_bird_count[a]
    else:
        b_to_b = 0
    
    if a in bird_to_human_count:
        b_to_h = bird_to_human_count[a]
    else:
        b_to_h = 0
        
    if a in human_to_human_count:
        h_to_h = human_to_human_count[a]   
    else:
        h_to_h = 0
    
    all_counts[a] = {"bird_to_bird":b_to_b, "bird_to_human": b_to_h, "human_to_human": h_to_h}
        
# print(all_counts)  

# write out counts to a file 
counts_outfile = "results/H7N9/h7n9_pb2_snp_counts.txt"

with open(counts_outfile, "w") as outfile:
    header = "\t".join(["amino_acid_change","count_in_human_to_human_transitions","count_in_bird_to_human_transitions","count_in_bird_to_bird_transitions"])
    outfile.write(header + "\n")
    
for a in all_counts: 
    line_to_write = "\t".join([a, str(all_counts[a]["human_to_human"]), str(all_counts[a]["bird_to_human"]), str(all_counts[a]["bird_to_bird"])])
        
    with open(counts_outfile, "a") as outfile: 
        outfile.write(line_to_write + "\n")
```

```
347 217 247
```

In [18]:

```
# Dataframe of all clades and associated mutation and host transition
df = pd.DataFrame(bird_to_bird_d+bird_to_human_d+human_to_human_d)

# Assign site and host to each mutation
df = (df.dropna(subset=['mut'])
      .assign(AASite=df['mut'].str[1:-1],
              AAParent=df['mut'].str[:1],
              AAChild=df['mut'].str[-1:],
             )
     )
df['host'] = df.apply(lambda row:row['transition'].split('_')[-1], axis=1)
df.head()
```

Out[18]:

|  | clade | mut | transition | AASite | AAParent | AAChild | host |
| --- | --- | --- | --- | --- | --- | --- | --- |
| 1 | NODE\_0000502 | V511I | bird\_to\_bird | 511 | V | I | bird |
| 2 | NODE\_0000502 | M535L | bird\_to\_bird | 535 | M | L | bird |
| 3 | NODE\_0000503 | T666M | bird\_to\_bird | 666 | T | M | bird |
| 4 | A/chicken/Wuxi/WX5/2014 | T21A | bird\_to\_bird | 21 | T | A | bird |
| 5 | A/chicken/Wuxi/WX5/2014 | K22A | bird\_to\_bird | 22 | K | A | bird |

#### Compare mutdiffsel values of human vs avian mutations¶

We compared the differential selection values of human versus avian mutations. We might expect human mutations to have high differential selection values (selected for in human over bird).

In [19]:

```
# Plot function
def pairedhorizontalhist(df, metric, metriclabel, 
                         leftcat, leftlabel, rightcat, rightlabel, 
                         p_df):
    
    #Calculate bins
    bins=50
    ylims = (df[metric].min(), df[metric].max())
    a = df[df['host']==leftcat][metric]
    b = df[df['host']==rightcat][metric]
    ahist = np.histogram(a, bins=bins, range=ylims)
    bhist = np.histogram(b, bins=bins, range=ylims)
    binwidth = ahist[1][1] - ahist[1][0]
    binmid = [x + binwidth/2 for x in ahist[1][:-1]]
    xlimmax = np.amax(np.concatenate((ahist[0], bhist[0])))
    xlimmax = xlimmax + xlimmax/10

    fig, axes = plt.subplots(ncols=2, sharey=True, figsize=(4,4))

    ax = axes[0]
    ax.barh(binmid, ahist[0], height=binwidth, align='center')
    ax.set_xlim([0, xlimmax])
    ax.spines['right'].set_visible(False)
    ax.spines['top'].set_visible(False)
    ax.set_title(leftlabel)
    ax.set_ylabel(metriclabel, rotation=0, labelpad=30)
    ax.invert_xaxis()

    ax = axes[1]
    ax.barh(binmid, bhist[0], height=binwidth, align='center')
    ax.set_xlim([0, xlimmax])
    ax.spines['right'].set_visible(False)
    ax.spines['top'].set_visible(False)
    ax.yaxis.set_visible(False)
    ax.spines['left'].set_visible(False)
    ax.set_title(rightlabel)

    plt.subplots_adjust(wspace=0.05, hspace=0)
    
#     # Adding legend, p value annotation
    p_dfsub = p_df[p_df['metric'] == metric]
    for nrow, (index, row) in enumerate(p_dfsub.iterrows()):
        annotstring = ('{0} >={1}: p = {2:1.2e} / {3:1.2e}'\
                       .format(metric, row['threshold'], row['fisher_p'], row['chi2_p']))
        axes[1].annotate(annotstring,
                         xycoords='axes fraction',
                         xy=(0.15, 1-0.1-0.05*nrow),
                        )
    fig.savefig('results/H7N9/pairedhist.pdf', bbox_inches='tight', dpi=300)
    return ahist, bhist

# Calculate p value function
def calc_p_vals(df, pairstotest, metrics, threshold):
    """
    Makes 2x2 contingency tables of mutations occurring in strain1 vs strain2, 
    against whether a mutation scores high or low on given metric (by my arbitrary threshold)
    Returns dataframe of p vals.
    """
    p_rowlist = []
    for (strain1, strain2) in pairstotest:
        for metric in metrics:
            for i in threshold[metric]:
                hi1 = len( df[(df['host']==strain1) & (df[metric]>=i)] )
                lo1 = len( df[(df['host']==strain1) & (df[metric]<i)] )
                hi2 = len( df[(df['host']==strain2) & (df[metric]>=i)] )
                lo2 = len( df[(df['host']==strain2) & (df[metric]<i)] )

                table = np.array([[hi1, hi2],
                                  [lo1, lo2]])
    #             for printing 2x2 contingency table
#                 pdtable = pd.DataFrame(table, columns=[strain1, strain2], 
#                                        index=[metric+' >={0}'.format(i), metric+' <{0}'.format(i)])
#                 print(pdtable)

                p_dict = {}
                p_dict['strain1'] = strain1
                p_dict['strain2'] = strain2
                p_dict['metric'] = metric
                p_dict['threshold'] = i
                p_dict['strain1_high_count'] = hi1
                p_dict['strain1_low_count'] = lo1
                p_dict['strain2high_count'] = hi2
                p_dict['strain2_low_count'] = lo2
                fisher_or, p_dict['fisher_p'] = stats.fisher_exact(table)
                chi2, p_dict['chi2_p'], chi2_dof, chi2_expected = stats.chi2_contingency(table)
                g, p_dict['g_p'], g_dof, g_expected = stats.chi2_contingency(table, lambda_="log-likelihood")
                p_rowlist.append(p_dict)

    #             print('fisher: {0}\nchi2: {1}\ng-test: {2}'\
    #                  .format(p_dict['fisher_p'], p_dict['chi2_p'], p_dict['g_p']))

    p_df = pd.DataFrame(p_rowlist)
    p_df = p_df[['strain1', 'strain2', 'metric', 'threshold',
                 'strain1_high_count', 'strain1_low_count', 'strain2_low_count', 'strain2high_count', 
                 'fisher_p', 'chi2_p', 'g_p']]
    return p_df
```

In [20]:

```
# Gather metrics from DMS experiment
dmssummarydf = pd.read_csv('results/diffsel/summary_prefs_effects_diffsel.csv')
dmssummarydf.head()

# Merge H7N9 mutations with DMS data
df.AASite = df.AASite.astype('int64')
dmssummarydf.site = dmssummarydf.site.astype('int64')
H7N9andDMSdf = (pd.merge(df, dmssummarydf, how='left', 
                        left_on=['AASite', 'AAChild'], right_on=['site', 'mutation']))

H7N9andDMSdf.head()
```

Out[20]:

|  | clade | mut | transition | AASite | AAParent | AAChild | host | site | wildtype | mutation | ... | prefCCL141 | log2prefA549 | log2prefCCL141 | effectA549 | effectCCL141 | log2effectA549 | log2effectCCL141 | mutdiffsel | Known human adaptive | Experimentally adaptive in |
| --- | --- | --- | --- | --- | --- | --- | --- | --- | --- | --- | --- | --- | --- | --- | --- | --- | --- | --- | --- | --- | --- |
| 0 | NODE\_0000502 | V511I | bird\_to\_bird | 511 | V | I | bird | 511 | V | I | ... | 0.104747 | -3.040447 | -3.255012 | 0.563084 | 0.941541 | -0.828578 | -0.086904 | -0.453744 | No | None |
| 1 | NODE\_0000502 | M535L | bird\_to\_bird | 535 | M | L | bird | 535 | M | L | ... | 0.086971 | -4.298134 | -3.523316 | 0.280786 | 0.473019 | -1.832458 | -1.080029 | -0.579487 | No | None |
| 2 | NODE\_0000503 | T666M | bird\_to\_bird | 666 | T | M | bird | 666 | T | M | ... | 0.040190 | -5.151229 | -4.637026 | 0.301605 | 0.320407 | -1.729269 | -1.642024 | -0.194933 | No | None |
| 3 | A/chicken/Wuxi/WX5/2014 | T21A | bird\_to\_bird | 21 | T | A | bird | 21 | T | A | ... | 0.034397 | -4.758455 | -4.861560 | 0.404254 | 0.614440 | -1.306667 | -0.702656 | -0.193902 | No | None |
| 4 | A/chicken/Wuxi/WX5/2014 | K22A | bird\_to\_bird | 22 | K | A | bird | 22 | K | A | ... | 0.046353 | -4.098906 | -4.431185 | 1.134356 | 0.491972 | 0.181873 | -1.023352 | 0.737047 | No | None |

5 rows × 21 columns

In [21]:

```
# Test for significance of enrichment
df = H7N9andDMSdf
strainpairs = [('human', 'bird')]
metrics = ['mutdiffsel']
threshold = {'mutdiffsel': [0.5, 1]}
p_df = calc_p_vals(df, strainpairs, metrics, threshold)
p_df
```

Out[21]:

|  | strain1 | strain2 | metric | threshold | strain1\_high\_count | strain1\_low\_count | strain2\_low\_count | strain2high\_count | fisher\_p | chi2\_p | g\_p |
| --- | --- | --- | --- | --- | --- | --- | --- | --- | --- | --- | --- |
| 0 | human | bird | mutdiffsel | 0.5 | 302 | 455 | 377 | 132 | 2.691687e-07 | 3.956573e-07 | 3.082381e-07 |
| 1 | human | bird | mutdiffsel | 1.0 | 106 | 651 | 466 | 43 | 2.475672e-03 | 3.519694e-03 | 2.979258e-03 |

In [22]:

```
# Plot distribution of mutations
dataframe = H7N9andDMSdf
metric = 'mutdiffsel'
metriclabel = 'Mutational\ndifferential\nselection'
leftcat, leftlabel = 'bird', 'Avian'
rightcat, rightlabel = 'human', 'Human'

hist_avian, hist_human = pairedhorizontalhist(dataframe, metric, metriclabel, 
                                              leftcat, leftlabel, rightcat, rightlabel, 
                                              p_df)
```

In [23]:

```
# What are the human mutations with highest mutdiffsel and counts?
tempdf = (H7N9andDMSdf.groupby(['mut', 'host', 'mutdiffsel'])
             .size().reset_index(name='counts')
             .sort_values('counts', ascending=False)
            )
tempdf_bird = tempdf[tempdf['host']=='bird'].drop(columns=['host'])
tempdf_human = tempdf[tempdf['host']=='human'].drop(columns='host')
tempdf = (pd.merge(tempdf_human, tempdf_bird, how='outer', on=['mut', 'mutdiffsel'], 
                  suffixes=['_human', '_bird']
                 )
          .fillna(0)
          .assign(counts_diff=lambda x: x['counts_human']-x['counts_bird'])
         )
tempdf[(tempdf['counts_human']>3) &
       (tempdf['mutdiffsel']>1)]
```

Out[23]:

|  | mut | mutdiffsel | counts\_human | counts\_bird | counts\_diff |
| --- | --- | --- | --- | --- | --- |
| 1 | D701N | 2.240536 | 40.0 | 2.0 | 38.0 |
| 2 | S534F | 2.036796 | 14.0 | 2.0 | 12.0 |
| 5 | E627V | 1.277026 | 8.0 | 1.0 | 7.0 |
| 11 | R355K | 1.093163 | 5.0 | 4.0 | 1.0 |
| 26 | G685R | 1.014202 | 4.0 | 0.0 | 4.0 |
| 28 | T521I | 3.165122 | 4.0 | 0.0 | 4.0 |

In [24]:

```
# Get mutations in each bin for annotating the histogram

# First get ranges for each bin
binranges = []
for (i, x) in enumerate(hist_avian[1][:-1]):
    binranges.append((hist_avian[1][i], hist_avian[1][i+1]))
# find bins where there is large difference in count
hist_diff = hist_human[0] - hist_avian[0]
bins_diff = dict(zip(binranges, hist_diff))

selectedbins = []
for binrange, diff in bins_diff.items():
    if binrange[1]>0.5:
        selectedbins.append((binrange, diff))
        
for (binrange, diff) in reversed(selectedbins):
    print('bin: ' + str(binrange[0]) + '-' + str(binrange[1]) + '\nbin_count_diff: ' + str(diff))
    dfsub = (tempdf[(tempdf['mutdiffsel']>=binrange[0]) & 
                    (tempdf['mutdiffsel']<binrange[1])]
             .sort_values('counts_diff', ascending=False)
            )
    display(HTML(dfsub.to_html()))
```

```
bin: 3.061650865136176-3.1741649020582683
bin_count_diff: 5
```

|  | mut | mutdiffsel | counts\_human | counts\_bird | counts\_diff |
| --- | --- | --- | --- | --- | --- |
| 28 | T521I | 3.165122 | 4.0 | 0.0 | 4.0 |

```
bin: 2.9491368282140837-3.061650865136176
bin_count_diff: 1
```

|  | mut | mutdiffsel | counts\_human | counts\_bird | counts\_diff |
| --- | --- | --- | --- | --- | --- |
| 268 | A274V | 3.009671 | 1.0 | 0.0 | 1.0 |

```
bin: 2.8366227912919917-2.9491368282140837
bin_count_diff: 0
```

|  | mut | mutdiffsel | counts\_human | counts\_bird | counts\_diff |
| --- | --- | --- | --- | --- | --- |

```
bin: 2.7241087543698987-2.8366227912919917
bin_count_diff: 0
```

|  | mut | mutdiffsel | counts\_human | counts\_bird | counts\_diff |
| --- | --- | --- | --- | --- | --- |

```
bin: 2.6115947174478067-2.7241087543698987
bin_count_diff: 0
```

|  | mut | mutdiffsel | counts\_human | counts\_bird | counts\_diff |
| --- | --- | --- | --- | --- | --- |

```
bin: 2.4990806805257146-2.6115947174478067
bin_count_diff: 3
```

|  | mut | mutdiffsel | counts\_human | counts\_bird | counts\_diff |
| --- | --- | --- | --- | --- | --- |
| 57 | T271A | 2.520153 | 2.0 | 0.0 | 2.0 |
| 140 | P430T | 2.551779 | 1.0 | 0.0 | 1.0 |

```
bin: 2.3865666436036226-2.4990806805257146
bin_count_diff: 0
```

|  | mut | mutdiffsel | counts\_human | counts\_bird | counts\_diff |
| --- | --- | --- | --- | --- | --- |

```
bin: 2.2740526066815305-2.3865666436036226
bin_count_diff: 0
```

|  | mut | mutdiffsel | counts\_human | counts\_bird | counts\_diff |
| --- | --- | --- | --- | --- | --- |

```
bin: 2.1615385697594376-2.2740526066815305
bin_count_diff: 38
```

|  | mut | mutdiffsel | counts\_human | counts\_bird | counts\_diff |
| --- | --- | --- | --- | --- | --- |
| 1 | D701N | 2.240536 | 40.0 | 2.0 | 38.0 |

```
bin: 2.0490245328373455-2.1615385697594376
bin_count_diff: 0
```

|  | mut | mutdiffsel | counts\_human | counts\_bird | counts\_diff |
| --- | --- | --- | --- | --- | --- |
| 288 | L708F | 2.109105 | 1.0 | 0.0 | 1.0 |
| 350 | K339E | 2.067135 | 1.0 | 1.0 | 0.0 |
| 337 | I260V | 2.133250 | 1.0 | 2.0 | -1.0 |

```
bin: 1.9365104959152535-2.0490245328373455
bin_count_diff: 14
```

|  | mut | mutdiffsel | counts\_human | counts\_bird | counts\_diff |
| --- | --- | --- | --- | --- | --- |
| 2 | S534F | 2.036796 | 14.0 | 2.0 | 12.0 |
| 212 | S489P | 2.041974 | 1.0 | 0.0 | 1.0 |
| 325 | L183S | 1.947803 | 1.0 | 0.0 | 1.0 |

```
bin: 1.8239964589931614-1.9365104959152535
bin_count_diff: -1
```

|  | mut | mutdiffsel | counts\_human | counts\_bird | counts\_diff |
| --- | --- | --- | --- | --- | --- |
| 151 | Q522L | 1.916284 | 1.0 | 0.0 | 1.0 |
| 371 | I647V | 1.905366 | 0.0 | 2.0 | -2.0 |

```
bin: 1.7114824220710694-1.8239964589931614
bin_count_diff: -2
```

|  | mut | mutdiffsel | counts\_human | counts\_bird | counts\_diff |
| --- | --- | --- | --- | --- | --- |
| 125 | R355G | 1.742070 | 1.0 | 2.0 | -1.0 |
| 481 | E304V | 1.712828 | 0.0 | 1.0 | -1.0 |

```
bin: 1.5989683851489764-1.7114824220710694
bin_count_diff: -1
```

|  | mut | mutdiffsel | counts\_human | counts\_bird | counts\_diff |
| --- | --- | --- | --- | --- | --- |
| 542 | N148K | 1.685194 | 0.0 | 1.0 | -1.0 |

```
bin: 1.4864543482268844-1.5989683851489764
bin_count_diff: -4
```

|  | mut | mutdiffsel | counts\_human | counts\_bird | counts\_diff |
| --- | --- | --- | --- | --- | --- |
| 490 | G150S | 1.519233 | 0.0 | 1.0 | -1.0 |
| 523 | D309G | 1.521226 | 0.0 | 1.0 | -1.0 |
| 357 | K121R | 1.566798 | 1.0 | 3.0 | -2.0 |

```
bin: 1.3739403113047923-1.4864543482268844
bin_count_diff: -1
```

|  | mut | mutdiffsel | counts\_human | counts\_bird | counts\_diff |
| --- | --- | --- | --- | --- | --- |
| 188 | S714N | 1.449092 | 1.0 | 1.0 | 0.0 |
| 433 | T129A | 1.406297 | 0.0 | 1.0 | -1.0 |

```
bin: 1.2614262743826998-1.3739403113047923
bin_count_diff: 8
```

|  | mut | mutdiffsel | counts\_human | counts\_bird | counts\_diff |
| --- | --- | --- | --- | --- | --- |
| 5 | E627V | 1.277026 | 8.0 | 1.0 | 7.0 |
| 84 | K627V | 1.277026 | 2.0 | 0.0 | 2.0 |
| 504 | A174T | 1.327523 | 0.0 | 1.0 | -1.0 |

```
bin: 1.1489122374606078-1.2614262743826998
bin_count_diff: 2
```

|  | mut | mutdiffsel | counts\_human | counts\_bird | counts\_diff |
| --- | --- | --- | --- | --- | --- |
| 30 | D256G | 1.224898 | 3.0 | 1.0 | 2.0 |
| 94 | A674T | 1.244557 | 2.0 | 1.0 | 1.0 |
| 97 | V648A | 1.174646 | 2.0 | 1.0 | 1.0 |
| 180 | T224S | 1.186035 | 1.0 | 0.0 | 1.0 |
| 301 | L281V | 1.216959 | 1.0 | 0.0 | 1.0 |
| 450 | T23A | 1.227517 | 0.0 | 1.0 | -1.0 |
| 471 | E677D | 1.184112 | 0.0 | 1.0 | -1.0 |
| 395 | D309N | 1.197917 | 0.0 | 2.0 | -2.0 |

```
bin: 1.0363982005385157-1.1489122374606078
bin_count_diff: -2
```

|  | mut | mutdiffsel | counts\_human | counts\_bird | counts\_diff |
| --- | --- | --- | --- | --- | --- |
| 11 | R355K | 1.093163 | 5.0 | 4.0 | 1.0 |
| 199 | R589K | 1.105742 | 1.0 | 0.0 | 1.0 |
| 244 | I141M | 1.116203 | 1.0 | 0.0 | 1.0 |
| 314 | N456D | 1.117627 | 1.0 | 0.0 | 1.0 |
| 400 | R369K | 1.132212 | 0.0 | 1.0 | -1.0 |
| 453 | R8G | 1.045575 | 0.0 | 1.0 | -1.0 |
| 455 | V272I | 1.055952 | 0.0 | 1.0 | -1.0 |
| 466 | V172L | 1.059193 | 0.0 | 1.0 | -1.0 |
| 491 | G173E | 1.102795 | 0.0 | 1.0 | -1.0 |
| 494 | G416S | 1.081125 | 0.0 | 1.0 | -1.0 |

```
bin: 0.9238841636164232-1.0363982005385157
bin_count_diff: 0
```

|  | mut | mutdiffsel | counts\_human | counts\_bird | counts\_diff |
| --- | --- | --- | --- | --- | --- |
| 26 | G685R | 1.014202 | 4.0 | 0.0 | 4.0 |
| 132 | Q566H | 1.000676 | 1.0 | 0.0 | 1.0 |
| 166 | T117N | 0.978627 | 1.0 | 0.0 | 1.0 |
| 260 | A674V | 0.947049 | 1.0 | 0.0 | 1.0 |
| 300 | L193F | 1.035545 | 1.0 | 1.0 | 0.0 |
| 397 | R15L | 0.964545 | 0.0 | 1.0 | -1.0 |
| 442 | T637A | 0.995078 | 0.0 | 1.0 | -1.0 |
| 460 | S155G | 0.979977 | 0.0 | 1.0 | -1.0 |
| 478 | E192D | 0.994182 | 0.0 | 1.0 | -1.0 |
| 507 | A274T | 0.930703 | 0.0 | 1.0 | -1.0 |
| 163 | T178A | 1.031231 | 1.0 | 3.0 | -2.0 |

```
bin: 0.8113701266943312-0.9238841636164232
bin_count_diff: 9
```

|  | mut | mutdiffsel | counts\_human | counts\_bird | counts\_diff |
| --- | --- | --- | --- | --- | --- |
| 24 | I147T | 0.886417 | 4.0 | 0.0 | 4.0 |
| 40 | V598A | 0.892716 | 3.0 | 0.0 | 3.0 |
| 86 | E681D | 0.911252 | 2.0 | 0.0 | 2.0 |
| 108 | D153N | 0.867341 | 2.0 | 0.0 | 2.0 |
| 283 | D195E | 0.823652 | 1.0 | 0.0 | 1.0 |
| 295 | M467I | 0.890967 | 1.0 | 0.0 | 1.0 |
| 351 | K586R | 0.821765 | 1.0 | 0.0 | 1.0 |
| 192 | R8K | 0.832167 | 1.0 | 1.0 | 0.0 |
| 198 | R62G | 0.865023 | 1.0 | 1.0 | 0.0 |
| 413 | V89L | 0.818129 | 0.0 | 1.0 | -1.0 |
| 472 | E700Q | 0.869941 | 0.0 | 1.0 | -1.0 |
| 560 | K339N | 0.910591 | 0.0 | 1.0 | -1.0 |
| 390 | V606I | 0.822475 | 0.0 | 2.0 | -2.0 |

```
bin: 0.6988560897722387-0.8113701266943312
bin_count_diff: -9
```

|  | mut | mutdiffsel | counts\_human | counts\_bird | counts\_diff |
| --- | --- | --- | --- | --- | --- |
| 48 | V588I | 0.736181 | 3.0 | 1.0 | 2.0 |
| 58 | I147V | 0.762469 | 2.0 | 1.0 | 1.0 |
| 80 | K214R | 0.777570 | 2.0 | 3.0 | -1.0 |
| 255 | A588V | 0.722098 | 1.0 | 2.0 | -1.0 |
| 420 | Q591P | 0.755394 | 0.0 | 1.0 | -1.0 |
| 432 | T178S | 0.775291 | 0.0 | 1.0 | -1.0 |
| 434 | T666S | 0.776814 | 0.0 | 1.0 | -1.0 |
| 500 | A674P | 0.770366 | 0.0 | 1.0 | -1.0 |
| 516 | D9G | 0.712415 | 0.0 | 1.0 | -1.0 |
| 551 | I539V | 0.793065 | 0.0 | 1.0 | -1.0 |
| 556 | I647M | 0.777781 | 0.0 | 1.0 | -1.0 |
| 561 | K22A | 0.737047 | 0.0 | 1.0 | -1.0 |
| 396 | V457I | 0.727562 | 0.0 | 2.0 | -2.0 |

```
bin: 0.5863420528501466-0.6988560897722387
bin_count_diff: 11
```

|  | mut | mutdiffsel | counts\_human | counts\_bird | counts\_diff |
| --- | --- | --- | --- | --- | --- |
| 6 | T117P | 0.591799 | 7.0 | 4.0 | 3.0 |
| 37 | K5R | 0.692851 | 3.0 | 0.0 | 3.0 |
| 21 | E249G | 0.593593 | 4.0 | 2.0 | 2.0 |
| 215 | S36T | 0.663448 | 1.0 | 0.0 | 1.0 |
| 358 | I676M | 0.651792 | 1.0 | 0.0 | 1.0 |
| 349 | K32R | 0.637225 | 1.0 | 0.0 | 1.0 |
| 298 | L668F | 0.635628 | 1.0 | 0.0 | 1.0 |
| 269 | A278V | 0.596038 | 1.0 | 0.0 | 1.0 |
| 258 | A674E | 0.677071 | 1.0 | 0.0 | 1.0 |
| 237 | F446L | 0.693528 | 1.0 | 0.0 | 1.0 |
| 264 | A44V | 0.677087 | 1.0 | 0.0 | 1.0 |
| 173 | V109A | 0.643203 | 1.0 | 0.0 | 1.0 |
| 170 | T106I | 0.682917 | 1.0 | 0.0 | 1.0 |
| 157 | V676M | 0.651792 | 1.0 | 0.0 | 1.0 |
| 103 | E188K | 0.630101 | 2.0 | 1.0 | 1.0 |
| 169 | T117A | 0.616568 | 1.0 | 1.0 | 0.0 |
| 275 | D740N | 0.613293 | 1.0 | 1.0 | 0.0 |
| 120 | V613I | 0.634035 | 1.0 | 1.0 | 0.0 |
| 323 | M64I | 0.682433 | 1.0 | 1.0 | 0.0 |
| 361 | K116R | 0.626371 | 1.0 | 1.0 | 0.0 |
| 74 | R508K | 0.619528 | 2.0 | 3.0 | -1.0 |
| 461 | S629T | 0.594134 | 0.0 | 1.0 | -1.0 |
| 463 | S497N | 0.641672 | 0.0 | 1.0 | -1.0 |
| 477 | E241K | 0.695242 | 0.0 | 1.0 | -1.0 |
| 501 | A684V | 0.675796 | 0.0 | 1.0 | -1.0 |
| 383 | R3G | 0.681648 | 0.0 | 2.0 | -2.0 |
| 386 | N711S | 0.626451 | 0.0 | 2.0 | -2.0 |

```
bin: 0.47382801592805457-0.5863420528501466
bin_count_diff: 99
```

|  | mut | mutdiffsel | counts\_human | counts\_bird | counts\_diff |
| --- | --- | --- | --- | --- | --- |
| 0 | E627K | 0.546131 | 119.0 | 15.0 | 104.0 |
| 36 | K482R | 0.580741 | 3.0 | 1.0 | 2.0 |
| 292 | M28I | 0.585033 | 1.0 | 0.0 | 1.0 |
| 347 | I66L | 0.490383 | 1.0 | 0.0 | 1.0 |
| 308 | M756V | 0.521275 | 1.0 | 0.0 | 1.0 |
| 304 | L607V | 0.513341 | 1.0 | 0.0 | 1.0 |
| 257 | A661S | 0.573675 | 1.0 | 0.0 | 1.0 |
| 127 | V588L | 0.516992 | 1.0 | 0.0 | 1.0 |
| 56 | G590S | 0.485203 | 2.0 | 1.0 | 1.0 |
| 131 | T569A | 0.512636 | 1.0 | 1.0 | 0.0 |
| 16 | K702R | 0.543500 | 4.0 | 4.0 | 0.0 |
| 317 | M66L | 0.490383 | 1.0 | 1.0 | 0.0 |
| 134 | Q73K | 0.521685 | 1.0 | 2.0 | -1.0 |
| 175 | T105A | 0.549533 | 1.0 | 2.0 | -1.0 |
| 226 | E65K | 0.480535 | 1.0 | 2.0 | -1.0 |
| 83 | K61R | 0.489419 | 2.0 | 3.0 | -1.0 |
| 475 | E391G | 0.515636 | 0.0 | 1.0 | -1.0 |
| 488 | F610L | 0.506308 | 0.0 | 1.0 | -1.0 |
| 498 | A624S | 0.521452 | 0.0 | 1.0 | -1.0 |
| 548 | M53V | 0.557448 | 0.0 | 1.0 | -1.0 |
| 559 | K32T | 0.568145 | 0.0 | 1.0 | -1.0 |
| 60 | I451V | 0.526788 | 2.0 | 4.0 | -2.0 |
| 367 | I529V | 0.534560 | 0.0 | 3.0 | -3.0 |

#### Plot tree to visualize where mutations are occurring¶

In [25]:

```
#Code adapted from Baltic/Augur pipeline for tree plotting

def plot_tree(tree, figure_name, tip_size, branchlen_attr,
                  color_branches_list, branchcol, color_circles_list, circlecol, color_circles_list2, circlecol2):
    """Plot a BioPython Phylo tree in the BALTIC-style.
    """
    # Plot tree in BALTIC style from Bio.Phylo tree.
    mpl.rcParams['savefig.dpi'] = 300 #120
    mpl.rcParams['figure.dpi'] = 100

    mpl.rcParams['font.weight']= 1 #300
    mpl.rcParams['axes.labelweight']= 1 #300
    mpl.rcParams['font.size']= 8 #14
    mpl.rcParams['font.sans-serif'] = "Arial"
    mpl.rcParams['font.family'] = "sans-serif"

    yvalues = [node.yvalue for node in tree.find_clades()]
    y_span = max(yvalues)
    y_unit = y_span / float(len(yvalues))

    #
    # Setup the figure grid.
    #

    fig = plt.figure(figsize=(2, 6), facecolor='w') #figsize=(3, 3)
    gs = gridspec.GridSpec(2, 1, height_ratios=[14, 1], width_ratios=[1], hspace=0.1, wspace=0.1)
    ax = fig.add_subplot(gs[0])

    L=len([k for k in tree.find_clades() if k.is_terminal()])

    # Setup arrays for tip and internal node coordinates.
    tip_circles_x = []
    tip_circles_y = []
    tip_circles_color = []
    tip_circle_sizes = []
    node_circles_x = []
    node_circles_y = []
    node_circles_color = []
    node_circle_sizes = []
    node_line_widths = []
    node_line_segments = []
    node_line_colors = []
    branch_line_segments = []
    branch_line_widths = []
    branch_line_colors = []
    branch_line_segments_selected = []
    branch_line_widths_selected = []
    branch_line_colors_selected = []

    for k in tree.find_clades(): ## iterate over objects in tree
        x=k.attr[branchlen_attr] ## or from x position determined earlier
        y=k.yvalue ## get y position from .drawTree that was run earlier, but could be anything else

        if k.up is None:
            xp = None
        else:
            xp=k.up.attr[branchlen_attr] ## get x position of current object's parent

        if x==None: ## matplotlib won't plot Nones, like root
            x=0.0
        if xp==None:
            xp=x

        c = 'silver'
        if k.name in color_branches_list:
            c = branchcol

        branchWidth=0.5 #2
    
        if k.is_terminal(): ## if leaf...
            if k.name in color_circles_list:
                s = tip_size ## tip size can be fixed
                ccol = circlecol
            elif k.name in color_circles_list2:
                s = tip_size ## tip size can be fixed
                ccol = circlecol2
            else:
                s = 0
                ccol = c
            tip_circle_sizes.append(s)
            tip_circles_x.append(x)
            tip_circles_y.append(y)
            tip_circles_color.append(ccol)
        else: ## if node...
            if k.name in color_circles_list:
                s = tip_size ## tip size can be fixed
                ccol = circlecol
            elif k.name in color_circles_list2:
                s = tip_size ## tip size can be fixed
                ccol = circlecol2
            else:
                s = 0
                ccol = c
            node_circle_sizes.append(s)
            node_circles_x.append(x)#
            node_circles_y.append(y)#
            node_circles_color.append(ccol)#

            if len(k.clades)==1:
                node_circles_x.append(x)
                node_circles_y.append(y)
                node_circles_color.append(circlecol)
            
            ax.plot([x,x],[k.clades[-1].yvalue, k.clades[0].yvalue], lw=branchWidth, color=c, ls='-', zorder=9, solid_capstyle='round')
            
        if k.name in color_branches_list:
            branch_line_segments_selected.append([(xp, y), (x, y)])
            branch_line_widths_selected.append(branchWidth)
            branch_line_colors_selected.append(c)
        else:
            branch_line_segments.append([(xp, y), (x, y)])
            branch_line_widths.append(branchWidth)
            branch_line_colors.append(c)

    branch_lc = LineCollection(branch_line_segments, zorder=9)
    branch_lc.set_color(branch_line_colors)
    branch_lc.set_linewidth(branch_line_widths)
    branch_lc.set_linestyle("-")
    ax.add_collection(branch_lc)
    
    branch_lc_selected = LineCollection(branch_line_segments_selected, zorder=10)
    branch_lc_selected.set_color(branch_line_colors_selected)
    branch_lc_selected.set_linewidth(branch_line_widths_selected)
    branch_lc_selected.set_linestyle("-")
    ax.add_collection(branch_lc_selected)

    # Add circles for tips and internal nodes.
    tip_circle_sizes = np.array(tip_circle_sizes)
    ax.scatter(tip_circles_x, tip_circles_y, s=tip_circle_sizes, facecolor=tip_circles_color, edgecolor='none',zorder=12) ## plot circle for every tip
    ax.scatter(node_circles_x, node_circles_y, s=node_circle_sizes, facecolor=node_circles_color, edgecolor='none', zorder=11)

    ax.spines['top'].set_visible(False) ## no axes
    ax.spines['right'].set_visible(False)
    ax.spines['left'].set_visible(False)

#     ax.grid(axis='x',ls='-',color='silver', linewidth=0.5)
    ax.tick_params(axis='y',size=0)
    ax.set_yticklabels([])
    if branchlen_attr=='num_date':
        ax.set_xticks([2006,2009,2012,2015,2018], minor=False)

    gs.tight_layout(fig)
    plt.savefig(figure_name)
```

In [26]:

```
humanhosts = ['human_to_human', 'bird_to_human']
avianhosts = ['bird_to_bird']
humanclades = []
for clade in tree.find_clades():
    try:
        if (host[clade.name] in humanhosts):
            humanclades.append(clade.name)
    except:
        pass
print(len(humanclades))
```

```
1465
```

In [27]:

```
muttoplotlist = ['E627K', 'D701N', 'E627V', 'S534F', 'R355K', 'R355G', 'T521I', 'G685R']
mutonclades = {}
for mut in muttoplotlist:
    mutonclades[mut] = []
    for clade in tree.find_clades():
        if hasattr(clade, "aa_muts") and len(clade.aa_muts['PB2'])!=0:
            if mut in clade.aa_muts['PB2']:
                mutonclades[mut].append(clade.name)
    print(mut, len(mutonclades[mut]))
```

```
E627K 134
D701N 42
E627V 9
S534F 16
R355K 9
R355G 3
T521I 4
G685R 4
```

In [18]:

```
# Plot the tree. 627 site.
branchlenattributes = ['num_date', 'div']
for branchlen_attr in branchlenattributes:
    plot_tree(
        tree,
        'results/H7N9/baltictreelong_{0}_{1}.pdf'.format(branchlen_attr, '627'),
        10,
        branchlen_attr,
        humanclades, 'k',
        mutonclades['E627K'], 'orange',
        mutonclades['E627V'], 'red',
    )
```

In [19]:

```
# Plot the tree. 355 site.
branchlenattributes = ['num_date', 'div']
for branchlen_attr in branchlenattributes:
    plot_tree(
        tree,
        'results/H7N9/baltictreelong_{0}_{1}.pdf'.format(branchlen_attr, '355'),
        10,
        branchlen_attr,
        humanclades, 'k',
        mutonclades['R355K'], 'orange',
        mutonclades['R355G'], 'red',
    )
```

In [20]:

```
# Plot the tree. other sites.
branchlenattributes = ['num_date', 'div']
for mut in ['D701N', 'S534F', 'R355K', 'T521I', 'G685R']:
    for branchlen_attr in branchlenattributes:
        plot_tree(
            tree,
            'results/H7N9/baltictreelong_{0}_{1}.pdf'.format(branchlen_attr, mut),
            10,
            branchlen_attr,
            humanclades, 'k',
            mutonclades[mut], 'orange',
            [], 'red',
        )
```

Our above phylogenetic analyses likely underestimate the true number of mutation counts in human hosts. Due to undersampling of avian sequences, human sequences often form a monophyletic clade. If a mutation occurs recurrently in that human clade, the most parsimonious phylogenetic model will infer that the mutation occurred only once. However, in reality, it is more likely that the mutation occurred *x* recurrent times, since H7N9 infections of humans tend to be dead end.

To try to correct for that, here we take a non-phylogenetic approach and simply count the number of times a mutation occurs in human versus avian hosts. To do so, we first align of all sequences on the tips of the H7N9 PB2 tree. We then simply count the number of amino acid variants found in human versus avian hosts. We then conduct a Fisher's exact test to see if the variants are at significantly different frequencies in human versus avian hosts.

We find that the 4 of the 5 mutations with high mutation differential selection that we identified by phylogenetic analysis are indeed statistically enriched in human over avian hosts. The one mutation that does not reach statistical significance nonetheless occurs 6 times in humans and none in bird.

In [28]:

```
# Read in sequences to df
aaseqs = SeqIO.parse('data/aa_alignment_h7n9_pb2.fasta', 'fasta') # same set as in as tree
aalist = []
for rec in aaseqs:
    seqlist = [rec.id] + list(rec.seq)
    aalist.append(seqlist)
df = pd.DataFrame(aalist)
df.columns = ['strain'] + list(range(1, 761))

# Read in metadata to annotate host species
metadata = (pd.read_table('data/metadata_h7n9_pb2.tsv')[['strain', 'host']]
            .replace(['anascarolinensis', 'anascrecca', 'turkey'], 'avian')
           )
df = pd.merge(df, metadata, on='strain')
df = df.query('host=="avian" or host=="human"')

print(len(df))

# # Convert to longform
df = pd.melt(df, id_vars=['strain', 'host'], value_vars=list(range(1, 761)), 
             var_name='site', value_name='AA')

# # Group by site and host
dfgroup = (df.groupby(['site','host','AA']).count()
           .rename(columns={'strain':'count'})
           .reset_index()
          )
# # Example
dfgroup[dfgroup['site']==627]
```

```
1391
```

Out[28]:

|  | site | host | AA | count |
| --- | --- | --- | --- | --- |
| 1787 | 627 | avian | E | 492 |
| 1788 | 627 | avian | K | 3 |
| 1789 | 627 | human | E | 302 |
| 1790 | 627 | human | K | 561 |
| 1791 | 627 | human | Q | 1 |
| 1792 | 627 | human | V | 32 |

In [29]:

```
# Function for site-wise Fisher's exact test

# Import rpy2 to use R's fisher's exact test which allows tests on m x n tables
# https://stackoverflow.com/questions/25368284/fishers-exact-test-for-bigger-than-2-by-2-contingency-table
# SciPy's fisher's exact works only on 2 x 2 tables
import rpy2.robjects.numpy2ri
from rpy2.robjects.packages import importr
rpy2.robjects.numpy2ri.activate()
stats = importr('stats')

def sitewiseFishers(dfgroup, sites, mincount):
    fishersresdict = {
        'site': [],
        'AAvars': [],
        'table': [],
        'fishersp': []
    }
    for i in sites:
        dfsite = (dfgroup[dfgroup['site']==i]
                  .sort_values(['host', 'count'], ascending=[True, False]) #Sort to get most frequent avian variants first
                 )
        dfsitetotalcount = (dfsite[['AA', 'count']]
                            .groupby(['AA'], sort=False).sum() #Turn off sort in groupby to keep above sort order
                            .reset_index()
                            .query('count >={0}'.format(mincount)) #Require that variant has mincount
                           )
        AAvars = dfsitetotalcount['AA'].unique()
        if len(AAvars)>1:
            table = []
            for aa in AAvars:
                try:
                    avian_count = int(dfsite[(dfsite['host']=='avian') & (dfsite['AA']==aa)]['count'])
                except:
                    avian_count = 0
                try:
                    human_count = int(dfsite[(dfsite['host']=='human') & (dfsite['AA']==aa)]['count'])
                except:
                    human_count = 0
                table.append([avian_count, human_count])
            # SciPy Fisher's exact -- up to only 2x2
    #         oddsratio, pval = stats.fisher_exact(table)
            # R's Fisher's exact -- m x n
            table = np.array(table)
            res = stats.fisher_test(table)
            pval = res[0][0]

            fishersresdict['site'].append(i)
            fishersresdict['AAvars'].append(AAvars)
            fishersresdict['table'].append(table)
            fishersresdict['fishersp'].append(pval)
    dffishers = pd.DataFrame.from_dict(fishersresdict)
    return dffishers
```

In [30]:

```
chosensites = [627, 701, 534, 355, 521]
dffishers_chosen = sitewiseFishers(dfgroup, chosensites, 6)
dffishers_chosen
```

Out[30]:

|  | site | AAvars | table | fishersp |
| --- | --- | --- | --- | --- |
| 0 | 627 | [E, K, V] | [[492, 302], [3, 561], [0, 32]] | 4.050262e-156 |
| 1 | 701 | [D, N] | [[494, 837], [1, 59]] | 9.822496e-11 |
| 2 | 534 | [S, F] | [[493, 878], [2, 18]] | 1.688218e-02 |
| 3 | 355 | [R, K] | [[489, 865], [4, 30]] | 3.067271e-03 |
| 4 | 521 | [T, I] | [[495, 890], [0, 6]] | 9.484440e-02 |

#### Copy files to paper figures directory¶

In [21]:

```
paperdir = './paper'
figuresdir = os.path.join(paperdir, 'figures/')
myfiguresdir = os.path.join(figuresdir, 'Fig6/')
if not os.path.isdir(myfiguresdir):
    os.mkdir(myfiguresdir)

files = !ls results/H7N9/*
for f in files:
    shutil.copy(f, myfiguresdir)
```

In [22]:

```
files
```

Out[22]:

```
['results/H7N9/baltictreelong_div_355.pdf',
 'results/H7N9/baltictreelong_div_627.pdf',
 'results/H7N9/baltictreelong_div_D701N.pdf',
 'results/H7N9/baltictreelong_div_G685R.pdf',
 'results/H7N9/baltictreelong_div_R355K.pdf',
 'results/H7N9/baltictreelong_div_S534F.pdf',
 'results/H7N9/baltictreelong_div_T521I.pdf',
 'results/H7N9/baltictreelong_num_date_355.pdf',
 'results/H7N9/baltictreelong_num_date_627.pdf',
 'results/H7N9/baltictreelong_num_date_D701N.pdf',
 'results/H7N9/baltictreelong_num_date_G685R.pdf',
 'results/H7N9/baltictreelong_num_date_R355K.pdf',
 'results/H7N9/baltictreelong_num_date_S534F.pdf',
 'results/H7N9/baltictreelong_num_date_T521I.pdf',
 'results/H7N9/h7n9_pb2_snp_counts.txt',
 'results/H7N9/pairedhist.pdf']
```
