## Supplementary material for "Comprehensive mapping of avian influenza polymerase adaptation to the human host": File S2: Fig7_Accessibility.html


### Accessibility of adaptive mutations from current PB2 sequences¶

Are top adaptive mutations as identified in A549 human cells likely to be accessible by single nucleotide mutations from current avian influenza PB2 sequences? I determined accessibility (number of nucleotide substitutions required) of mutations from avian influenza PB2 sequences collected from 2015 to 2018.

###### Import modules, define directories, define functions¶

In [1]:

```
import os
import shutil
import string
import warnings
warnings.simplefilter('ignore') # don't print warnings to avoid clutter in notebook

import numpy as np
from scipy import stats
import pandas as pd
import matplotlib as mpl
import matplotlib.pyplot as plt

from Bio import SeqIO, AlignIO
from Bio.Data import CodonTable
from Bio.Seq import Seq

import dms_tools2
from dms_tools2.utils import codonEvolAccessibility

from IPython.display import display, HTML, Markdown, Image

# Directories
resultsdir = './results/' 
accessibilitydir = os.path.join(resultsdir, 'accessibility/')
if not os.path.isdir(accessibilitydir):
    os.mkdir(accessibilitydir)
```

In [2]:

```
# Set plotting parameters
import seaborn as sns
sns.set(context='paper', style='ticks', palette='deep', font='Arial', font_scale=1.042, color_codes=True,
        rc = {'font.size': 10,
 'axes.labelsize': 10,
 'axes.titlesize': 10,
 'xtick.labelsize': 9,
 'ytick.labelsize': 9,
 'legend.fontsize': 9,
 'axes.linewidth': 1.0,
 'grid.linewidth': 0.8,
 'lines.linewidth': 1,
 'lines.markersize': 4.5,
 'patch.linewidth': 0.8,
 'xtick.major.width': 1.0,
 'ytick.major.width': 1.0,
 'xtick.minor.width': 0.8,
 'ytick.minor.width': 0.8,
 'xtick.major.size': 4.5,
 'ytick.major.size': 4.5,
 'xtick.minor.size': 3,
 'ytick.minor.size': 3}
       )
# sns.plotting_context()
# Set colors to use throughout
palette = {"None":"gray",
          'All mutations': 'gray', 
          'Known human adaptive': "#d95f02", 
          'Top adaptive in A549': "#d95f02"
         }
```

In [3]:

```
# Gather annotations of mutations (known, novel)
dmssummarydf = pd.read_csv('results/diffsel/summary_prefs_effects_diffsel.csv')
tempdf = (dmssummarydf
#           .dropna(subset=['mutdiffsel'])
          [['site', 'wildtype', 'mutation','Known human adaptive', 'Experimentally adaptive in']]
          .sort_values('site')
         )
knowndf = (tempdf[tempdf['Known human adaptive']=='Yes']
           .assign(Type='Known human adaptive')
           .drop(columns=['Known human adaptive', 'Experimentally adaptive in'])
          )
exptdf = (tempdf[tempdf['Experimentally adaptive in']=='A549']
           .assign(Type='Top adaptive in A549')
           .drop(columns=['Known human adaptive', 'Experimentally adaptive in'])
          )
alldf = (tempdf.assign(Type='All mutations')
         .drop(columns=['Known human adaptive', 'Experimentally adaptive in'])
        )
mutannotdf = pd.concat([knowndf, exptdf, alldf])
mutannotdf.head()
```

Out[3]:

|  | site | wildtype | mutation | Type |
| --- | --- | --- | --- | --- |
| 8357 | 9 | D | N | Known human adaptive |
| 11458 | 74 | G | S | Known human adaptive |
| 3952 | 158 | E | G | Known human adaptive |
| 10814 | 189 | K | R | Known human adaptive |
| 6263 | 192 | E | K | Known human adaptive |

In [5]:

```
# Functions to annotate mutations with accessibility

def accessibility_threshold(row):
    if row['min Subst']==0:
        return '0'
    elif row['min Subst']<=1.1: # Picked this cut off to accommodate values slightly >1 when considering many sequences
        return '1'
    else:
        return '>1'
def seqsToCodonAcc(filename, seqlen):
    allowed_chars = set('ATCG')
    seqs = []
    for seq_record in SeqIO.parse(filename, 'fasta'):
        if len(seq_record.seq)==seqlen:
            if set(seq_record.seq).issubset(allowed_chars):
                seqs.append(str(seq_record.seq))
    #         else:
    #             print('Invalid bases:', seq_record.id, len(seq_record.seq))
        else:
            print('Not full length:', seq_record.id, len(seq_record.seq))
    accessibilitydf = (codonEvolAccessibility(seqs)
                     .melt(id_vars='site', value_vars=dms_tools2.AAS_WITHSTOP, 
                           var_name='toAA', value_name='min Subst')
                       .sort_values('site')
                    )
    accessibilitydf['accessibility'] = accessibilitydf.apply(lambda row: accessibility_threshold(row), axis=1)
    return accessibilitydf

accessibilityAvianPB2 = seqsToCodonAcc('data/Avian-PB2_2015-2018.fa', 2280)
accessibilityAvianPB2.head()
```

```
Not full length: cds:ASU48438 2277
Not full length: cds:ATY73852 2277
Not full length: cds:BBC21031 2277
```

Out[5]:

|  | site | toAA | min Subst | accessibility |
| --- | --- | --- | --- | --- |
| 0 | 1 | A | 2.0 | >1 |
| 2280 | 1 | E | 2.0 | >1 |
| 7600 | 1 | M | 0.0 | 0 |
| 8360 | 1 | N | 2.0 | >1 |
| 760 | 1 | C | 3.0 | >1 |

In [6]:

```
accessibilitydf = pd.merge(mutannotdf, accessibilityAvianPB2, how='left',
                          left_on=['site', 'mutation'], right_on=['site', 'toAA'])
accessibilitydf.head()
```

Out[6]:

|  | site | wildtype | mutation | Type | toAA | min Subst | accessibility |
| --- | --- | --- | --- | --- | --- | --- | --- |
| 0 | 9 | D | N | Known human adaptive | N | 0.998743 | 1 |
| 1 | 74 | G | S | Known human adaptive | S | 1.555625 | >1 |
| 2 | 158 | E | G | Known human adaptive | G | 0.999371 | 1 |
| 3 | 189 | K | R | Known human adaptive | R | 1.000000 | 1 |
| 4 | 192 | E | K | Known human adaptive | K | 1.000000 | 1 |

In [7]:

```
# Removing 'All mutations' from dataframe for swarmplot 
# (too many 'All mutations' for swarmplot to handle)
accessibilitydf2 = accessibilitydf[accessibilitydf['Type']!='All mutations']
```

In [8]:

```
fig, ax = plt.subplots(1, 1, figsize=(5,3.5))
ax = sns.violinplot(x='Type', y="min Subst", data=accessibilitydf, inner=None,
                   order=['All mutations', 'Known human adaptive', 'Top adaptive in A549'],
                   palette=palette, ax=ax)
ax = sns.swarmplot(x='Type', y="min Subst", data=accessibilitydf2, 
                   order=['All mutations', 'Known human adaptive', 'Top adaptive in A549'],
                    color="black", ax=ax)
ax.set_ylabel('Mean\nnucleotide\nsubstitutions\nfrom recent\navian\nsequences', 
             rotation='horizontal', va='center', labelpad=35)
ax.set_xlabel('')
ax.set(yticks=[0, 1, 2, 3])
sns.despine()
fig.savefig('results/accessibility/accessibility.pdf', bbox_inches='tight', dpi=300)
```

What are the known human adaptive mutations that require >1 nucleotide substitutions? I will check if what original sequences these mutations arose on.

In [9]:

```
accessibilitydf[(accessibilitydf['Type']=='Known human adaptive') &
                (accessibilitydf['accessibility']=='>1')
               ]
```

Out[9]:

|  | site | wildtype | mutation | Type | toAA | min Subst | accessibility |
| --- | --- | --- | --- | --- | --- | --- | --- |
| 1 | 74 | G | S | Known human adaptive | S | 1.555625 | >1 |
| 12 | 534 | S | F | Known human adaptive | F | 1.658077 | >1 |
| 14 | 588 | A | I | Known human adaptive | I | 1.817725 | >1 |
| 17 | 636 | L | F | Known human adaptive | F | 1.875550 | >1 |

What are the top new human adaptive mutations that require only 1 nucleotide substitution? It appears that 3 of 5 have already occurred in nature! : R355G, T521I, D701N.

In [10]:

```
accessibilitydf[(accessibilitydf['Type']=='Top adaptive in A549') &
                (accessibilitydf['accessibility']=='1')
               ]
```

Out[10]:

|  | site | wildtype | mutation | Type | toAA | min Subst | accessibility |
| --- | --- | --- | --- | --- | --- | --- | --- |
| 27 | 69 | E | K | Top adaptive in A549 | K | 1.000629 | 1 |
| 39 | 292 | I | T | Top adaptive in A549 | T | 1.063482 | 1 |
| 40 | 355 | R | G | Top adaptive in A549 | G | 1.030798 | 1 |
| 45 | 521 | T | I | Top adaptive in A549 | I | 1.032684 | 1 |
| 56 | 701 | D | N | Top adaptive in A549 | N | 0.998743 | 1 |

#### Copy files to paper figures directory¶

In [11]:

```
paperdir = './paper'
figuresdir = os.path.join(paperdir, 'figures/')
myfiguresdir = os.path.join(figuresdir, 'Fig7/')
if not os.path.isdir(myfiguresdir):
    os.mkdir(myfiguresdir)

files = !ls results/accessibility/*
for f in files:
    shutil.copy(f, myfiguresdir)
```

In [12]:

```
files
```

Out[12]:

```
['results/accessibility/accessibility.pdf']
```
