## Supplementary material for "Comprehensive mapping of avian influenza polymerase adaptation to the human host": File S2: FigS1_compareS009.html


### Phylogenetic relationship of A/Green-winged Teal/Ohio/175/1986 to other human and avian sequences¶

To check that the PB2 sequence of the avian influenza strain we chose to use in our experiments was representative of avian influenza strains in general, we examined the phylogenetic relationship of our chosen PB2 sequence to a sampling of other influenza strains. We are simply checking that it is not an outlier with respect to other influenza strains.

For influenza strains to compare to, we used strains sampled across different host species and years, as identified in Doud, Ashenberg, and Bloom (2015). We also included a set of lineage-defining strains (human H3N2 (Aichi 1968), human pandemic H1N1 (California/04/2009)), and representatives of recent sporadic human cases of avian influnza strains (H5N1, H7N9). These sequences are provided in file `compareS009/compareS009.fa`.

We first made a multiple alignment of these coding sequences with `mafft`, then inferred a phylogenetic tree using `raxml`.

In [2]:

```
import os
import shutil
import string

import pandas as pd
import numpy as np
import matplotlib as mpl
import matplotlib.pyplot as plt

from Bio.SeqRecord import SeqRecord
from Bio import SeqIO
from Bio import AlignIO

import seaborn as sns
# Set matplotlib rcParams for figures
sns.set(context='paper', style='ticks', palette='deep', font='Arial', font_scale=1.042, color_codes=True,
        rc = {'font.size': 10,
 'axes.labelsize': 10,
 'axes.titlesize': 10,
 'xtick.labelsize': 8, #9,
 'ytick.labelsize': 8, #9,
 'legend.fontsize': 9,
 'axes.linewidth': 1.0,
 'grid.linewidth': 0.8,
 'lines.linewidth': 1,
 'lines.markersize': 4.5,
 'patch.linewidth': 0.8,
 'xtick.major.width': 1.0,
 'ytick.major.width': 1.0,
 'xtick.minor.width': 0.8,
 'ytick.minor.width': 0.8,
 'xtick.major.size': 4.5,
 'ytick.major.size': 4.5,
 'xtick.minor.size': 3,
 'ytick.minor.size': 3}
       )
```

In [20]:

```
log = !mafft-linsi compareS009/compareS009.fa > compareS009/compareS009.fa.aln
```

In [3]:

```
pwd = !pwd
pwd = pwd[0]
raxmlout = os.path.join(pwd, 'compareS009/raxmlout')
if not os.path.isdir(raxmlout):
    os.mkdir(raxmlout)
log = !raxmlHPC -w {raxmlout} -n compareS009 -p 1 -m GTRCAT -s compareS009/compareS009.fa.aln
```

### Calculate amino-acid pairwise identity between PB2 sequences¶

It appears from the tree that the avian PB2 sequences are all quite similar. To confirm this, we calculated pairwise amino-acid identity of avian influenza PB2 sequences used to build the tree above. For comparison, we did the same for seasonal human influenza PB2 sequences.

In [2]:

```
# Modified from Biopython Tutorial and Cookbook
def make_protein_record(nuc_record):
    """Returns a new SeqRecord with the translated sequence (default table)."""
    return SeqRecord(seq = nuc_record.seq.translate(cds=True), 
                     id = nuc_record.id,
                    )

proteins = []
for nuc_rec in SeqIO.parse("compareS009/compareS009.fa.aln", "fasta"):
    if ("_HOST_Human_" in nuc_rec.id) or ("_HOST_Avian_" in nuc_rec.id):
        proteins.append(make_protein_record(nuc_rec))
SeqIO.write(proteins, "compareS009/compareS009_avian_human.faa", "fasta")
```

Out[2]:

```
58
```

In [6]:

```
def identity(seq1, seq2):
    j = 0 # counts positions in first sequence
    i = 0.0 # counts identity hits
    for amino_acid in seq2:
        if amino_acid == '-' or amino_acid == '*':
            pass
        else:
            if amino_acid == seq1[j]:
                i += 1
        j += 1
    seq = str(seq1)
    gap_strip = seq.replace('-','')
    gapstop_strip = gap_strip.replace('*','')
    percent = 100*i/len(gapstop_strip)
    return percent

def identityPairwise(infile):
    alignment = AlignIO.read(infile, "fasta")
    i = 0
    for n in range(len(alignment)):
        for m in range(1,len(alignment)-n):
            i += 1
    print(i)
    result = pd.DataFrame(np.empty([i,3]), columns=["seq1","seq2","Percent amino-acid identity"])
    i = 0
    for n, record in enumerate(alignment):
        seq1 = record.seq
        seq1name = record.id
        for m in range(1,len(alignment)-n):
            seq2 = alignment[n+m].seq
            seq2name = alignment[n+m].id
            ident = identity(seq1, seq2)
            result.loc[i] = [seq1name, seq2name, ident]
            i +=1
    return result
```

In [7]:

```
pi = identityPairwise('compareS009/compareS009_avian_human.faa')
pi.to_csv('compareS009/compareS009_avian_human_pairwiseid.txt', sep="\t", index = False)
```

```
1653
```

In [8]:

```
def annotate_pairwise(row):
    if ("HOST_Human_" in row['seq1']) and ("HOST_Human_" in row['seq2']):
        return "Human vs Human"
    elif ("HOST_Avian_" in row['seq1']) and ("HOST_Avian_" in row['seq2']):
        return "Avian vs Avian"
    else:
        return "Human vs Avian"
pi['comparison'] = pi.apply(lambda x:annotate_pairwise(x), axis=1)
pi.head()
```

Out[8]:

|  | seq1 | seq2 | Percent amino-acid identity | comparison |
| --- | --- | --- | --- | --- |
| 0 | A/green-winged\_teal/Ohio/175/1986\_H2N1\_HOST\_Av... | ref\_A/Albany/6/58\_H2N2\_HOST\_Human\_AY209936 | 96.179183 | Human vs Avian |
| 1 | A/green-winged\_teal/Ohio/175/1986\_H2N1\_HOST\_Av... | ref\_A/Panama/1/66\_H2N2\_HOST\_Human\_AY209945 | 96.179183 | Human vs Avian |
| 2 | A/green-winged\_teal/Ohio/175/1986\_H2N1\_HOST\_Av... | ref\_A/Tokyo/31/72\_H3N2\_HOST\_Human\_AY210150 | 95.520422 | Human vs Avian |
| 3 | A/green-winged\_teal/Ohio/175/1986\_H2N1\_HOST\_Av... | ref\_A/Memphis/1/1986\_H3N2\_HOST\_Human\_CY002759 | 95.388669 | Human vs Avian |
| 4 | A/green-winged\_teal/Ohio/175/1986\_H2N1\_HOST\_Av... | ref\_A/New\_York/433/2000\_H3N2\_HOST\_Human\_CY003255 | 95.125165 | Human vs Avian |

We plotted the distribution of amino acid identity for each set of comparisons.

In [21]:

```
g = sns.FacetGrid(pi, col="comparison",  
                  col_order=['Avian vs Avian', 'Human vs Human', 'Human vs Avian'],
                  sharey=False, 
                  height=2.5)
bins = np.linspace(92, 100, (100-92+1)*2)
g.map(plt.hist, "Percent amino-acid identity", color="steelblue", bins=bins)
g.savefig('compareS009/pairwiseid.pdf', bbox_inches="tight", dpi=300)
```

Get summary statistics for percent identity

In [10]:

```
groupby_comparison = pi['Percent amino-acid identity'].groupby(pi['comparison'])
summary = groupby_comparison.describe()
summary['meanAAdiff'] = (1-summary['mean']/100)*759
summary
```

Out[10]:

|  | count | mean | std | min | 25% | 50% | 75% | max | meanAAdiff |
| --- | --- | --- | --- | --- | --- | --- | --- | --- | --- |
| comparison |  |  |  |  |  |  |  |  |  |
| Avian vs Avian | 231.0 | 98.742935 | 0.725879 | 96.574440 | 98.287220 | 98.945982 | 99.341238 | 99.868248 | 9.541126 |
| Human vs Avian | 792.0 | 95.476837 | 1.000080 | 92.885375 | 94.861660 | 95.388669 | 96.047431 | 98.814229 | 34.330808 |
| Human vs Human | 630.0 | 97.163561 | 1.339284 | 93.939394 | 96.080369 | 97.233202 | 98.287220 | 99.868248 | 21.528571 |

Check mean pairwise identity specifically for our chosen avian influenza strain. We see that it has a slightly mean pairwise identity of 99%, slightly higher than all avian-avian comparisons. As such, there is no indication that it is a outlier.

In [17]:

```
pi_avian = pi[(pi['comparison']=='Avian vs Avian') & 
              (pi['seq1']=='A/green-winged_teal/Ohio/175/1986_H2N1_HOST_Avian_CY018884')]
pi_avian['Percent amino-acid identity'].describe().to_frame()
```

Out[17]:

|  | Percent amino-acid identity |
| --- | --- |
| count | 21.000000 |
| mean | 99.008721 |
| std | 0.618842 |
| min | 97.364954 |
| 25% | 98.814229 |
| 50% | 99.209486 |
| 75% | 99.341238 |
| max | 99.736495 |

#### Copy files to paper figures directory¶

In [22]:

```
paperdir = './paper'
figuresdir = os.path.join(paperdir, 'figures/')
myfiguresdir = os.path.join(figuresdir, 'FigS1/')
if not os.path.isdir(myfiguresdir):
    os.mkdir(myfiguresdir)

files = (
         ['compareS009/pairwiseid.pdf',
          'compareS009/raxmlout/RAxML_bestTree.compareS009']
        )
for f in files:
    shutil.copy(f, myfiguresdir)
```
